# Unraveling anti-inflammatory metabolic signatures of *Glycyrrhiza uralensis* and isoliquiritigenin through multiomics

**DOI:** 10.1101/2025.04.08.647718

**Authors:** Saki Kiuchi, Mi Hwa Chung, Taiki Nakaya, Katsuya Ohbuchi, Kazuya Tsumagari, Koshi Imami, Yasuhiro Otoguro, Tomoaki Nitta, Hiroyuki Yamamoto, Kazunori Sasaki, Hiroshi Tsugawa

## Abstract

*Glycyrrhiza uralensis*, known for its diverse pharmacological effects, including immunoregulation, anti-tumor, and antioxidant properties, is a widely used medicinal plant found in more than 70% of traditional herbal medicines (Kampo) in Japan. Although over 300 compounds have been discovered in *G. uralensis*, the molecular mechanisms underlying its bioactivity remain largely unknown due to the chemical diversity of its compounds. Here, we performed a multiomics analysis incorporating untargeted hydrophilic metabolomics, lipidomics, and phosphoproteomics to elucidate the molecular mechanisms distinguishing the effects of a single bioactive compound, isoliquiritigenin (ILG), and the extract of *G. uralensis* (GU). Multiomics time-course data were obtained for lipopolysaccharide (LPS)-stimulated RAW264.7 cells under four experimental conditions: control, LPS(+), LPS(+)/ILG(+), and LPS(+)/GU(+), where 182 hydrophilic metabolites, 381 lipids, and 13,211 phosphopeptides were characterized. The metabolic signatures of inflammatory macrophages, including increased levels of glycolytic intermediates, succinate, citrulline, triacylglycerols, and cholesteryl esters, were attenuated in both the GU(+) and ILG(+) groups. Using a multivariate approach based on a partial least squares algorithm with an imposed inflammation level order information, we identified upregulated phosphorylation of sirtuin 1 and 2 (SIRT1/2) along with alterations in nicotinamide adenine dinucleotide metabolism in the ILG(+) group. The inhibition of SIRT2 suppressed the anti-inflammatory effect of ILG, as indicated by a reduction in interleukin-6 (IL-6) levels. Furthermore, we discovered a substantial increase in γ-aminobutyric acid (GABA) and its downstream metabolite, 4-guanidinobutyric acid, in the GU(+) group. These increases were attributed to endogenous GABA production through glutamic acid decarboxylase rather than uptake via GABA transporters. Exogenous GABA administration significantly suppressed IL-6 and IL-1β expression in LPS-stimulated cells, and the simultaneous administration of GABA and ILG enhanced the anti-inflammatory effects. Consequently, this study presents an approach to elucidating the importance of traditional herbal formulations and demonstrates the utility of multiomics in uncovering that endogenous GABA production would facilitate anti-inflammatory effects with ILG in GU administration.

## Introduction

Herbal medicines have been used for centuries, either alone or in combination, primarily based on empirical knowledge rather than a comprehensive understanding of their mechanisms of action and safety profiles^1,2^. Unlike conventional pharmaceuticals that typically contain a single active ingredient, herbal medicines consist of complex mixtures of plant-derived metabolites, including specialized (secondary) metabolites, such as alkaloids, flavonoids, and saponins. These compounds exhibit diverse pharmacological activities, including antibacterial, antiviral, antioxidant, anti-inflammatory, and anticancer effects^3,4,5^. Although many bioactive compounds produce similar therapeutic outcomes, their underlying molecular mechanisms can vary significantly^6^. A prevailing hypothesis is that the therapeutic efficacy of herbal formulations arises not only from the additive effects of individual bioactive constituents but also from intricate interactions among diverse components, including those that may not exhibit bioactivity on their own. Multiple studies have demonstrated that herbal extracts or multi-herb formulations often exhibit superior efficacy compared to isolated active compounds or single-herb preparations administered at equivalent doses, highlighting the importance of synergistic interactions in herbal medicine^1,2^. Thus, a deeper understanding of these interactions would improve our knowledge of the safety and efficacy of herbal therapies as well as facilitate the development of optimized treatment strategies based on scientific evidence.

*Glycyrrhiza uralensis* is widely used in pharmacotherapy due to its diverse pharmacological properties, including anti-inflammatory, antioxidant, and antiviral effects^7, 8^. While *G. uralensis* and *G. glabra* are used as licorice worldwide, *G. uralensis* is incorporated into more than 70% of traditional herbal medicines (Kampo) in Japan^7, 8^. Over 300 natural products have been characterized from *G. uralensis* through metabolomics studies, with major components including glycyrrhizin, liquiritin, isoliquiritin, and their derivatives^7, 9^. These bioactive compounds exert anti-inflammatory effects by modulating key signaling pathways such as nuclear factor-kappa B (NF-κB), mitogen-activated protein kinase (MAPK), and high mobility group box-1, thereby attenuating inflammatory responses^10,11,12^. Clinically, licorice and its active constituents, such as glycyrrhizin and isoliquiritigenin, have been used in the treatment of inflammatory conditions, including acute lung injury, pharyngitis, bronchitis, and allergic disorders^10,12,13,14^. Moreover, licorice modulates the effects of herbal and Western medicines, as demonstrated in its application for drug addiction treatment and the alleviation of medication side effects^8,15^. These findings suggest that the anti-inflammatory properties of licorice extend beyond localized physiological effects and may represent a fundamental mechanism contributing to broader disease modulation. However, conventional approaches that focus on a limited number of molecules remain insufficient to fully elucidate the pharmacological mechanisms underlying the diverse bioactive constituents of herbal medicines.

Multiomics analysis is a powerful approach for hypothesis generation by integrating multiple omics layers, including genomics, transcriptomics, proteomics, and metabolomics. Not only does it allow for the identification of correlations, such as those between proteins and metabolites based on co-expression patterns, but it also facilitates the inference of causal relationships through integrative analysis of biological databases and literature mining. Therefore, in this study, we used a multiomics approach to provide scientific evidence supporting the importance of herbal medicine consumption. Specifically, we aimed to compare metabolic changes in response to two different treatment conditions: the administration of isoliquiritigenin (ILG), a well-characterized anti-inflammatory component of licorice, and the administration of the *G. uralensis* extract (GU), which contains a diverse array of bioactive compounds. To achieve this, we conducted metabolomics, lipidomics, and phosphoproteomics analyses. Among post-translational modifications, phosphorylation is the most prevalent and plays a critical role in regulating signal transduction^16^. Meanwhile, metabolomics profiling captures the dynamic biochemical outcomes of cellular processes, offering valuable insights into functional metabolic states. Therefore, integrating these multiomics datasets can reveal essential regulatory elements that drive the bioactivity of herbal medicines, thereby advancing our understanding of their mechanisms of action.

One limitation of this study is the potential inconsistency in the time points at which omics data were collected. Moreover, although it would be ideal to obtain shotgun proteomics and transcriptomics data simultaneously, these were not included in our current dataset. We acknowledge that monitoring molecular dynamics following drug administration is most effective when all omics data are collected at precisely matched time points and that incorporating a broader range of omics layers would enhance the accuracy and effectiveness of data mining. However, we believe that such limitations can partially be compensated for through mathematical modeling or multivariate analysis. To address this, we integrated three omics datasets with relevant biological databases and a variety of multivariate analytical techniques to delineate component-dependent metabolic signatures. Furthermore, we aimed to demonstrate the significance of multi-component administration by formulating new hypotheses based on the metabolic signatures and validating them through inhibition assays and targeted component administration experiments.

## Methods

### Cell culture and treatment for acquiring time-resolved multiomics data

RAW264.7 cells were grown at 37 °C in Dulbecco’s modified Eagle medium (DMEM) supplemented with 10% EquaFETAL (Atlas Biologicals, Inc.) in a humidified 5% CO2 atmosphere. For multiomics sample preparation, the cells were incubated with 30 μM ILG (Tokyo Chemical Industry Co., Ltd.) or 500 μg/mL GU (Tsumura & Co) for 2 h before stimulation with 100 ng/mL lipopolysaccharide (from *E. coli* O55:B5, Sigma Aldrich). ILG and GU were dissolved in dimethyl sulfoxide (DMSO) and DMEM, respectively. All samples, including vehicle controls, were prepared in DMEM with a final DMSO concentration of 0.1% to ensure consistency across experimental conditions. At several time points up to 24 h after LPS stimulation, cell collection and metabolome and proteome extraction were performed according to the described protocols in the following sections.

### Hydrophilic metabolome analysis

RAW264.7 cells incubated in six-well plates (Thermo Fisher Scientific, Nunclon Delta) were collected at 1 and 24 h after LPS stimulation. The medium was removed and washed twice with 5% mannitol. After the addition of 800 μL of methanol, the solvent was incubated for 30 s, followed by the addition of 550 μL of Human Metabolome Technologies internal standard solution 1 (Human Metabolome Technologies, Inc., Tsuruoka, Japan), with an additional 30 s incubation period. An aliquot of 1000 μL was transferred to a 1.5 mL tube and centrifuged at 4 °C and 2,300 g for 5 min. The supernatant (350 μL) was subjected to ultrafiltration to remove macromolecules. Thereafter, the filtrate was dried *in vacuo* and reconstituted in 50 μL of Milli-Q water for capillary electrophoresis coupled with mass spectrometry (CE/MS) analysis.

CE/MS experiments were performed using a CE System 7100 (Agilent Technologies Inc., Santa Clara, CA, USA), an Agilent 1260 isocratic HPLC pump, and a Q Exactive Plus (Thermo Fisher Scientific Inc., Waltham, MA) equipped with an electrospray ionization (ESI) adapter, following a previous study^17^. A variable-data-independent MS/MS acquisition (DIA) was utilized for comprehensive metabolome profiling ^18^. Metabolites were separated using a fused silica capillary (50 μm i.d. × 80 cm total length; Polymicro Technologies, Inc., Phoenix, AZ) with 50 mM ammonium acetate (pH 8.5) for anion analysis and 1 M formic acid for cation analysis with 10 μL/min of the sheath-flow rate. For the mass spectrometer setting, the full scan ranges for cation and anion analyses were set to *m/z* 60–500 and *m/z* 70–800, respectively.

For data processing, peak picking, chromatogram deconvolution, annotation, migration time (MT) correction, and peak alignment of the CE-MS/MS dataset were performed using MS-DIAL 5^19^. Using the retention time correction function, migration time was linearly corrected with the migration time values of the internal standards, and alignment tolerance was set to 0.5 min. The in-house MT library was used to predict the MT values of compounds in the MSMS spectral library and applied to metabolite annotation. Details of the metabolite annotation method using comprehensive MS/MS are as described in our previous study^18^.

### Lipidome analysis

RAW264.7 cells incubated in six-well plates (Thermo Fisher Scientific) were collected at 0, 0.25, 1, 6, and 24 h after LPS stimulation. Lipidome extraction was performed using the Bligh & Dyer method^20^. Briefly, the cells were washed twice with phosphate buffered saline (PBS) and quenched with ice-cold methanol. The cells, suspended in 500 µL methanol, were vortexed with 200 µL each of water (Fujifilm wako) and chloroform (Fujifilm wako). The mixture was sonicated for 5 min (30 s × 10 cycles) using a Bioruptor 2 (Sonicbio) and then shaken for 10 min at 1,200 rcf using a shaker (Thermo Fisher Scientific). Following centrifugation at 4 °C, 16,000 g for 3 min, 700 µL of the supernatant was collected and vortexed with an additional 280 µL of water. After phase separation by centrifugation at 4 °C, 16,000 g for 3 min, 250 µL of the liposoluble fraction from the lower phase was collected. The liposoluble fraction was dried using a centrifugal evaporator CC-105 (TOMY Digital Biology) and redissolved in 100 µL methanol containing 1 µM fatty acid (FA) 16:0 (d3), 1 µM FA 18:0 (d3), and 1% EquiSPLUSH as internal standards. The solution was centrifuged at 4 °C, 16,000 g for 3 min, and the resulting supernatant was transferred to a glass amber vial with a micro insert (Agilent Technologies). A 1 μL aliquot was injected.

Liquid chromatography coupled with tandem mass spectrometry (LC-MS/MS) measurements were performed using ExionLC AD (SCIEX) and Zeno TOF 7600 system (SCIEX). Separation by reversed-phase liquid chromatography was performed on YMC-Triart C18 (50 × 2.0 mm, 1.9 µm, YMC, Japan) at 40 °C. The mobile phases were (A) acetonitrile (ACN)/methanol/water (1/1/3, v/v/v) with 0.5% ammonium acetate and 0.02% ethylenediaminetetraacetic acid (EDTA), and (B) ACN/2-propanol (1/9, v/v) with 0.5% ammonium acetate and 0.02% EDTA. The flow rate was set to 300 μL/min. The gradient condition was (B) 0.5−40% for 4 min, (B) 40−64% for 2.5 min, (B) 64−71% for 4.5 min, (B) 71−82.5% for 0.5 min, (B) 82.5−85% for 6.5 min, (B) 85−99% for 1 min, (B) 99% for 2 min, (B) 99%−0.5% for 0.1 min, and (B) 0.5% for 4.4 min. A data-dependent MS/MS acquisition mode was used. The MS conditions are as follows: MS1 scan range, *m/z* 70–1250; MS/MS scan range, *m/z* 50-1250; MS1 accumulation time, 200 ms; MS2 accumulation time, 50 ms; Q1 resolution, units; maximum candidate ions, 10; ion source temperature, 250 °C; and CAD gas, 7. The following settings were used for positive/negative ion mode, independently: ion source gas 1, 40/50 psi; ion source gas 2, 80/50 psi; curtain gas, 30/35 psi; spray voltage, 5500/−4500 V; de-clustering potential, 80/−80 V; and collision energy, 40/−42 ± 15 eV. Mass calibration was automatically performed using a SCIEX calibration delivery system.

The data processing was performed using MS-DIAL version 5.1.230719 with the following parameters: minimum amplitude, 1,000; retention time (RT) tolerance for identification using the txt library, 0.2 min; RT tolerance for peak alignment, 0.1 min. Data normalization was performed using internal standards according to the previous report^21^.

### Phosphoproteome analysis

RAW264.7 cells incubated in 10 cm plates (Thermo Fisher Scientific) were collected at 0, 0.25, 2, and 24 h after LPS stimulation. Total cellular protein was extracted using the phase transfer surfactant (PTS) method^22^. Pellets of RAW264.7 cells subjected to a series of pretreatments were dissolved in 750 µL of PTS solution containing 12 mM sodium deoxycholate (Fujifilm Wako), 12 mM sodium lauryl sulfate (Fujifilm Wako), and 100 mM Tris-HCl (pH 9.0, Fujifilm Wako), following sonication for 5 min and shaking for 10 min. The solution was heated at 95°C for 5 min, after which the protein concentration was measured using the bicinchoninic acid (BCA) assay. One hundred micrograms of protein was treated with (±)-dithiothreitol (Fujifilm Wako) for 30 min at RT (25°C). Iodoacetamide (final conc., 50 mM; Fujifilm Wako) was added and incubated for 30 min at 25°C in the dark. Subsequently, the sample was diluted 5-fold with 50 mM ammonium bicarbonate (Fujifilm Wako), and 1 µg of trypsin (Promega) was added, followed by overnight incubation. To remove PTS, equal volumes of ethyl acetate and trifluoroacetic acid (final conc., 0.5%; Fujifilm Wako) were added to the sample. The mixture was centrifuged at 15,700 g for 2 min, and the upper organic layer was discarded. The aqueous layer was dried using a centrifuge evaporator and redissolved in 200 µL of buffer composed of ACN:water (5:95) containing 0.5% trifluoroacetic acid.

Phosphopeptide enrichment was performed using the hydroxy acid-modified metal oxide chromatography (HAMMOC) method^23,24^. Centrifugations during conditioning, sample application, and elution were performed at 1,000 g for 3 min at 4°C. An Empore Disk C8 (47 mm, GL Sciences) was picked by a Kel-F Hub needle (point style 3, 20 gauge, HAMILTON) and packed with a Plunger Assembly 1701 (HAMILTON). The tip was packed with 0.5 µg of titanium (IV) oxide (GL Sciences) and sequentially conditioned with 20 µL of buffer B containing ACN/water (5/95,v/v) with 0.1% trifluoroacetic acid (TFA) and 20 µL of buffer C containing 300 mg/mL DL-lactate (Fujifilm Wako) in buffer B. The samples diluted 2-fold with buffer C were loaded and centrifuged. Subsequently, the column was washed with 20 µL of buffer C and 50 µL of buffer B. Finally, phosphopeptides were eluted with 20 µL of 0.5% pyrrolidine (Fujifilm Wako). Further desalting was performed using a stop-and-go-extraction tip (StageTip)^25^. An Empore Disk SDB-XC (47 mm diameter, GL Science) was picked by a Kel-F Hub needle (point style 3, 16 gauge, HAMILTON) and packed with a Plunger Assembly 1702 (HAMILTON) into the tip. The column was conditioned sequentially with 20 µL of buffer B and 20 µL of buffer A containing ACN/water (80/20, v/v) with 0.1% TFA. The sample was applied, followed by a wash with 20 µL of buffer A. Phosphopeptides were eluted with 20 µL of buffer B. Eluted samples were dried using the centrifugal evaporator CC-105 (TOMY digital biology), redissolved in 10 µL of buffer A, and centrifuged at 16,000 g for 3 min at 4°C. The supernatant was then analyzed by nanoLC-MS/MS.

A nanoLC-MS/MS system comprising an EASY-nLC 1200 (Thermo Fisher Scientific) pump and an Orbitrap Eclipse mass spectrometer (Thermo Fisher Scientific) was used. The mobile phases consisted of (A) 0.1 % formic acid and (B) 0.1 % formic acid and 80 % ACN. Peptides were loaded on a self-made 20 cm fused-silica emitter (75 µm inner diameter) packed with ReproSil-Pur C18-AQ (1.9 µm, Dr. Maisch, Ammerbuch, Germany) and separated by a linear gradient (3−40% B in 105 min, 40−99% B in 5 min, and 99% B for 10 min) at a flow rate of 300 nL/min. All MS1 spectra were acquired over 375–1500 *m/z* in the Orbitrap analyzer (resolution = 120,000, maximum injection time = 50 ms, and automatic gain control = “Standard”). For the subsequent MS/MS analysis, precursor ions were selected and isolated in a top-speed mode (cycle time = 1 s and isolation window = 1.6 *m/z*), activated by higher-energy collisional dissociation (normalized collision energy = 28), and detected in the Orbitrap analyzer (resolution = 30,000, maximum injection time = 54 ms, and auto gain control = 100%). Dynamic exclusion time was set to 60 s.

NanoLC/MS/MS raw data were processed using the FragPipe (v.18.0) suite with MSFragger (v.3.5), Philosopher (v.4.2.1), and IonQuant (v.1.8.0)^26,27,28^. Database search was implemented against the UniProt mouse proteome database including isoform sequences (April 2022). Generally, default parameters were used. The precursor and fragment ion mass tolerances were set to 20 ppm with mass calibration and parameter optimization enabled. Strict trypsin cleavage specificity was applied, with up to two missed cleavages allowed. Cysteine carbamidomethylation was set as a fixed modification, while oxidation on methionine, acetylation on the protein *N*-terminus, and phosphorylation on serine, threonine, and tyrosine were allowed as variable modifications. The match-between-runs function was enabled. The search results were filtered for < 1% at the peptide-spectrum match, ion, peptide, and protein levels.

### Enzyme-linked immunosorbent assay (ELISA) for IL-6 production

Following one-hour incubation with different agents including GU, components from *G. uralensis*, EX527 (Cayman Chemical), and AGK-2 (TCI, Japan), LPS was added to the culture. After 24 h, culture supernatants were collected, and levels of interleukin (IL)-6 in the culture supernatants were determined using ELISA kits (R&D Systems) with four biological replicates per condition. The reagents were obtained from the following sources: glycyrrhizin (Fujifilm Wako), glycyrrhetinic acid (TCI, japan), EX527, and AGK-2.

### RNA isolation and reverse transcription-quantitative polymerase chain reaction (RT-qPCR)

RAW264.7 cells were incubated overnight in six-well plates and treated with different compounds as follows: For the evaluation of γ-aminobutyric acid (GABA) receptor activity, the cells were incubated for 1 h with muscimol (2 µM, Sigma-Aldrich), baclofen (50 µM, TCI, Japan), bicuculline (50 µM, TCI, Japan), and CGP 55845 (5 µM, Sigma-Aldrich). After incubation, GABA (1 µM, TCI, Japan) was added for 1 h, followed by LPS (100 ng/mL) stimulation for 2 h. For the evaluation of the anti-inflammatory activity of ILG and GABA, the cells were incubated with ILG (10 µM) and/or GABA for 1 h, followed by LPS (100 ng/mL) stimulation for 2 h.

Total RNA was isolated using ISOGEN II (Fujifilm Wako). cDNA was synthesized using iScript™ Reverse Transcription Supermix (Bio-Rad) following the manufacturer’s instructions. Quantitative real-time PCR analysis was performed with iTaq Universal SYBR Green Supermix (Bio-Rad) on the CFX Duet Real-Time PCR System (Bio-Rad). The primer sequences are shown in **Table S1**. Changes in the expression of target genes were calculated using the 2^-*ΔΔ*Cq^ method and normalized to the value of the housekeeping gene, *Gapdh*.

### Western blotting

After the cells were washed twice with Dulbecco’s Phosphate Buffered Saline (D-PBS) (-) (Fujifilm Wako), cellular proteins were solubilized and extracted using 1 mL of Radio-Immunoprecipitation Assay (RIPA) buffer (Fujifilm Wako). The lysates were boiled at 95 °C for 5 min to denature the proteins, and protein concentrations were determined using a BCA protein assay kit. Protein samples were separated based on molecular mass by SDS–PAGE and then transferred onto a cellulose nitrate membrane. The membranes were washed twice with distilled water and blocked with 50 mg/mL skim milk powder for 30 min at room temperature (25°C) and incubated with primary antibodies for 24 h. Subsequently, the membranes were washed thrice with Tris-buffered saline with 0.1% Tween-20 (TBST), followed by incubation with a secondary antibody for 1 h at 25°C. After the membrane was washed thrice with TBST, Clarity and Clarity Max ECL Western Blotting Substrate (Bio-Rad) was added to the membrane. The proteins were quantified using the FUSION-Chemiluminescence Imaging System (M&S Instruments). The following antibodies for immunoblotting were used: anti-GAPDH [14C10] (Abcam), anti-SIRT1 [19A7AB4] (Abcam), and anti-SIRT2 [EPR20411-105] (Abcam).

### Hydrophilic metabolome analysis of G. uralensis extract using LC-MS/MS

Metabolites in 10 mg of GU were extracted using the Bligh & Dyer method^20^. As described in the protocol for lipid extraction, 900 µL of the sample was dissolved in 900 µL of a methanol:water:chloroform (5:2:2, v/v/v) extraction solution. Following sonication and vortexing, 700 µL of the upper layer was collected by centrifugation at 16,000 g for 3 min at 4 °C. After liquid–liquid phase separation by adding cold water (280 μL), 600 µL of the upper phase, representing the hydrophilic fraction, was recovered and dried using a centrifugal evaporator. The dried sample was resuspended in 100 µL of 80% methanol, centrifuged at 16,000 g for 3 min, and the resulting supernatant was subjected to LC-MS/MS analysis.

LC-MS/MS analysis for GABA quantification was performed using the Zeno TOF 7600 system (SCIEX) with ExionLC AD (SCIEX). The reverse-phase LC separation was achieved by an ACQUITY UPLC BEH column (particle size, 1.7 μm, 100 mm by 2.1 mm i.d., Waters Corporation) at 30°C. The mobile phase was prepared by mixing solvents (A) 0.1% formic acid (v/v) in water and (B) methanol at a flow rate of 150 μl/min. The data-dependent acquisition mode was used under the following conditions: interface temperature, 450 °C; ESI spray voltage, 5500 V; polarity, positive; MS1 mass range, *m/z* 50–1500; MS2 mass range, *m/z* 50–1500; collision energy, 35 V; and collision energy spread, 15 V. The identification and quantification of GABA, ILG, and glycyrrhizin were performed based on the analysis of standard mixtures of GABA, ILG, and glycyrrhizin.

### GABA isotope tracing experiment

RAW264.7 cells were cultured in six-well dishes for 1 h with 1 μM GABA-d6 (COSMO BIO) in the presence of 500 μg/mL GU and stimulated with 100 ng/mL LPS. At 1 and 24 h after LPS stimulation, cells were washed twice with 1 mL PBS and quenched with 500 μL ice-cold MeOH. Hydrophilic metabolites were extracted using the Bligh & Dyer method as described in the previous section by adding 200 μL each of H2O and CHCl3 to the collected samples.

LC-MS/MS analysis was performed using 6546 LC/Q-TOF (Agilent) with 1260 infinity II (Agilent). The reverse-phase LC separation was achieved using an ACQUITY UPLC BEH column (particle size, 1.7 μm, 100 mm by 2.1 mm i.d., Waters Corporation) at 30°C. The mobile phase was prepared by mixing solvents (A) 0.1% formic acid (v/v) in water and (B) methanol at a flow rate of 150 μl/min. The data-dependent acquisition mode was used under the following conditions: interface temperature, 450 °C; ESI spray voltage, 5500 V; polarity, positive; MS1 mass range, *m/z* 50–1250; MS2 mass range, *m/z* 50–1250; collision energy, 15 V.

### GABA quantification in RAW264.7 cells with glutamate decarboxylase (GAD) inhibitor

The cells were incubated with 10 mM of allylglycine (TCI, Japan) for 2 h, and GU was added to achieve a final concentration of 500 μg/mL. After 2 h of incubation, cells were washed with PBS and collected in 500 μL ice-cold MeOH. Hydrophilic metabolites were extracted using the Bligh & Dyer method as described in the previous section. LC-MS/MS analysis was performed using ExionLC AD (SCIEX) and the Zeno TOF 7600 system (SCIEX) with MS1 and MS2 range at *m/z* 50-750 according to the method previously described in the metabolome analysis section of the *G*.*uralensis* extract.

### Multiomics data integration using multiset partial least squares with rank order of groups (PLS-ROG)

The multiset PLS-ROG was performed using the R package loadings (version 0.5.1)^29^. The explanatory variable matrix X was defined as multiomics data collected 24 hours after LPS stimulation, with X1 representing hydrophilic metabolome data, X2 phosphoproteome data, and X3 lipidome data. The response variable Y was set as a matrix representing group information, while a ranking matrix D was used to define the group order based on IL-6 levels as an inflammation marker. To minimize the influence of data outliers, the coupling strength parameter *τ* between groups and omics datasets was set to 0.499. Additionally, the smoothing parameter *κ* was set to 0.999 to ensure that group rankings were effectively reflected in the multiset PLS-ROG scores. Data matrix of X, Y, tau and D and R scripts for multiset PLS-ROG is shown in Source data. Pathway enrichment analysis using phosphoproteome and metabolome data was performed by MetaboAnalyst^30^.

## Results

### Multiomics profiling of LPS-inflamed RAW264.7 cells treated with G. uralensis components

We evaluated the anti-inflammatory effects of several *G. uralensis*-derived natural products, including glycyrrhizin (GL), its hydrolysis product glycyrrhetinic acid (GA), ILG, and GU, in RAW264.7 cells. We assessed IL-6 production 24 h after LPS stimulation. The results showed that GU and ILG significantly suppressed IL-6 production in a dose-dependent manner while GL and GA did not alter IL-6 production rates, although a previous study has shown the anti-inflammatory effect of GL in RAW264.7 cells^6^ (**Figures 1a and S1**). As the anti-inflammatory effects of GU and ILG in acute inflammatory conditions were confirmed in our experimental setup^6^, we selected these two agents for multiomics analysis. A time-series multiomics analysis was conducted on RAW264.7 cells treated with ILG (30 µM) and GU (500 µg/mL) under LPS-stimulated conditions (**Figure 1b**). RAW264.7 cells were pre-incubated with ILG or GU for 1 h, followed by stimulation with 100 ng/mL LPS inducing toll-like receptor 4-mediated inflammation. Samples were collected at different time points to capture temporal changes: 1 and 24 h for hydrophilic metabolomics, 0, 0.25, 1, 6, and 24 h for lipidomics, and 0, 0.25, 2, and 24 h for phosphoproteomics. The score plots of principal component analysis (PCA) using the auto-scaling method as data pre-processing revealed that the metabolome, lipidome, and phosphoproteome profiles substantially changed due to the treatments of ILG and GU (**Figure 1c**). We observed changes in the lipidome and phosphoproteome at 0 h, indicating that molecular profiles were affected solely by the administration of ILG and GU. In addition, slight changes in the phosphoproteome were observed at 0 h following LPS administration, where samples were collected immediately after stimulation. However, the LPS-treated group was not separated from the control group in PCA score plots based on the first and second principal components. These results suggest that omics profiles are rapidly influenced by various environmental factors. Nevertheless, our data indicate that single-agent ILG and multi-component GU elicit distinct metabolome and phosphoproteome signatures at the early stages of inflammation.

**Figure 1.**
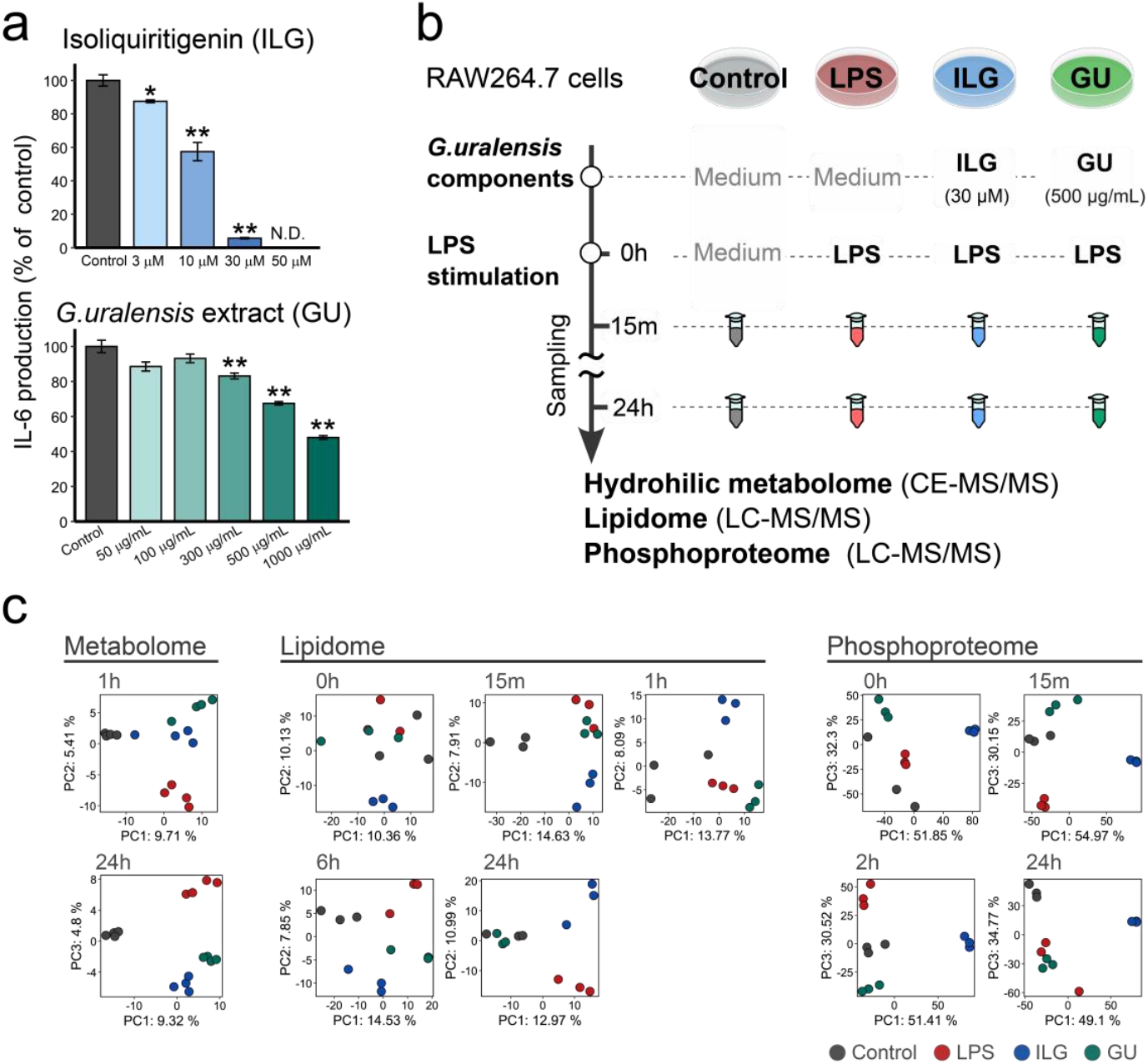
Overview of multiomics profiling in LPS-stimulated RAW264.7 cells treated with isoliquiritigenin and *Glycyrrhiza uralensis* extract. (a) IL-6 production in RAW264.7 cells incubated with ILG or *G. uralensis* extract. Following 18 h of stimulation with 100 ng/mL lipopolysaccharide (LPS), culture supernatants were collected, and IL-6 production was quantified using ELISA (*n*=3 for biological replicates). Each bar plot is the mean of relative IL-6 production compared to the LPS group with error bars representing the standard error of the mean (SEM). Statistical significance was determined using Tukey’s honestly significant difference (HSD) test comparing each treatment to the LPS group (two-sided). (b) Experimental design of multiomics analysis. (c) Score plots of principal component analysis using the auto-scaled table of hydrophilic metabolome (left), lipidome (middle), and phosphoproteome (right) data at each time point after LPS stimulation.

### Investigating ILG- and GU-specific metabolome alterations in RAW264.7 cells

Using capillary electrophoresis coupled with the data-independent MS/MS acquisition method^18^, we characterized 182 hydrophilic metabolites, comprising carbohydrates, amino acids, and nucleic acids, in RAW264.7 macrophages 1 and 24 h after LPS stimulation (**Figures 2a and S2a**). The heatmap and the number of significantly altered metabolites, based on the cell number-normalized metabolome table, indicated that most hydrophilic metabolites increased under LPS-stimulated conditions, with more pronounced shifts observed after 24 h in the LPS and ILG groups (**Figures 2a, 2b, S2a, and S2b**). In addition, several unique metabolic changes were observed in the GU-treated group at 1 h after LPS administration. While these metabolites can be attributed to components of GU, our results suggest that macrophages hydrolyze glycosylated plant metabolites to their aglycones. For example, isoliquiritin, a major component of GU, increased at 1 h, while its aglycone, ILG, increased at 24 h (**Figure 2a**). Although the degradation activity of glycosylated plant metabolites in immune cells is poorly understood, a previous study has also demonstrated similar enzymatic activity in RAW264.7 cells involving a different class of plant metabolites, such as genistein glycosides^31^. Moreover, GABA and its downstream metabolite, 4-guanidinobutyric acid, were enriched in the GU-treated group at 1 and 24 h after LPS stimulation, whereas their precursors such as glutamine and glutamic acid exhibited no significant alterations (**Figures 2a and 2c**). The production mechanism and anti-inflammatory effect of GABA are further investigated in a later section.

**Figure 2.**
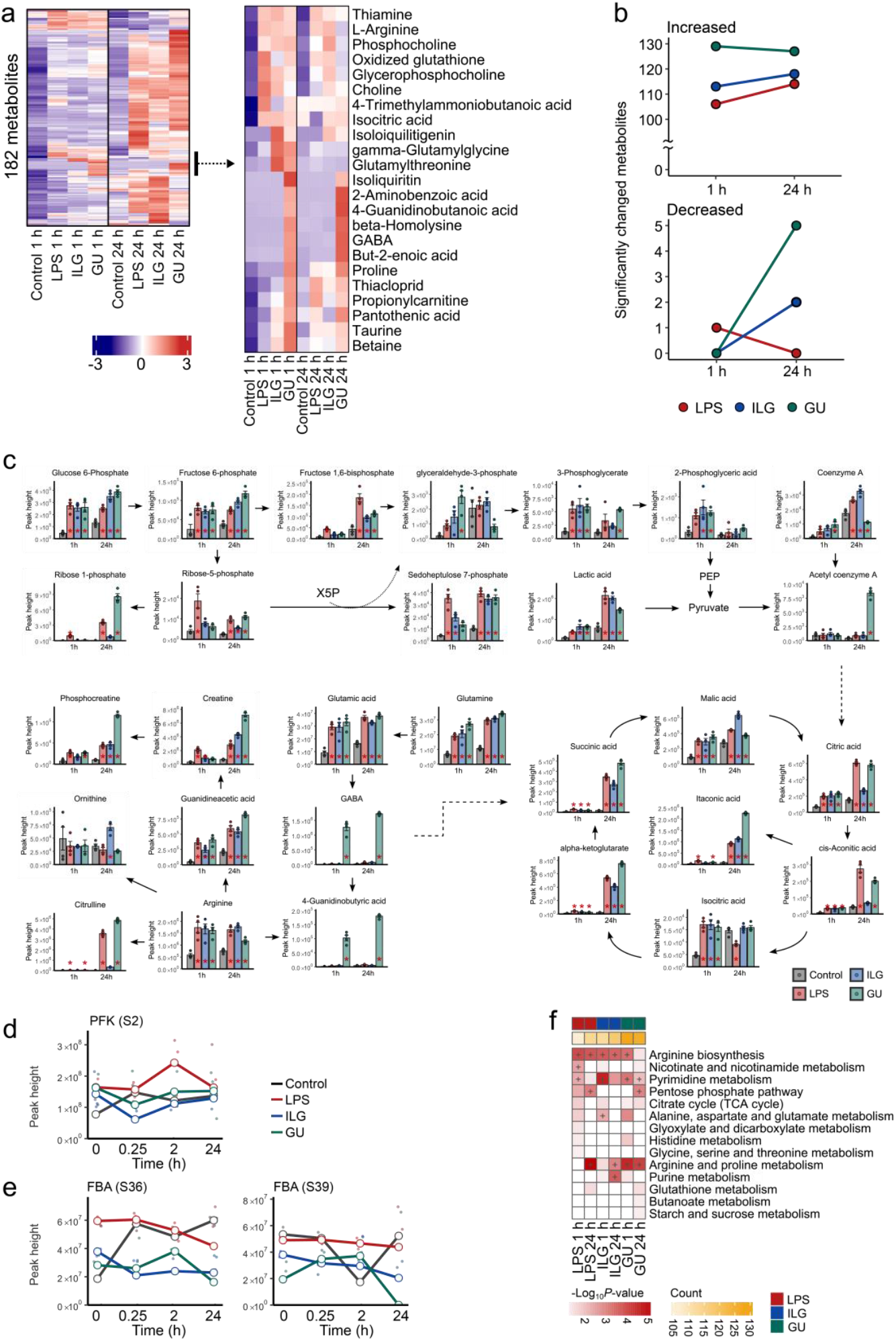
Hydrophilic metabolome alterations by ILG and GU treatments in LPS-stimulated RAW264.7 cells. (a) Heatmap showing ion abundance ratios among four groups. Peak heights were normalized by the number of cells at each sampling point, and Z-scaled data were used for clustering analysis. (b) Summary of significantly changed metabolites compared to control at each time point. *P*-values were calculated using the Tukey-Kramer method. The number of metabolites exhibiting more than a two-fold change with *p*-values < 0.05 is shown. (c) Mapping of metabolic profiles onto glycolysis, the TCA cycle, and the urea cycle after 1 and 24 h of LPS stimulation. Bar plots represent the mean values for each group, with individual data points overlaid as a scatter plot. Error bars indicate the standard error of the mean (SEM) and *p*-values were calculated using the Tukey’s HSD test (two-sided) and labeled if *p*-value < 0.05 compared to the control group at the same time point. (d) Peak heights of the phosphopeptide of phosphofructokinase (PFK) at each time point, where S2 indicates phosphorylation at second serine residue. (e) Peak height profiles of phosphorylated peptides at S36 and S39 of fructose 1,6 bisphosphate aldolase (FBA). (f) Results of metabolite set enrichment analysis using MetaboAnalyst version 5.0. The number of significantly changed metabolites is also shown with column annotations. Pathway terms that were significant within the top three in each group are indicated by “+” label.

We investigated unique metabolic signatures of ILG and GU in the context of biochemical pathways. At 1 h after LPS stimulation, intermediates of the glycolysis pathway such as 2-phosphoglyceric acid and 3-phosphoglyceric acid in addition to metabolites associated with phosphatidylcholine (PC) metabolism increased (**Figures 2c and S2c**). These results are consistent with previous studies indicating that LPS stimulation induces a shift toward anaerobic glycolysis and triggers membrane phospholipid remodeling in macrophages within minutes to hours^32,33^. Notably, the metabolic shifts were attenuated in the ILG- and GU-treated groups, suggesting that *G. uralensis*-derived metabolites suppress LPS-induced inflammation by modulating glycolysis and phospholipid metabolisms. The number of phosphorylations of key glycolytic enzymes, including phosphofructokinase (PFK) and fructose-1,6-bisphosphate aldolase (FBA) significantly decreased in the ILG and GU groups. This suggests that the suppression mechanism of GU-derived natural products on anaerobic glycolysis is exerted through the phosphorylation of these metabolic enzymes (**Figures 2d and 2e)**. Furthermore, the ontology terms related to the arginine metabolism and TCA cycle pathways were enriched (**Figure 2f**). Although this trend was observed across all treatment conditions, there were unique metabolic signatures in each group on the same metabolic pathway. The levels of citrulline and itaconate, both of which accumulate in LPS-stimulated macrophages, were markedly elevated at 1 h after stimulation, with further accumulation at 24 h^34,35,36^ . Meanwhile, the accumulation of citrulline was significantly suppressed by ILG treatment but was unaffected by GU treatment (**Figure 2c**).

### ILG and GU treatment-specific lipidome alterations in LPS-stimulated RAW264.7 cells

We conducted untargeted lipidomics using LC-MS/MS, profiling 381 lipids across 31 lipid classes (**Figure 3a**). Peak intensities were normalized using internal standards and cell numbers. The PCA score plots using the auto-scaled data showed that the lipidomes between 0 and 24 h after LPS stimulation were clearly distinguished in the first principal component (PC1) in all groups, i.e., the LPS(+), LPS(+)/ILG(+), and LPS(+)/GU(+) groups (**Figure 3b**). The number of lipids that significantly increased in number in the ILG and GU groups peaked 15 min after LPS stimulation and gradually declined over time (**Figure 3c**). Moreover, the number of significantly changed lipids increased again at 24 h in the LPS(+) and LPS(+)/ILG(+) groups, aligning with the observations in the hydrophilic metabolome. These results suggest that GU treatment induces early metabolome changes, while ILG treatment leads to unique alterations compared to the control at a later stage. The increases in metabolites and lipids in the LPS(+)/GU(+) group should partially be attributed to the presence of GU-derived plant metabolites in the metabolome and lipidome datasets.

**Figure 3.**
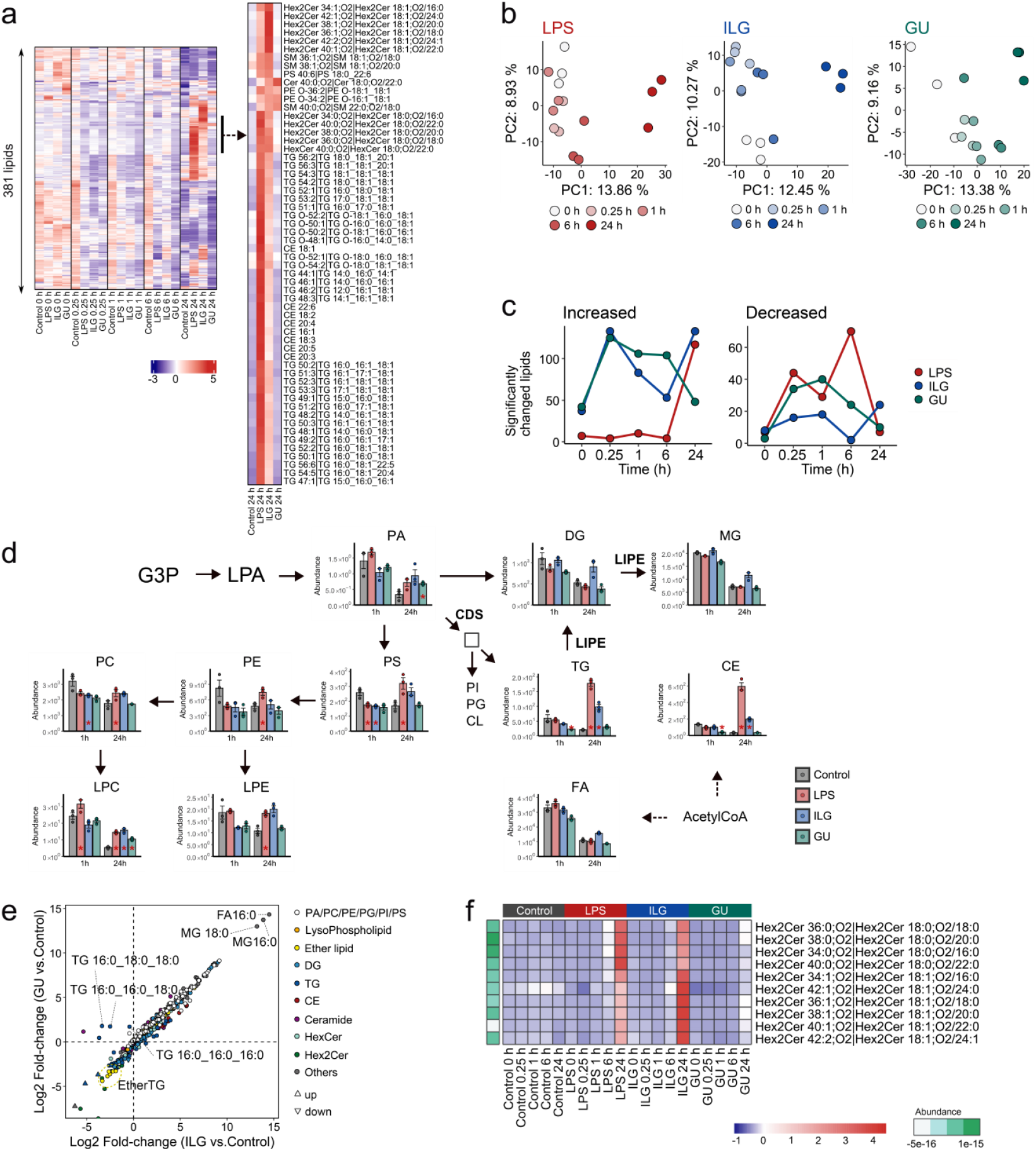
ILG- and GU-specific lipidome alterations in LPS-stimulated RAW264.7 cells. (a) Heatmap analysis of 381 lipid molecules, also highlighting clusters that increased in the ILG and LPS groups after 24 h. (b) PCA score plots illustrating time-resolved changes in the lipidome profiles of the LPS, ILG, and GU groups. (c) Line plots displaying the number of significantly changed lipids compared to the control at the same time point. Lipids with an adjusted *p*-value < 0.05 and absolute log2(fold-change) > 1 were defined as significantly changed. *P*-values were calculated using the Tukey’s HSD test (two-sided). (d) Mapping of lipidome data onto the de novo lipid biosynthesis pathway, showing lipid profiles at 1 and 24 h after LPS stimulation. Bar plots represent the mean values for each group, with individual data points overlaid as a scatter plot. Error bars indicate the SEM. *P*-values were calculated using the Tukey’s HSD test (two-sided) and annotated if p-value < 0.05 compared to the control group at the corresponding time point. (e) Relationship of log2 fold changes in the ILG and GU groups compared to the control 24 h after LPS stimulation. (f) Heatmap showing Hex2Cer lipid abundances following 24 hours of LPS stimulation, with log2-transformed value of summed peak heights among samples also displayed.

Triacylglycerol (TG) and cholesteryl ester (CE), the primary constituents of lipid droplets, along with phosphatidylserine, a key regulator of eat-me signaling in macrophages, exhibited significantly elevated levels in the LPS group, particularly after 24 h, but were suppressed in the ILG and GU groups (**Figures 3a and 3d**). Moreover, the phosphorylation levels of phosphatidate cytidylyltransferase 2, which catalyzes the conversion of phosphatidic acid to cytidine diphosphate-diacylglycerol, also increased in the LPS group and decreased due to ILG and GU treatment (**Figure S3a**). The increases in TG and CE in LPS-stimulated macrophages are consistent with previous reports^37,38,39^. In addition, dihexosylceramide (Hex2Cer) levels increased progressively in LPS-stimulated macrophages (**Figures 3d and S3b**). In the ILG group, diacylglycerol (DG) and monoacylglycerol (MG) levels were elevated, alongside increased phosphorylation of hormone-sensitive lipase (LIPE) at Ser602, a site that inhibits LIPE translocation to lipid droplets when phosphorylated by AMP-activated protein kinase (AMPK) **(Figure S3c)**^40^. Furthermore, in terms of the characteristics of each lipid molecule, the addition of GU increased the number of TG molecules consisting of saturated fatty acids, while overall TG levels tended to decrease (**Figure 3f**). In addition, LPS- and ILG treatment groups facilitated the increase of Hex2Cer molecules, where the sphingosine (18:1;O2) containing Hex2Cer molecules were increased in ILG group while those containing sphinganine (18:0;O2) were higher in LPS group (**Figures 3f and S3b**).

### ILG- and GU-specific phosphoproteome alterations in LPS-stimulated RAW264.7 cells

A total of 13,211 phosphopeptides were characterized in ILG- and GU-treated RAW264.7 macrophages. The peptides detected in fewer than two of the three biological replicates were excluded, resulting in 12,021 peptides remaining. The missing values were imputed using Perseus^41^. An analysis of variance (ANOVA) was performed at each time point, and 8,401 peptides that exhibited significant changes at least at one time point were selected for further analysis (**Figure 4a**). To characterize time-dependent changes of phosphoproteome, PCA and hierarchical clustering analysis were performed for each treatment group, which revealed that the ILG and LPS groups showed different phosphoproteome profiles from the 0-hour time point as early as 15 min and 2 h post-stimulation (**Figure 4b**). These observations are consistent with the idea that phosphorylation-mediated signaling works on a fast timescale, responding within seconds to minutes^42^. In contrast, the GU group exhibited stable phosphoproteome profiles from 0 to 2 hours after LPS stimulation, whereas the phosphoproteome at 24 h substantially differed from that of the other samples (**Figure 4b**). In addition, the number of significantly changed phosphorylated peptides was higher in the ILG group than in the others across the 0-to 24-hour period (**Figure 4c**). Thus, our time-resolved multiomics data characterized ILG, a single bioactive component, as primarily inducing rapid signaling responses via phosphorylation, whereas early metabolic changes were observed in the GU-treated group. These findings indicate that the pharmacological mechanism of GU may involve not only phosphorylation-mediated pathways but also alternative regulatory mechanisms, such as ligand–protein interactions facilitated by its diverse natural compounds.

**Figure 4.**
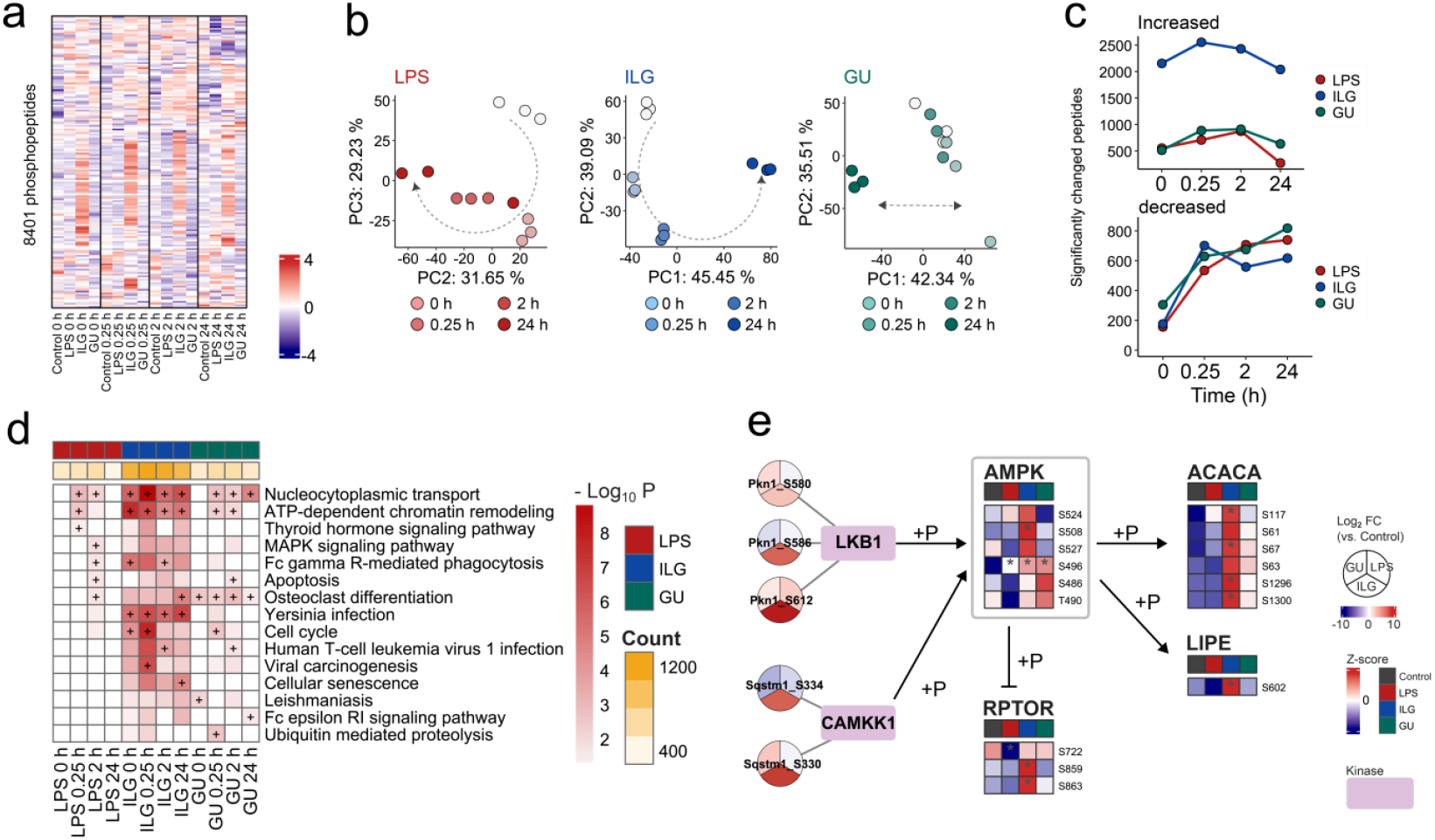
Investigating unique phosphoproteome alterations in ILG- and GU-treated RAW264.7 cells. (a) Heatmap analysis using the z-scaled table of 8401 phosphopeptides. Intensites were normalized by log2 transformation, and missing values were imputed from normal distribution using Perseus software. (b) PCA score plots showing time-dependent changes in the LPS, ILG, and GU groups. (c) Numbers of significantly changed phosphopeptides compared to control at each time point. *P*-values were calculated using the Tukey’s HSD test (two-sided), and phosphopeptides with an absolute log2 fold-change greater than 1 compared to the control were defined as significantly altered phosphopeptides. (d) GSEA using significantly upregulated phosphopeptides. Enrichment analysis was performed based on KEGG pathways using the clusterProfiler package. *P*-value was calculated using Fisher’s exact test and adjusted using the Benjamini-Hachberg method. The yellow color scale represents the number of phosphopeptides that showed a significant increase compared to the control at the corresponding time point. Pathway terms that ranked within the top 5 in each group are indicated by a “+” label. (e) Phosphoproteome and predicted kinase profiles focusing on the AMPK signaling pathway 15 min after LPS stimulation. The responsible kinase was predicted using PhosphoSitePlus based on the amino acid sequence surrounding the phosphorylation site and is indicated by a square node. The pie chart and the heatmap show fold changes against control and z-score values of phosphopeptides, respectively. *P*-values were calculated using Tukey’s HSD test (two-sided) and labeled if *p*-value < 0.05 compared to the control group at the corresponding time point.

To systematically interpret the phosphoproteome changes, we conducted gene set enrichment analysis (GSEA) using the abundance profiles of phosphopeptides and the Kyoto Encyclopedia of Genes and Genomes (KEGG) database^43^. This analysis focused on proteins with significantly altered phosphorylation levels at each time point. In the LPS group, MAPK signaling and phagocytosis pathways, both associated with inflammatory macrophage activation, were significantly enriched 2 h after LPS stimulation (**Figure 4d**). Moreover, pathway terms related to Yersinia infection and osteoclast differentiation, including focal adhesion kinase (FAK) and steroid receptor coactivator (Src) signaling, NF-κB signaling, and MAPK signaling, were significantly enriched in the ILG and GU groups during the 0–24-hour period after stimulation. ILG negatively regulates the downstream signaling of Src via its hydrolyzed metabolite, 2,4,2’,4’-tetrahydroxychalcone, impairing cytoskeletal reorganization and focal adhesion formation, thereby inhibiting macrophage migration^44^. Our findings suggest that the extensive phosphorylation of kinases and adapter proteins within the FAK-Src axis, induced by ILG treatment, represents a potential anti-inflammatory mechanism of ILG.

To identify upstream signaling events specifically responsive to ILG and GU, we performed kinase inference on 191 phosphorylation sites associated with the Yersinia infection and osteoclast differentiation pathways. This analysis inferred the involvement of 88 unique kinases, focusing on phosphorylation sites that showed a selective increase in number in the ILG and GU groups, where we observed liver kinase B1 (LKB1), a key upstream kinase of AMPK, as well as calcium/calmodulin-dependent protein kinase kinase 1, as potential regulators. Consistent with this, the phosphorylation level of AMPK was elevated in the ILG and GU groups (**Figure 4e**). Furthermore, phosphorylation of S722 of the regulatory-associated protein of mTOR (RPTOR) was restored to control levels by ILG and GU treatment. The S722 of mTOR is phosphorylated by AMPK and negatively regulates the mTORC1 complex^45^. ILG and GU treatment also enhanced the phosphorylation of acetyl-CoA carboxylase (ACACA) and hormone-sensitive lipase (LIPE), direct substrates of AMPK. This may explain the observed lipidomic alterations, characterized by reduced TG accumulation and increased DG and MG levels (**Figures 3d, 3e, and Figure 4e**). These results suggest that GU and its active component, ILG, exert their anti-inflammatory effects through the activation of AMPK signaling.

### Integrating multiomics data by a partial least square with the rank order of groups method

We used multiset Partial Least Squares with Rank Order of Groups (PLS-ROG)^29^ to integrate multiomics data unbiasedly. This approach uses a rank order’s information as response variables and calculates *p*-values based on the loadings for latent variables, enabling the systematic extraction of metabolites. In this study, IL-6 expression levels were used to define the rank order (**Figures S4a and S4b**). The integrated result of the multiomics data at 24 h after LPS stimulation revealed that the first latent variable axis reflected differences between the ILG and GU groups, while the second latent variable axis reflected the order of IL-6 expression levels **(Figure 5a)**. Using the loadings and *p*-values on the second latent variable axis for pathway analysis provided the enrichment terms of arginine metabolism and TCA cycle, consistent with the observations in the hydrophilic metabolome section of this study (**Figures S4c, 2c, and 2f**). To further characterize the unique metabolic responses of GU and ILG, pathway analysis was conducted using the metabolites and phosphoproteins having a *p*-value less than 0.05 for the first latent variable. The GU group significantly enriched the pantothenate and acetyl coenzyme A (CoA) biosynthetic pathway, while the ILG group exhibited enrichment of the NAD metabolic pathway, including nicotinamide and SIRT1/2 (**Figure 5c**). The level of pantothenate increased in the GU group from 1 h after stimulation, with significant accumulation of acetyl-CoA at 24 h compared to the control and LPS groups (**Figure S4d**). In the ILG group, nicotinamide accumulation was observed as early as 1 h after stimulation and increased at 24 h (**Figure 5c**). Phosphorylation of SIRT, an enzyme in the NAD^+^ salvage pathway, was also upregulated from 0 to 24 h post-stimulation (**Figures 5d and 5e**). Protein levels of SIRT1 and SIRT2 did not change significantly 15 min after stimulation, with or without LPS or ILG treatment, indicating that phosphorylation, rather than protein expression, was the primary driver of SIRT regulation (**Figure S4e**). The SIRT family is an NAD-dependent enzyme family that regulates transcription factors such as NF-κB, p38, and forkhead box proteins via histone deacetylation activity^46,47^. The phosphorylation of SIRT1 at Ser46 by MAPK8 enhances its nuclear localization and enzymatic activity, while the CDK-dependent phosphorylation of SIRT2 at Ser368 delays cell cycle progression, which suggests that SIRT phosphorylation plays a crucial role in regulating enzymatic activity, thereby influencing inflammatory macrophage function and immune responses^48,49^. To further evaluate the anti-inflammatory effects of SIRT activity, RAW264.7 cells were treated with the SIRT1 inhibitor (EX527) or SIRT2 inhibitor (AGK-2) in the presence of ILG. The inhibition of IL-6 production by ILG was reduced by co-treatment with SIRT2 inhibitor (**Figures 5f and S4f**), although few changes were observed in using two inhibitors simultaneously. Nevertheless, these results indicate that ILG suppresses inflammation via SIRT phosphorylation partially, leading to changes in its enzymatic activity.

**Figure 5.**
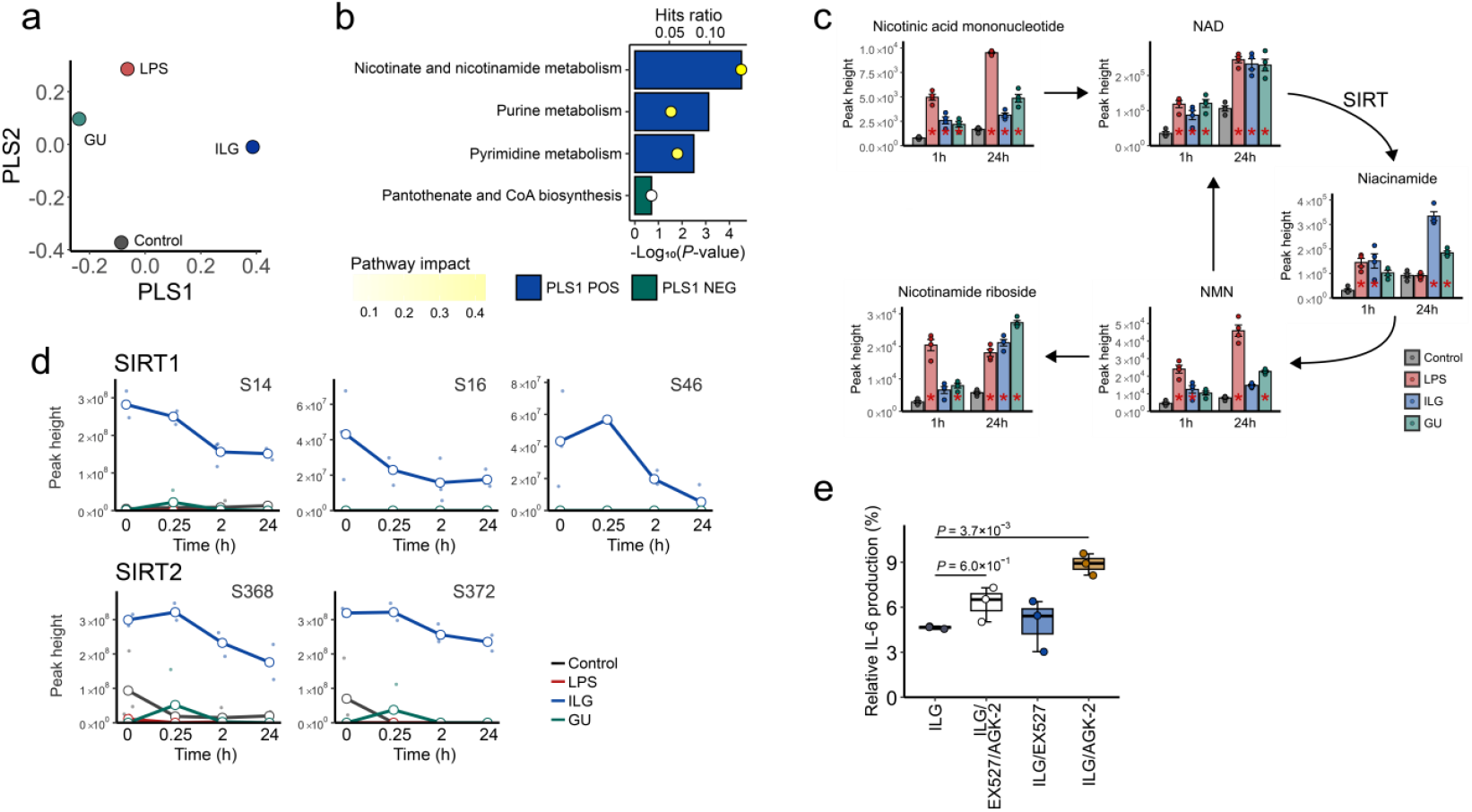
Integrated omics analysis by multiset PLS-ROG showing ILG-dependent unique molecule signatures. (a) The scatter plots using the first and second latent variables with multiset PLS-ROG algorithm. Multiomics data of 24 h after LPS stimulation were subjected to multiset PLS-ROG analysis. (b) Results of MetaboAnalyst joint-pathway enrichment analysis using the first latent variable axes. Bar graphs represent enriched pathways extracted based on *P*-values and loadings. Metabolites and phosphoproteins that reflect group-specific characteristics were extracted and provided for enrichment analysis of MetaboAnalyst. *P*-value was calculated by Fisher’s exact test, and degree centrality was used for the topology measure. The hit ratio means the proportion of features hit out of the total features, and the yellow colour scale indicates the impact of the pathway. (c) NAD metabolism pathway 24 h after LPS stimulation. Bar plots represent the mean values for each group, with individual data points overlaid as a scatter plot and error bars indicate the SEM. *P*-values were calculated using Tukey’s HSD test (two-sided) and labeled if *P*-value < 0.05 compared to the control group at the same time point. (d) Time-dependent changes in phosphorylation levels of SIRT1 and SIRT2. The phosphorylation sites are shown in the plot titles. (e) IL-6 production with or without SIRT1 and SIRT2 inhibitors following 18 h of LPS stimulation. RAW264.7 cells were incubated with EX527 (1 μM) or/and AGK-2 (10 μM) for 1 hour with ILG (10 μM), followed by administration of LPS (*n* = 3 biologically independent samples). *P*-values compared to the ILG group were calculated using the Tukey HSD.

### Elucidating the production mechanism and biological importance of GABA in GU-treated macrophages

Our untargeted metabolomic analysis captured the accumulation of GABA and its downstream metabolite 4-guanidinobutanoic acid as specific metabolic signatures in the GU treatment group. We estimated the concentration of GABA in GU using LC-MS/MS as low as 1.4 fmol/mg (**Figure S5a**), suggesting that GABA accumulation in the GU group would be due to biosynthesis of GABA from the substrates with the responsible enzyme rather than cellular uptake of GABA via the transporter. Nevertheless, we first evaluated the transporter activity using a stable isotope metabolite of GABA-d6, where RAW264.7 cells were incubated with 1 μM of GABA-d6 under LPS stimulation with or without GU addition because a previous study reported that murine primary peritoneal exudate macrophages expressed GAT2 (SLC6A12) and GAT3 (SLC6A13), two of the four GABA transporter subtypes in mice^50^. As a result, GU treatment did not significantly increase GABA isotope levels, while non-labeled GABA accumulated significantly under the GU administration conditions (**Figure 6a**). Moreover, a significant increase in endogenous GABA levels was also observed 2 h after GU addition, regardless of LPS stimulation. Because the concentration of GABA in the GU used in this study (500 μg/mL) was low (approximately 0.7 nM), these findings also suggest that endogenous GABA production can be higher than the contribution from GABA uptake via its transporter. We therefore evaluated GABA accumulation in GU-treated RAW264.7 cells in the presence of allylglycine (AG), a selective GAD inhibitor. As a result, GU treatment induced early GABA accumulation, showing a peak intensity reaching approximately 50% of the 10 μM GABA-d6 used as an internal standard (**Figures 6b and S5b**). Furthermore, GABA production was significantly reduced by AG. Although this suppression did not fully restore GABA levels to those observed in the control group, our findings suggest that endogenous GABA synthesis via GAD is a contributor to the GU-induced increase in GABA.

**Figure 6.**
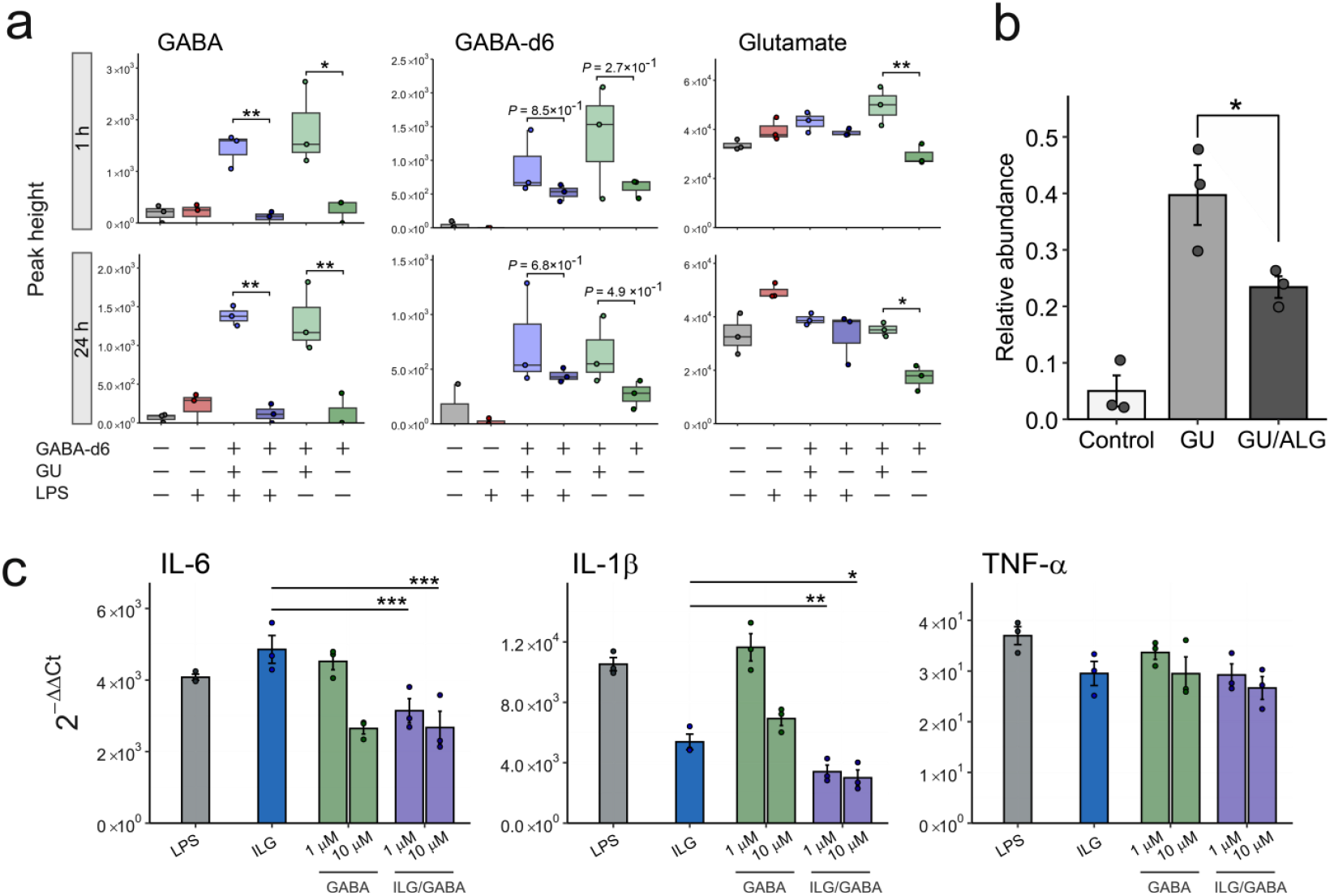
Investigating GABA production mechanism and biological importance in anti-inflammatory effect. (a) Peak intensity of GABA, GABA-d6 isotope, and glutamate following GU treatment with or without LPS stimulation. Hydrophilic metabolome analysis of RAW264.7 cells was performed at 1 and 24 h after LPS stimulation using LC-MS/MS (*n*= 3 biologically independent samples). (b) Relative abundance of GABA in GU-treated RAW264.7 cells with or without preincubation of allylglycine (ALG) (10 mM). The peak height of GABA in each sample was normalized with the peak height of the internal standard, GABA-d6. The values are presented as the means ± SEMs with *n*= 3 biologically independent samples. (c) Effects of GABA and/or ILG on mRNA expression of inflammatory cytokines in RAW264.7 cells. RAW264.7 cells were treated with GABA, ILG (10 μM), or their combination, and the mRNA expression levels of inflammatory cytokines were analyzed. Each bar plot is the mean value of 2^-*ΔΔCt*^ values with error bars representing SEM. The *Ct* values of GAPDH were used for normalization in each sample, with untreated cells serving as the calibrators for each gene. *P*-values for (a), (b), and (c) were calculated using the Tukey HSD (two-sided) and indicated as follows: *P* < 0.05 (*), *P* < 0.01 (**).

We further evaluated the biological activity of GABA in RAW264.7 cells because GABA is a well-characterized inhibitory neurotransmitter with documented anti-inflammatory properties, including suppression of inflammatory cytokine expression in immune cells and alleviation of LPS-induced sepsis. Our result showed that GABA treatment significantly reduced LPS-induced IL-6 and IL-1β expression in stimulated RAW264.7 cells (**Figure 6c**). This anti-inflammatory effect was further validated by GABA receptor agonist treatment and was mitigated by antagonist treatment (**Figure S5c**). Moreover, simultaneous treatment of ILG and GABA further suppressed IL-6 and IL-1β expression compared to treatment with ILG alone (**Figure 6c**). In particular, even when GABA or ILG alone did not significantly reduce cytokine expression levels, their combination showed a significant suppression effect. These results indicate that the combination of natural products from GU and the metabolic changes in mammalian cells induced by GU components would exert synergistic anti-inflammatory effects.

## Discussion

We conducted untargeted multiomics profiling of LPS-stimulated macrophages treated with GU and its bioactive component, ILG, focusing on the hydrophilic metabolome and lipidome, in addition to the phosphoproteome. Our time-course multiomics analysis revealed that ILG and GU induce more pronounced fluctuations in the phosphoproteome and metabolome, respectively, compared to LPS stimulation, with these changes emerging early and persisting over time (**Figures 2b, 3c, and 4c**). This tendency would potentially suggest the existence of unique molecular machinery in GU- and ILG treatment groups for LPS-induced acute inflammatory conditions.

Our metabolomics data showed substantial changes in citrulline and itaconate in GU and ILG treated groups. Previous studies have reported that ILG suppresses the expression of inducible nitric oxide synthase (iNOS) at the mRNA and protein levels, and the decrease of citrulline levels in the ILG-treated group would reflect the decrease of iNOS production levels^12,51^. The levels of itaconate, a known immunomodulatory metabolite produced by the LPS-inducible enzyme immune-responsive gene 1, also exhibited a significant increase in the LPS-treated group. Itaconate plays a critical role in inflammatory responses by covalently modifying Keap1, activating the Nrf2 pathway and suppressing the production of pro-inflammatory cytokines, including IL-6 and IL-1β^52^. Our findings suggest that the early accumulation of itaconate induced by *G. uralensis*-derived natural products contributes to the attenuation of excessive immune responses. Moreover, the phosphoproteome analysis in this study suggests that ILG can modulate AMPK activity and its substrates, ACACA and LIPE, through phosphorylation, thereby promoting β-oxidation and suppressing lipid droplet formation. In addition, multiset PLS-ROG highlighted enhanced SIRT phosphorylations and NAM accumulation as key metabolic responses to ILG treatment, and further investigation indicated that SIRT*2* inhibitor partly suppressed the anti-inflammatory effect of ILG. Previous studies indicate that SIRT and AMPK are functionally interconnected, with AMPK activation depending on increased NAD^+^ levels, the NAD/NADH ratio, and/or nicotinamide phosphoribosyltransferase activity and expression, ultimately leading to SIRT activation. In contrast, SIRT deacetylates and activates LKB1, which would subsequently activate AMPK^53^. Taken together, our results suggest that the metabolic responses induced by ILG—including the accumulation of DG, MG, and free fatty acids, as well as increased levels of NAD cycle intermediates—can be mediated through positive feedback regulation of the SIRT-AMPK relationship.

In addition, we characterized the unique lipidome profiles in both GU and ILG treatment conditions. While GU and ILG suppressed the accumulation of TG in inflammatory macrophages, the levels of TG, which has saturated fatty acids, increased in the GU group (**Figures 3f and S3d**). These findings suggest that GU enhances fatty acid supply through de novo fatty acid synthesis and acyl chain incorporation to TG. However, no changes were observed in the phosphorylation levels of key enzymes involved in fatty acid and TG synthesis, including FASN (fatty acid synthase), stearoyl-CoA 9-desaturase, glycerol-3-phosphate acyltransferase, or diacylglycerol acyltransferase, upon GU treatment. Thus, the mechanism for saturated fatty acid containing TG accumulation in GU treatment group is unclear in our study, while they may be elucidated by shotgun proteomics and/or transcriptomics analyses. Nevertheless, the lipidomic shifts may reflect a weaker anti-inflammatory effect in GU than ILG because saturated fatty acid can be upregulated via increased FASN expression level in inflammatory macrophages^54,55^. For the distinct lipidome changes in the ILG group, Hex2Cer accumulation was observed 24 h after LPS stimulation. A similar metabolic shift occurred in the LPS group, while ILG and LPS exhibited selective enrichment in Hex2Cer species containing sphinganine (18:0;O2) and sphingosine (18:1;O2) as long-chain bases, respectively (**Figure 3g**). This suggests that ILG would facilitate desaturation in sphingobase via DEGS (dihydroceramide desaturase) enzymes with a dysfunction that is linked to increased levels of reactive oxygen species and ER stress^56,57^.

Moreover, we focused on the early accumulation of GABA as a hallmark of the GU-treated RAW264.7 cells in the hydrophilic metabolome, which was most dramatically altered by GU treatment. Our results suggested that the increase in GABA levels is primarily driven by endogenous synthesis via GAD rather than exogenous uptake. Previous studies have reported that GABA receptor activation negatively regulates NF-κB and NOD-like receptor protein 3 signaling^58,59^. Consistent with these findings, our results demonstrated that GABA treatment significantly suppressed IL-6 and IL-1*β* mRNA expression, although these inhibitory effects were only reproduced partially through muscimol or baclofen treatments. Additionally, the combination of GABA and ILG suppressed inflammatory marker expression to levels greater than the additive anti-inflammatory effects of each compound alone. This result indicates that endogenously biosynthesized GABA not only functions as a direct anti-inflammatory mediator in LPS-induced inflammation but also synergistically enhances anti-inflammatory effects in combination with other GU components such as ILG. Consequently, our study highlighted the utility of multiomics approaches in resolving the metabolic dynamics in herbal medicine treatments and offer a paradigm for understanding how component interactions shape therapeutic effects.

## Data availability

The mass spectrometry raw data of hydrophilic metabolomics, lipidomics, and phosphoproteomics are available on the RIKEN DROP Met repository (http://prime.psc.riken.jp) under the index number DM0077. Source Data for figures are provided with this paper.

## Acknowledgments

This study represents a portion of the dissertation submitted by Saki Kiuchi to the Tokyo University of Agriculture and Technology in partial fulfillment of the requirements for her Ph.D. This study was supported by the JSPS KAKENHI (21K18216, 24K02011, 24H00043, 24H00392, 24K21269, H.T.), National Cancer Center Research and Development Fund (2023-A-08, H.T.), JST National Bioscience Database Center (JPMJND2305, H.T.), JST FOREST program (JPMJFR230H, H.T.), JST ERATO “Arita Lipidome Atlas Project” (JPMJER2101, M.A. and H.T.) and Technologically Advanced research through Marriage of Agriculture and engineering as Groundbreaking Organization (TAMAGO to J.M. and Hiroshi T.).

## Author Contributions

H.T. designed and managed the study. T.N. and K.O. provided *Glycyrrhiza uralensis* extracts and supported the experiments for this research. S.K., K.T., and K.I. performed the phosphoproteomics analysis. S.K., Y.O., T.N., H.Y., and K.S. performed the hydrophilic metabolome analysis. S.K. and M.H.C. performed experiments using ELISA, qPCR, and western blotting. S.K. and H.T. wrote the paper, and all authors have thoroughly discussed this project and helped improve the manuscript.

