## Supporting information for "Unraveling anti-inflammatory metabolic signatures of *Glycyrrhiza uralensis* and isoliquiritigenin through multiomics"

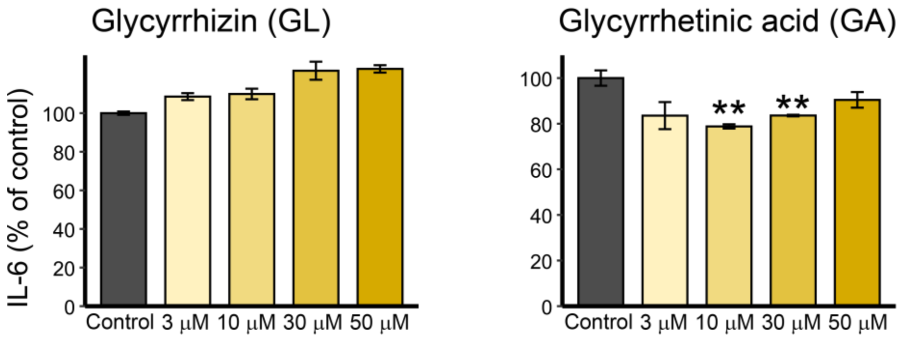


**Supplementary Figure 1. IL-6 productions in RAW264.7 cells incubated with glycyrrhizin or glycyrrhetinic acid.** Following 24 h of lipopolysaccharide stimulation (100 ng/mL), the culture supernatants were collected, and the production of IL-6 was quantified using ELISA ($n$ = 4 for GL and $n$ = 3 for GA, biologically independent samples). Relative IL-6 levels compared to the LPS group were presented for bar plots as mean ± standard error. $P$-values were calculated using Tukey's honestly significant difference (HSD) test (two-sided), and those comparing each condition to the LPS group were displayed above the corresponding bars.

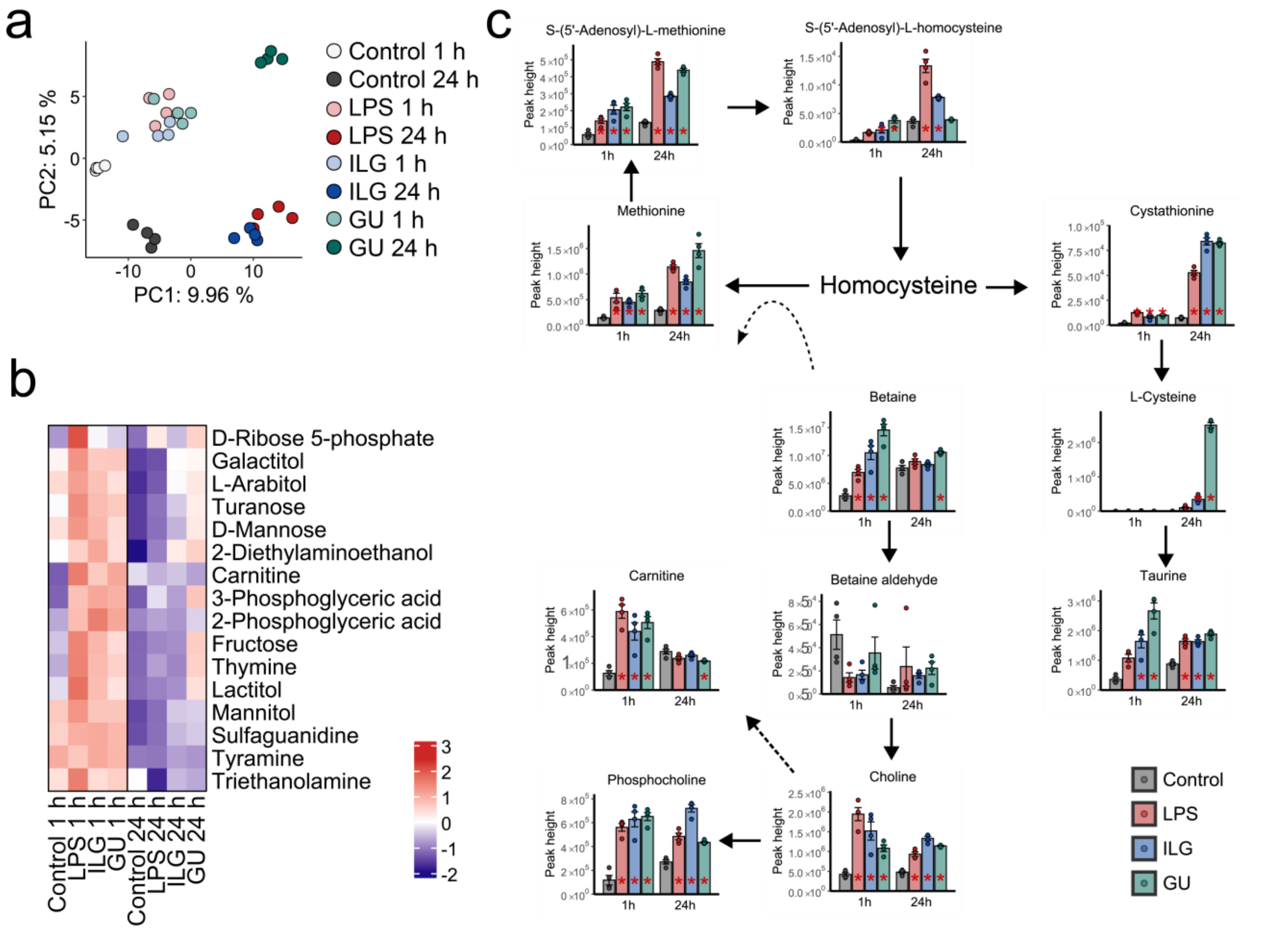


**Supplementary Figure 2. Hydrophilic metabolome alterations among four treatment groups in RAW264.7 cells.** (a) PCA score plot using the auto-scaled data that contains all biological samples acquired in this study. (b) Heatmap of metabolites associated with the cluster that showed an increase in number at 1 h after LPS stimulation. Z-scaled data were used for the clustering analysis. (c) Metabolic pathways of choline metabolism at 1 and 24 h after LPS stimulation. *P*-values were calculated using the Tukey HSD and annotated when $p < 0.05$ compared to the control group at the corresponding time point. Each bar plot is the mean of corrected peak height with error bars representing the standard error of the mean (SEM).

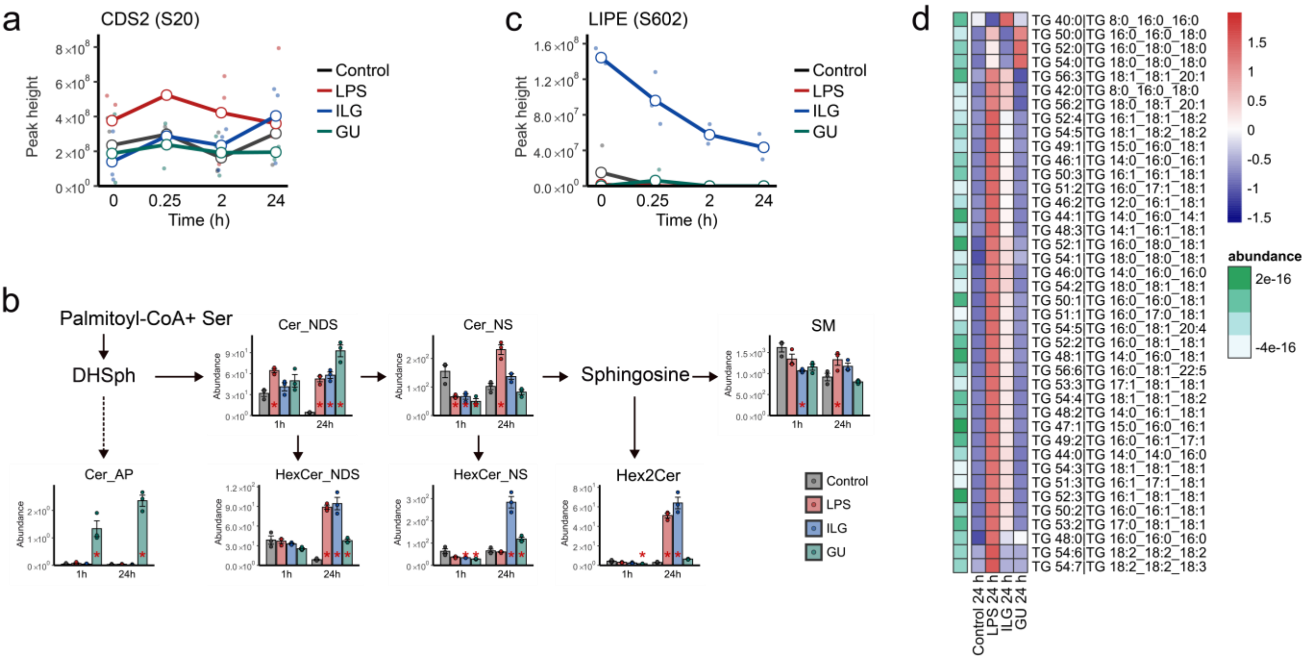


**Supplementary Figure 3. Characteristics of lipidome alterations among four treatment groups in RAW264.7 cells.** (a) Time-resolved changes in phosphorylation levels of phosphatidate cytidylyltransferase 2 (CDS2) at S20. (b) Mapping the lipidome profile onto ceramide metabolism. *P*-values were calculated using the Tukey HSD and annotated when $p < 0.05$ compared to the control group at the corresponding time point. Each bar plot is the means of normalized abundance with error bars representing the standard error of the mean (SEM). (c) Time-resolved changes in phosphorylation levels of hormone-sensitive lipase (LIPE) at S602. (d) Heatmap of triacylglycerol (TG) showing ion abundance ratios among four groups at 24 h after LPS stimulation. The green color scale (left) shows the log2-transformed value of summed normalized abundances of each lipid among samples.

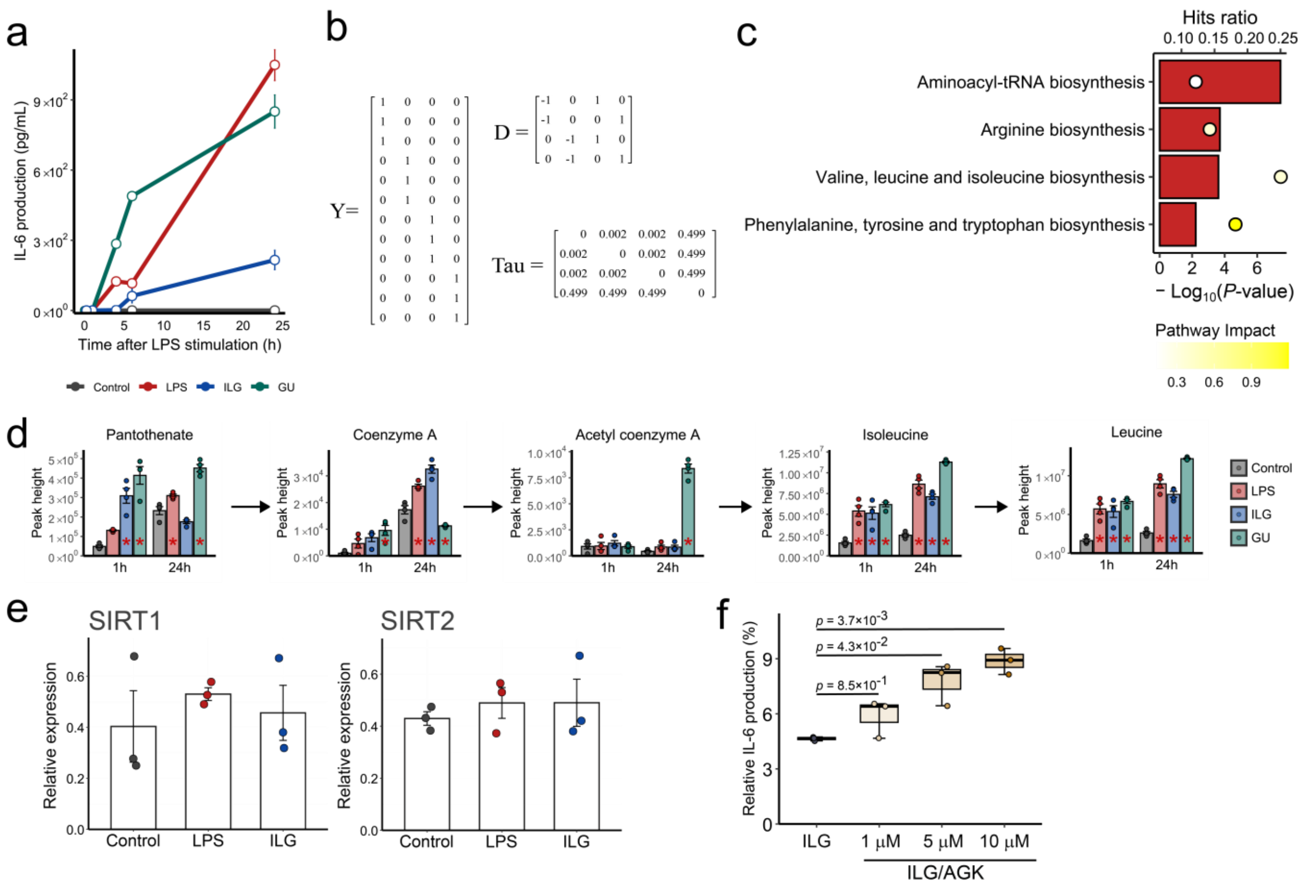


**Supplementary Figure 4. Using multiset partial least square with rank order of the biological groups (PLS-ROG) to interpret multiomics data.** (a) Time-resolved IL-6 production of LPS-stimulated RAW264.7 cells. ILG or GU was administered 1 h before LPS stimulation. Following 0, 0.25, 4, 6, and 24 h, the cell culture supernatant was collected, and IL-6 levels were determined by ELISA (*n*= 4 biologically independent samples). Data are expressed as the mean ± SEM. (b) Parameter settings of multiset PLS-ROG used in this study. The terms Y, D, and tau represent matrices that set the response variable, the order of the groups, and the strength of the coupling between the t data or between the groups and each data, respectively. (c) Results of joint-pathway enrichment analysis of MetaboAnalyst for the second latent variable axes. The hit ratio means the proportion of features hit out of the total features, and the yellow colour scale indicates the impact of the pathway. (d) Metabolic pathway of CoA biosynthesis. Bar plots represent the mean values for each group, with individual data points overlaid as a scatter plot. Error bars indicate the standard error of the mean (SEM) and P-values were calculated using the Tukey's HSD test (two-sided) and labeled if $p < 0.05$ compared to the control group at the same time point. (e) Protein expression levels of SIRT1 (left) and SIRT2 (right). RAW264.7 cells 15 min after LPS stimulation were collected, and relative expression level was determined by western blotting (*n*= 3 biologically independent samples). Untreated RAW264.7 cells were used as vehicle control. Each bar plot is the mean of relative protein expression levels with error bars representing the standard error of the mean (SEM). Band intensity of GAPDH was used for the normalization of SIRT expression levels. (f) IL-6 production of LPS-stimulated RAW264.7 cells incubated with ILG and/or AGK-2 (*n*=3 biologically independent samples). The cell culture supernatant was collected 24 h after LPS stimulation. *P*-values compared to the ILG group were calculated using the Tukey HSD (two-sided).

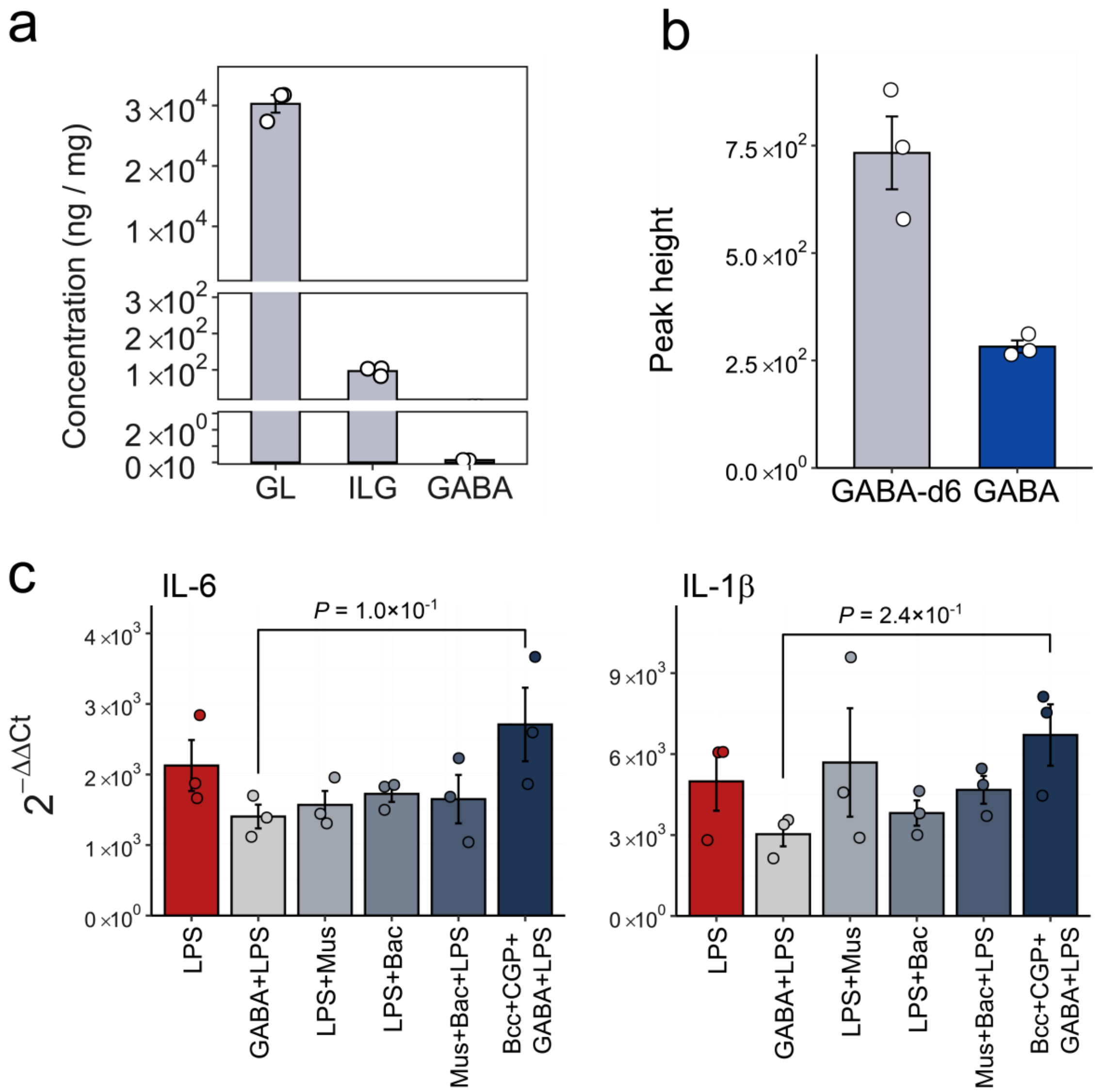


**Supplementary Figure 5. Elucidating the production mechanism and biological importance of GABA in GU-treated macrophages.** (a) GABA concentration in GU. Hydrophilic metabolites were extracted from GU and quantified using LC-MS/MS. Data are expressed as the mean ± SEM of GABA per mg of GU. (b) Peak height of internal standard and GABA in GU-treated RAW264.7 cells. GABA-d6 with a final concentration of 10 µM was used as an internal standard. The values are presented as the means ± SEMs with n=3 biologically independent samples. (c) mRNA expressions of inflammatory cytokine in RAW264.7 cells incubated with GABAR1/2 agonist or antagonist. The cells were incubated for 1 h with muscimol (Mus, GABAR1 agonist) at 2 µM, baclofen (Bac, GABAR2 agonist) at 50 µM, bicuculline (Bcc, GABAR1 antagonist) at 50 µM, CGP 55845 (CGP, GABAR2 antagonist) at 5 µM, and GABA (1 µM), followed by LPS stimulation for 2 h ($n$=3 biologically independent samples). Each bar plot is the mean value of $2^{-\Delta\Delta Ct}$ values with error bars representing SEM. The *Ct* values of GAPDH were used for normalization in each sample, with untreated cells serving as the calibrators for each gene. Statistical significance was determined using Tukey's HSD test (two-sided) comparing each treatment to the LPS group.

**Supplementary tables**

**Table S1** The primer sequences for RT-qPCR

| Target gene | Forward | Reverse |
|---|---|---|
| *Gapdh* | 5′-TGCCACCTTTTGACAGTGATG-3′ | 5′-TGATGTGCTGCTGCGAGATT-3′ |
| *Il-6* | 5′-AGTTGCCTTCTTGGGACTGA-3′ | 5′-CAGAATTGCCATTGCACAAC-3′ |
| *IL-1β* | 5′-TGTGTCCGTCGTGGATCTGA-3′ | 5′-TTGCTGTTGAAGTCGCAGGAG-3′ |
| *Tnf-α* | 5′-GTGGAACTGGCAGAAGAGGC-3′ | 5′-AGACAGAAGAGCGTGGTGGC-3′ |
