## Supplementary material for "Unraveling anti-inflammatory metabolic signatures of *Glycyrrhiza uralensis* and isoliquiritigenin through multiomics": Source data: Source code for Figure 1 (ELISA).Rmd

````
---
title: "R Notebook"
output: html_notebook
---

This is an [R Markdown](http://rmarkdown.rstudio.com) Notebook. When you execute code within the notebook, the results appear beneath the code. 

Try executing this chunk by clicking the *Run* button within the chunk or by placing your cursor inside it and pressing *Ctrl+Shift+Enter*. 

```{r Source code for Figure 1a and S1}
library(exactRankTests)
library(ggplot2)
library(dplyr)
library(openxlsx)
library(ggsignif)
library(plotrix)
library(loadings)
library(tidyverse)
library(pheatmap)
library(plotrix)
library(DescTools)
library(scales)


setwd("D:/2024/01_kanzo_multiomics/05_paper_work/Source data/1_ELISA")
df <- read.xlsx("Source-data_ELISA.xlsx", 1)

df_unkwn <- df %>% filter(Class != "Standard" & Class != "Control") %>% mutate(across(5:9, as.numeric))
Experiment_num <- df_unkwn %>% distinct(Experiment.No.) %>% unlist()

#ILG
tardf <- df_unkwn %>% filter(Experiment.No. == Experiment_num[2]) %>% filter(Group %in% c("LPS", "ILG"))
mean_LPS <- tardf %>% filter(Group == "LPS") %>% .$`IL-6(pg/mL)` %>% mean()
tardf.2 <- tardf %>% mutate("percent_of_LPS" = .$`IL-6(pg/mL)`/mean_LPS*100) %>% separate(Class, into = c("First", "Second"), sep = "_", remove = FALSE) %>% mutate(Class_label = ifelse(Group == "LPS", First, Second))
group <- tardf.2 %>% distinct(Group) %>% unlist()


pallete <- colorRampPalette(c("#BBDEFB", "#0D47A1"))

     class <- tardf.2 %>% distinct(Class) %>% unlist()
     tardf.2 <- transform(tardf.2 , Class = factor(Class, levels = class),
                          Class_label = factor(Class_label, levels = distinct(tardf.2, Class_label) %>% unlist()))

     df_summary <- tardf.2 %>% group_by(Class_label) %>% summarise(mean_value = mean(percent_of_LPS, na.rm = TRUE))
    
    
    Tukey_result <- TukeyHSD(aov(tardf.2$percent_of_LPS ~ tardf.2$Class_label))
    Tukey_result_df <- data.frame(Comparison = rownames(Tukey_result$`tardf.2$Class_label`),
                                  Tukey_result$`tardf.2$Class_label`) %>% filter(grepl("-LPS", Comparison)) %>%
      mutate(p_value = signif(p.adj, 2)) %>% mutate(Class_label = distinct(tardf.2, Class_label) %>% unlist() %>% .[c(2,3,4)])
    # dunn_result <- dunnTest(percent_of_LPS ~ Class_label, data = tardf.2, method = "bh")
    # dunn_result_df <- dunn_result$res %>% filter(grepl("- LPS", Comparison))
    p_txt <- df_summary %>% left_join(Tukey_result_df, by = "Class_label")
     
 
    p <- ggplot(tardf.2, aes(x = Class_label, y = percent_of_LPS, group = Class_label, fill = Class_label))+
        stat_summary(fun = "mean", geom = "bar", width = .7, col= "black") +
        geom_jitter(shape =21, width = .1, size = 3, fill = "white")+
        stat_summary(fun.max = function(x) mean(x) + std.error(x), 
                 fun.min = function(x) mean(x) - std.error(x), 
                 geom = 'errorbar', width = .2, linewidth = 0.5)+
     
      labs(y = "IL-6 (% of LPS)")+
      scale_fill_manual(values = c("grey30", pallete(length(class)-1)))+
      scale_y_continuous(expand = c(0,0), breaks = seq(0,100,20), limits = c(0, max(tardf.2$percent_of_LPS,na.rm = TRUE)*1.1))+
      theme(
        legend.title = element_blank(),
        legend.position = "right",
        panel.background = element_rect(fill = "transparent", colour = NA),
        axis.line = element_line(colour="black", size = 1),
        axis.title.y = element_text(size = 12,colour="black"),
        axis.title.x = element_blank(),
        axis.text = element_text(size = 10,colour="black"),
        panel.grid = element_blank(),
        axis.ticks.x=element_blank(),
        aspect.ratio = 0.8)+
     geom_text(data = p_txt[-1,], aes(x = Class_label, y = mean_value * 1.1,label = p_value), vjust = 0, size = 3)

    p

  svg("ILG.svg", width = 4,height = 3)
  print(p)
  dev.off()

  
  #GU
  tardf <- df_unkwn %>% filter(Experiment.No. == Experiment_num[1]) %>% filter(Group %in% c("LPS", "GU"))
  mean_LPS <- tardf %>% filter(Group == "LPS") %>% .$`IL-6(pg/mL)` %>% mean()
  tardf.2 <- tardf %>% mutate("percent_of_LPS" = .$`IL-6(pg/mL)`/mean_LPS*100) %>%
    separate(Class, into = c("First", "Second"), sep = "_", remove = FALSE) %>% mutate(Class_label = ifelse(Group == "LPS", First, Second))
  group <- tardf.2 %>% distinct(Group) %>% unlist()

  pallete <- colorRampPalette(c("#b4ded9ff","#00695cff"))


  

   class <- tardf.2 %>% distinct(Class) %>% unlist()
   tardf.2 <- transform(tardf.2 , Class = factor(Class, levels = class),
                          Class_label = factor(Class_label, levels = distinct(tardf.2, Class_label) %>% unlist()))
   df_summary <- tardf.2 %>% group_by(Class_label) %>% summarise(mean_value = mean(percent_of_LPS, na.rm = TRUE))
    
    
    Tukey_result <- TukeyHSD(aov(tardf.2$percent_of_LPS ~ tardf.2$Class_label))
    Tukey_result_df <- data.frame(Comparison = rownames(Tukey_result$`tardf.2$Class_label`),
                                  Tukey_result$`tardf.2$Class_label`) %>% filter(grepl("-LPS", Comparison)) %>%
      mutate(p_value = signif(p.adj, 2)) %>% mutate(Class_label = distinct(tardf.2, Class_label) %>% unlist() %>% .[-1])
    
    dunn_result <- dunnTest(percent_of_LPS ~ Class_label, data = tardf.2, method = "bh")
    dunn_result_df <- dunn_result$res %>% filter(grepl("- LPS", Comparison))

    p_txt <- df_summary %>% left_join(Tukey_result_df, by = "Class_label")
     
 
    p <- ggplot(tardf.2, aes(x = Class_label, y = percent_of_LPS, group = Class_label, fill = Class_label))+
        stat_summary(fun = "mean", geom = "bar", width = .7, col= "black") +
       geom_jitter(shape =21, width = .1, size = 3, fill = "white")+
        stat_summary(fun.max = function(x) mean(x) + std.error(x), 
                 fun.min = function(x) mean(x) - std.error(x), 
                 geom = 'errorbar', width = .2, linewidth = 0.5)+
      
      labs(y = "IL-6 (% of LPS)")+
      scale_fill_manual(values = c("grey30", pallete(length(class)-1)))+
      scale_y_continuous(expand = c(0,0), breaks = seq(0,120,20), limits = c(0, max(tardf.2$percent_of_LPS,na.rm = TRUE)*1.2))+
      theme(
        legend.title = element_blank(),
        legend.position = "right",
        panel.background = element_rect(fill = "transparent", colour = NA),
        axis.line = element_line(colour="black", size = 1),
        axis.title.y = element_text(size = 12,colour="black"),
        axis.title.x = element_blank(),
        axis.text.x = element_text(angle = 20),
        axis.text = element_text(size = 10,colour="black"),
        panel.grid = element_blank(),
        axis.ticks.x=element_blank(),
        aspect.ratio = 0.7)+
     geom_text(data = p_txt[-1,], aes(x = Class_label, y = mean_value * 1.15,label = p_value), vjust = 0, size = 3)

    p

  svg("GU.svg", width = 4.5,height = 3)
  print(p)
  dev.off()

  
  
  
  #Figure S1
  #GA
  tardf <- df_unkwn %>% filter(Experiment.No. == Experiment_num[2]) %>% filter(Group %in% c("LPS", "GA"))
  mean_LPS <- tardf %>% filter(Group == "LPS") %>% .$`IL-6(pg/mL)` %>% mean()
  tardf.2 <- tardf %>% mutate("percent_of_LPS" = .$`IL-6(pg/mL)`/mean_LPS*100) %>%
    separate(Class, into = c("First", "Second"), sep = "_", remove = FALSE) %>% mutate(Class_label = ifelse(Group == "LPS", First, Second))
  group <- tardf.2 %>% distinct(Group) %>% unlist()

  pallete <- colorRampPalette(c("#fff2c0ff", "#d6aa00ff"))

  class <- tardf.2 %>% distinct(Class) %>% unlist()
  tardf.2 <- transform(tardf.2 , Class = factor(Class, levels = class),
                          Class_label = factor(Class_label, levels = distinct(tardf.2, Class_label) %>% unlist()))
  df_summary <- tardf.2 %>% group_by(Class_label) %>% summarise(mean_value = mean(percent_of_LPS, na.rm = TRUE))
    
    
    Tukey_result <- TukeyHSD(aov(tardf.2$percent_of_LPS ~ tardf.2$Class_label))
    Tukey_result_df <- data.frame(Comparison = rownames(Tukey_result$`tardf.2$Class_label`),
                                  Tukey_result$`tardf.2$Class_label`) %>% filter(grepl("-LPS", Comparison)) %>%
      mutate(p_value = signif(p.adj, 2)) %>% mutate(Class_label = distinct(tardf.2, Class_label) %>% unlist() %>% .[-1])
    # dunn_result <- dunnTest(percent_of_LPS ~ Class_label, data = tardf.2, method = "bh")
    # dunn_result_df <- dunn_result$res %>% filter(grepl("- LPS", Comparison))
    p_txt <- df_summary %>% left_join(Tukey_result_df, by = "Class_label")
     
     p <- ggplot(tardf.2, aes(x = Class_label, y = percent_of_LPS, group = Class_label, fill = Class_label))+
        stat_summary(fun = "mean", geom = "bar", width = .7, col= "black") +
        geom_jitter(shape =21, width = .1, size = 3, fill = "white")+
        stat_summary(fun.max = function(x) mean(x) + std.error(x), 
                 fun.min = function(x) mean(x) - std.error(x), 
                 geom = 'errorbar', width = .2, linewidth = 0.5)+
     
      labs(y = "IL-6 (% of LPS)")+
      scale_fill_manual(values = c("grey30", pallete(length(class)-1)))+
      scale_y_continuous(expand = c(0,0), breaks = seq(0,100,20), limits = c(0, max(tardf.2$percent_of_LPS,na.rm = TRUE)*1.1))+
      theme(
        legend.title = element_blank(),
        legend.position = "right",
        panel.background = element_rect(fill = "transparent", colour = NA),
        axis.line = element_line(colour="black", size = 1),
        axis.title.y = element_text(size = 12,colour="black"),
        axis.title.x = element_blank(),
        axis.text = element_text(size = 12,colour="black"),
        panel.grid = element_blank(),
        axis.ticks.x=element_blank(),
        aspect.ratio = 0.8)+
     geom_text(data = p_txt[-1,], aes(x = Class_label, y = mean_value * 1.1,label = p_value), vjust = 0, size = 3)

    p

  svg("GA.svg", width = 4.5,height = 3)
  print(p)
  dev.off()
  
  #GL
  tardf <- df_unkwn %>% filter(Experiment.No. == Experiment_num[3]) %>% filter(Group %in% c("LPS", "GL"))
  mean_LPS <- tardf %>% filter(Group == "LPS") %>% .$`IL-6(pg/mL)` %>% mean()
  tardf.2 <- tardf %>% mutate("percent_of_LPS" = .$`IL-6(pg/mL)`/mean_LPS*100) %>%
    separate(Class, into = c("First", "Second"), sep = "_", remove = FALSE) %>% mutate(Class_label = ifelse(Group == "LPS", First, Second))
  group <- tardf.2 %>% distinct(Group) %>% unlist()

  class <- tardf.2 %>% distinct(Class) %>% unlist()
  tardf.2 <- transform(tardf.2 , Class = factor(Class, levels = class),
                          Class_label = factor(Class_label, levels = distinct(tardf.2, Class_label) %>% unlist()))
  df_summary <- tardf.2 %>% group_by(Class_label) %>% summarise(mean_value = mean(percent_of_LPS, na.rm = TRUE))
    
    
    Tukey_result <- TukeyHSD(aov(tardf.2$percent_of_LPS ~ tardf.2$Class_label))
    Tukey_result_df <- data.frame(Comparison = rownames(Tukey_result$`tardf.2$Class_label`),
                                  Tukey_result$`tardf.2$Class_label`) %>% filter(grepl("-LPS", Comparison)) %>%
      mutate(p_value = signif(p.adj, 2)) %>% mutate(Class_label = distinct(tardf.2, Class_label) %>% unlist() %>% .[-1])
    p_txt <- df_summary %>% left_join(Tukey_result_df, by = "Class_label")
     
     p <- ggplot(tardf.2, aes(x = Class_label, y = percent_of_LPS, group = Class_label, fill = Class_label))+
        stat_summary(fun = "mean", geom = "bar", width = .7, col= "black") +
        geom_jitter(shape =21, width = .1, size = 3, fill = "white")+
        stat_summary(fun.max = function(x) mean(x) + std.error(x), 
                 fun.min = function(x) mean(x) - std.error(x), 
                 geom = 'errorbar', width = .2, linewidth = 0.5)+
     
      labs(y = "IL-6 (% of LPS)")+
      scale_fill_manual(values = c("grey30", pallete(length(class)-1)))+
      scale_y_continuous(expand = c(0,0), breaks = seq(0,120,20), limits = c(0, max(tardf.2$percent_of_LPS,na.rm = TRUE)*1.1))+
      theme(
        legend.title = element_blank(),
        legend.position = "right",
        panel.background = element_rect(fill = "transparent", colour = NA),
        axis.line = element_line(colour="black", size = 1),
        axis.title.y = element_text(size = 12,colour="black"),
        axis.title.x = element_blank(),
        axis.text = element_text(size = 12,colour="black"),
        panel.grid = element_blank(),
        axis.ticks.x=element_blank(),
        aspect.ratio = 0.8)+
     geom_text(data = p_txt[-1,], aes(x = Class_label, y = mean_value * 1.1,label = p_value), vjust = 0, size = 3)

    p

  svg("GL.svg", width = 4.5,height = 3)
  print(p)
  dev.off()
  
```


```{r Source code for Figure 5f}

setwd("D:/2024/01_kanzo_multiomics/05_paper_work/figure/05_sirt")
df <- read.xlsx("Source-data_ELISA.xlsx", 2)

df_unkwn <- df %>% filter(Class != "Standard")%>% filter(`IL-6(pg/mL)` != "<0.000") %>% mutate(across(4:8, as.numeric))

class <- df_unkwn %>% distinct(Class) %>% .$Class
group <- df_unkwn %>% distinct(Group) %>% .$Group


df_unkwn <- transform(df_unkwn , Class = factor(Class, levels = class[c(1:3,6,5,4,9,8,7)])) %>%
  mutate(percent_of_LPS = df_unkwn$`IL-6(pg/mL)`/mean(df_unkwn$`IL-6(pg/mL)`[c(1:4)])*100) %>% filter(Class != "LPS")

Tukey_result <- TukeyHSD(aov(df_unkwn$percent_of_LPS ~ df_unkwn$Class))
Tukey_result$`df_unkwn$Class`
write.csv(Tukey_result$`df_unkwn$Class`, "Tukey_result.csv")


plotdf <- df_unkwn %>% filter(Class %in% c("ILG", "ILG/EX10uM/AGK10uM","ILG/EX1uM", "ILG/AGK10uM"))
plotdf <- df_unkwn %>% filter(!(Class %in% c("ILG/EX10uM/AGK10uM")))
plotdf <- df_unkwn %>% filter(Group %in% c("ILG", "ILG/EX")) #EX527
plotdf <- df_unkwn %>% filter(Group %in% c("ILG", "ILG/AGK")) #AGK-2

p <- ggplot(plotdf, aes(x = Class, y = percent_of_LPS, fill = Class))+
  geom_boxplot(width = .6, col = "black",alpha =0.6)+
  stat_boxplot(geom="errorbar", width=0.2) +
  # scale_fill_manual(values = c("#4A4C61","white", "#0D47A1", "#B57000"), labels= c("ILG","ILG/EX527/AGK-2","ILG/EX527","ILG/AGK-2"))+
  # scale_fill_manual(values = c("#4A4C61",colorRampPalette(c("#d8e6fc","#0D47A1"))(3), colorRampPalette(c("#E8D4B2","#B57000"))(3)))+ #AGK-2
  # # scale_fill_manual(values = c("#4A4C61",colorRampPalette(c("#d8e6fc","#0D47A1"))(3)))+ #EX527
  scale_fill_manual(values = c("#4A4C61",colorRampPalette(c("#E8D4B2","#B57000"))(3)))+ #AGK-2
  geom_jitter(size =2, width = .2, shape = 21)+
  scale_y_continuous(limits = c(0, max(plotdf$percent_of_LPS)*1.2), expand = c(0,0))+
  labs(y = "Relative IL-6 production (%)")+
  theme(
    axis.text.x = element_blank(),
    panel.grid = element_blank(),
    axis.text.y = element_text(size = 14, color= "black"),
    axis.title.y = element_text(size = 12, color= "black" ),
    axis.title.x = element_blank(),
    plot.title = element_text(hjust=0.5, size = 18, color= "black"),
    legend.key.size = unit(0.5, 'cm'),
    legend.position = "right",
    legend.text = element_text(size = 12),
    axis.line = element_line(colour="black", size = 0.75),
    panel.background = element_rect(fill = "transparent"),
    legend.title = element_blank(), 
    aspect.ratio = 0.8)


p

# svg("D:/2024/01_kanzo_multiomics/05_paper_work/figure/05_sirt/SIRT-inhibitor_v3.svg", width = 5, height = 3) #5,4
# svg("SIRT-inhibitor_woCombination_v3.svg", width = 5, height = 3) #5,4
# svg("SIRT-inhibitor_EX527_v3.svg", width = 5, height = 3) #5,4
svg("SIRT-inhibitor_AGK2_v4.svg", width = 5, height = 3) #5,4
print(p)
dev.off()

```


```{r Source code for Figure S4a}
scientific_notation <- function(x) {
 
  
  
  
   x <- format(x, scientific = TRUE)
  x <- gsub("^(.*)e", "'\\1'e", x)
  x <- gsub("e", "%*%10^", x)
  x <- gsub('\\+', '', x)
  parse(text = x)
}

setwd("D:/2024/01_kanzo_multiomics/05_paper_work/Source data/1_ELISA")
df <- read.xlsx("Source-data_ELISA.xlsx", 3)
col_groups <- c("#424242", "#B71C1C", "#0D47A1","#00695C")


df_unkwn <- df %>% filter(Class != "Standard") %>% mutate(across(c(9,10), as.numeric))
class <- df_unkwn %>% distinct(Class) %>% .$Class
group <- df_unkwn %>% distinct(Group) %>% .$Group


df_unkwn <- transform(df_unkwn , Class = factor(Class, levels = class[c(1:3,6,5,4,9,8,7)]))

linedf <- df_unkwn %>% .[,c(2:5,9,10)] %>%
  group_by(Time) %>% mutate(percent_of_LPS = (`IL-6(pg/mL)`/ mean(`IL-6(pg/mL)`[Group == "LPS"], na.rm = TRUE)) * 100) %>%
  ungroup() %>% mutate("x_axis" = rep(rep(c(0,0.25,1,4,6,24), each = 4), 4))


linedf <- transform(linedf , Group = factor(Group, levels = group)) %>% mutate(`IL-6(pg/mL)` = replace_na(`IL-6(pg/mL)`, 0))


p <- ggplot(linedf, aes(x = x_axis, y = `IL-6(pg/mL)`, group = Group, col = Group))+
  stat_summary(fun = "mean", geom = "line", linewidth = 1.5) +
  stat_summary(fun.max = function(x) mean(x) + std.error(x), 
               fun.min = function(x) mean(x) - std.error(x), 
               geom = 'errorbar', width = .2, linewidth = 1)+
  stat_summary(fun = "mean", geom = "point",shape =16, size = 3) +
  scale_color_manual(values = col_groups)+
  labs(y = "IL-6 production (pg/mL)", x = "Time after LPS stimulation (h)")+
  scale_y_continuous(expand = c(0,0), labels = scientific_notation)+
  theme(
    legend.title = element_blank(),
    legend.position = "right",
    panel.background = element_rect(fill = "transparent", colour = NA),
    axis.line = element_line(colour="black", size = 1),
    axis.title = element_text(size = 12,colour="black"),
    axis.text = element_text(size = 10,colour="black"),
    panel.grid = element_blank(),
    axis.ticks.x=element_blank(), 
    aspect.ratio = 1.25
  )
  p
svg("D:/2024/01_kanzo_multiomics/05_paper_work/Source data/1_ELISA/timecourse_test.svg", width = 4, height = 4)
print(p)
dev.off()

```
````
