## Supplementary material for "Unraveling anti-inflammatory metabolic signatures of *Glycyrrhiza uralensis* and isoliquiritigenin through multiomics": Source data: Source code for Figure 2 (metabolome).Rmd

````
---
title: "R Notebook"
output: html_notebook
---

This is an [R Markdown](http://rmarkdown.rstudio.com) Notebook. When you execute code within the notebook, the results appear beneath the code. 

Try executing this chunk by clicking the *Run* button within the chunk or by placing your cursor inside it and pressing *Ctrl+Shift+Enter*. 

```{r cell counts correction}
library(exactRankTests)
library(ggplot2)
library(dplyr)
library(openxlsx)
library(ggsignif)
library(plotrix)
library(loadings)
library(tidyverse)
library(pheatmap)
library(plotrix)

setwd("D:/2024/01_kanzo_multiomics/05_paper_work/Source data/2_metabolome")

metabolome <- read.xlsx("Source-data_hydrophipic metabolome.xlsx", 3) #Fig 1c and 2
cellcount <- read.xlsx("Source-data_hydrophipic metabolome.xlsx", 1) %>% .[,-1] #Sample metadata for metabolome
metabolitename <- read.xlsx("Source-data_hydrophipic metabolome.xlsx", 2) %>% .[,9] #Metadata for metabolite name
colnames(metabolome) <- c("Name",	"class", "group", "time", metabolitename)


df_metabolome <- data.frame()
df_info <- data.frame()

for (i in c(1:nrow(cellcount))) {
  # i=1
  df_metabo <- metabolome %>% filter(class == cellcount[i,1])
  df_metabolome <- rbind(df_metabolome, df_metabo)
  df_info <- rbind(df_info, cellcount[i,],cellcount[i,],cellcount[i,],cellcount[i,])
}


samplename <- paste0(df_info$class,"_", rep(c(1:4), 8))
col_names <- colnames(df_metabolome)
col_names <- gsub("\\.", " ", col_names)
colnames(df_metabolome) <- col_names

output_metabolome <- data.frame(Name = samplename) %>% cbind(df_metabolome[,-1])
output_metabolome$class <- gsub("control", "Control", output_metabolome$class)

```


```{r PCA for Figure 1c and S2a  }

xdata <- output_metabolome # Figure S2a: %>% filter(time== "1h") / filter(time== "24h") 
pca <- prcomp(xdata[,-c(1:4)], scale=TRUE)
score <- xdata[,c(1:4)]


score <- xdata[,c(1:4)] %>% cbind(pca$x)
group <- xdata %>% distinct(group, .keep_all = F) %>% .$group
order <- xdata %>% distinct(class, .keep_all = F) %>% .$class
score <- transform(score, class = factor(class, levels = order))
rotation <- pca$rotation %>% as.data.frame()
write.csv(rotation, "D:/2024/01_kanzo_multiomics/05_paper_work/Source data/2_metabolome/score_all.csv", row.names = TRUE)


col_class <- c("#424242", "#B71C1C", "#0D47A1","#00695C") #Figure 1c (1h)
col_class <- c("#F5F5F5", "#EDB7B9", "#B6CAE5","#8FC1BC") #Figure 1c (24h)

col_con <- colorRampPalette(c("#F5F5F5", "#424242")) # Figure S2a (all sample)
col_LPS <- colorRampPalette(c("#EDB7B9", "#B71C1C"))
col_ILG <- colorRampPalette(c("#B6CAE5", "#0D47A1"))
col_GU <- colorRampPalette(c("#8FC1BC", "#00695C"))


#Score plot
p <- ggplot(score, aes(x=PC1, y=PC2, fill = class))+
  geom_point(color = "black",size = 4, shape = 21)+ 
  labs(x = paste("PC1:",round(pca$sdev[1], digits=2),"%"),
       y = paste("PC2:",round(pca$sdev[2], digits=2),"%"))+

  scale_fill_manual(values = c(col_class), labels = group)+ #Fig.1c
  scale_fill_manual(values = c(col_con(2), col_LPS(2), col_ILG(2), col_GU(2)))+#Fig.S2a

  theme(panel.background = element_rect(fill = "transparent", colour = "transparent", size = 1),
        panel.grid = element_blank(),
        axis.text = element_text(size = 15, color = "black"), 
        axis.title = element_text(size = 18, colour = "black"),
        axis.line = element_line(colour="black"),
        legend.position = "bottom",
        # legend.position = "none",
        legend.key.size = unit(0.1, "cm"),
        legend.key = element_rect(fill = NA, color = NA), 
        legend.title = element_blank(),
        legend.text = element_text(size = 12),
        aspect.ratio = 1)

p

svg(paste0(path,"PCA_all.svg"), width = 5, height = 4)
print(p)
dev.off()


```


```{r Figure 2a}

library(circlize)
library(ComplexHeatmap)
path <- "D:/2024/01_kanzo_multiomics/01_HydrophilicMetabolomics/heatmap/"

# calculation of sample average
classes <- output_metabolome %>% distinct(class) %>% .$class
groups <- output_metabolome %>% distinct(group) %>% .$group
times <- output_metabolome %>% distinct(time) %>% .$time

output_metabo_average <- data.frame()
colname <- output_metabolome[,-c(1:4)] %>% colnames()

for (i in c(1:length(classes))) {
  # i=1
  tardf <- output_metabolome %>% filter(class == classes[[i]])
  tartime <- tardf %>% distinct(time) %>% .$time
  targroup <-  tardf %>% distinct(group) %>% .$group
  average <- tardf[,-c(1:4)] %>% colSums() %>% t() %>% as.data.frame()
  tarline <- data.frame(class = classes[[i]], group = targroup, time = tartime, average)
  output_metabo_average <- rbind(output_metabo_average, tarline)
  
}

colnames(output_metabo_average) <- c("class", "group", "time", metabolitename)
df_heat <- output_metabo_average %>%
  mutate(time = factor(time, levels = times)) %>% # times[-4] times
  arrange(time)


# Heat map for Figure 2a
hdata <- data.frame(df_heat[,-c(1:3)]) %>% scale() %>% as.data.frame()
rownames(hdata) <- df_heat$class
colnames(hdata) <- df_heat[,-c(1:3)] %>% colnames()

hdata_t<- t(hdata) %>% as.data.frame()
col_fun <- colorRamp2(c(min(hdata_t), 0, max(hdata_t)), c("navy", "white", "firebrick3"))
split_vector <- rep(1:2, each = 4, length.out = ncol(hdata_t))


# without ylabel (metabolite name)
heatmap <- Heatmap(hdata_t,
                   col = col_fun,
                   cluster_columns = FALSE,
                   column_split = split_vector,
                   border_gp = gpar(col = "black", lwd = 1),
                   show_row_names = FALSE,
                   show_column_names = TRUE,
                   column_title = NULL,
                   show_row_dend = FALSE,
                   column_gap = unit(0, "mm"), 
                   column_names_gp = gpar(fontsize = 8),
                   heatmap_legend_param = list(title = "Z-score", at = c(floor(min(hdata_t)), 0, ceiling(max(hdata_t)))),
                   width = unit(3, "cm"), height = unit(4, "cm")
)

heatmap

 svg(paste0(path, "heatmap_vs250320.svg"), width = 4, height = 6)
 plot(heatmap)
 dev.off()


# obtain row cluster number
ht_draw <- draw(heatmap)
dend <- row_dend(ht_draw)
clusters <- cutree(as.hclust(dend), h = 4)  
cluster_membership_df <- data.frame(Cluster = clusters)

cluster_colors <- setNames(viridis(7), unique(clusters))
row_anno <- rowAnnotation(Cluster = as.factor(clusters),
                          col = list(Cluster = cluster_colors),
                          show_legend = TRUE)


cluster2_rows <- cluster_membership_df %>% filter(Cluster == 2) %>% rownames() # cluster2; increased in GU group
cluster4_rows <- cluster_membership_df %>% filter(Cluster == 4) %>% rownames() # cluster4; increased following 1h after LPS stilmulation
filtered_matrix <- hdata_t[cluster4_rows, , drop = FALSE] 

# Heatmap (zoomed up)
final_heatmap <- Heatmap(filtered_matrix,
                   col = col_fun,
                   cluster_columns = FALSE,
                   column_split = split_vector,   
                   border_gp = gpar(col = "black", lwd = 1),
                   show_column_names = TRUE,
                   column_title = NULL,
                   show_row_dend = FALSE,
                   column_gap = unit(0, "mm"), 
                   column_names_gp = gpar(fontsize = 12),  
                   width = unit(3, "cm"), height = unit((10/length(cluster2_rows)*length(cluster4_rows)), "cm")
)
final_heatmap

 svg(paste0(path, "heatmap_cluster-4.svg"), width = 10, height = 10) #3.5
 plot(final_heatmap)
 dev.off()


```


```{r Significantly changed metabolites for Figure 2b}

# Tukey's HSD (vs. Control)
for (i in c(1:length(times))) {
  # i=1
  tardf <- output_metabolome %>% filter(time == times[i])
  
  for (j in c(5:ncol(tardf))) {
    # j=5
    testdf <- tardf[,c(1:4)] %>% mutate("Height" = tardf[,j])
    tukeydf <- TukeyHSD(aov(testdf$Height ~ testdf$group))
    result <- data.frame(pair = rownames(tukeydf$`testdf$group`), tukeydf$`testdf$group`)
    
    p_LPS <- filter(result, pair == "LPS-Control") %>% .$p.adj
    p_ILG <- filter(result, pair == "ILG-Control") %>% .$p.adj
    p_GU <- filter(result, pair == "GU-Control") %>% .$p.adj
    
    p <- rbind(p_LPS,p_ILG,p_GU) %>% as.data.frame()
    colnames(p) <- colnames(tardf)[j]
    

     if(j == 5){
      p_mtb <- data.frame("Name"= paste0(c("LPS", "ILG", "GU"), "_", times[i]),  "group"= groups[-1], "time" = times[i]) %>% cbind(p)
      } else{
      p_mtb <- cbind(p_mtb, p)
      }
  }
  
       if(i == 1){
      df_p_mtb <- p_mtb
      } else{
      df_p_mtb <- rbind(df_p_mtb, p_mtb)
      }
  
}


## Fold-change (vs. Control)
for (i in c(1:length(times))) {
  # i=1
  tardf <- output_metabolome %>% filter(time == times[i])
  
  for (j in c(5:ncol(tardf))) {
    # j=5
    testdf <- tardf[,c(1:4)] %>% mutate("Height" = tardf[,j])
    
    Mean_con <- filter(testdf, group == "Control")  %>% .$Height %>% mean()
    fc_LPS <- filter(testdf, group == "LPS") %>% .$Height %>% mean()/Mean_con
    fc_ILG <- filter(testdf, group == "ILG") %>% .$Height %>% mean()/Mean_con
    fc_GU <- filter(testdf, group == "GU") %>% .$Height %>% mean()/Mean_con
    
    fc <- data.frame(c(fc_LPS, fc_ILG, fc_GU))
    colnames(fc) <- colnames(tardf)[j]
    
     if(j == 5){
      fc_i <- data.frame("Name"= paste0(c("LPS", "ILG", "GU"), "_", times[i]), "group"= groups[-1],"time" = times[i]) %>%
      cbind(fc)
      } else{
      fc_i <- cbind(fc_i, fc)
      }
  }
  
       if(i == 1){
      df_fc <- fc_i
      } else{
      df_fc <- rbind(df_fc, fc_i)
      }
}

df_siginf <- data.frame(t(df_p_mtb[,-c(1:3)]), t(df_fc[,-c(1:3)]))
colnames(df_siginf) <- c(paste0("p_",df_p_mtb$Name), paste0("fc_",df_fc$Name))


for (i in c(1:6)) {
  # i=1
  tardf <- df_siginf[,c(i, (i+6))]
  lab <- ifelse(tardf[,1] < 0.05, ifelse(tardf[,2] > 2, "up", ifelse(tardf[, 2] < 0.5, "down", "x")),"xx")
  if (i==1) {
    df_siglab <- data.frame("Name"= rownames(df_siginf), lab)
  }else{
    df_siglab <- data.frame(df_siglab, lab)
  }
}

colnames(df_siglab) <- c("Name",df_p_mtb$Name)


# count of significantly changed metabolites
df_count_up <- data.frame(df_p_mtb[,c(1:3)], "count" = colSums(df_siglab[,-1] == "up"), type = "up") %>%
  transform(time = factor(time, levels = times), group = factor(group, levels = groups[-1]))
df_count_down <- data.frame(df_p_mtb[,c(1:3)], "count" = colSums(df_siglab[,-1] == "down"), type = "down") %>%
  transform(time = factor(time, levels = times), group = factor(group, levels = groups[-1]))
df_count <- rbind(df_count_up, df_count_down)

df_count <- transform(df_count, time = factor(time, levels = times), group = factor(group, levels = groups[-1]), type = factor(type, levels = c("up", "down"))) %>%
  mutate("lab" = paste0(group, "_",type))

# Figure 2b
   p <- ggplot(df_count_down, aes(x=time, y=count, fill = group))+
     geom_line(aes(group = group, col = group),size = 1)+
     scale_color_manual(values= c("#B71C1C", "#0D47A1","#00695C"))+
     geom_point(color = "black",size = 3, shape = 21)+
     scale_fill_manual(values= c("#B71C1C", "#0D47A1","#00695C"))+
     # scale_y_continuous(limits = c(80, 130))+
     labs(y = "Significantly changed metabolites")+
     theme(panel.background = element_rect(fill = "transparent", colour = "transparent", size = 1),
        panel.grid = element_blank(),
        axis.text = element_text(size = 12, color = "black"), 
        axis.title = element_text(size = 12, colour = "black"),
        axis.line = element_line(colour="black"),
        legend.position = "right",
        axis.title.x = element_blank(),
        legend.key.size = unit(0.1, "cm"),
        legend.key = element_rect(fill = NA, color = NA), 
        legend.title = element_blank(),
        legend.text = element_text(size = 12),
        aspect.ratio = 0.8)

 p
 
 filename <- paste0(path, "lineplot_down",".svg")
 svg(filename, width = 3.5, height = 2) #3.5
 print(p)
 dev.off()


```


```{r Figure 2c and 5c}
scientific_notation <- function(x) {
  x <- format(x, scientific = TRUE)
  x <- gsub("^(.*)e", "'\\1'e", x)
  x <- gsub("e", "%*%10^", x)
  x <- gsub('\\+', '', x)
  parse(text = x)
}

groups <- output_metabolome %>% distinct(group, .keep_all = F) %>% .$group
times <- output_metabolome %>% distinct(time, .keep_all = F) %>% .$time
col_groups <- c("#424242", "#B71C1C", "#0D47A1","#00695C")


linedata <- data.frame(output_metabolome[,c(1:4)], xtime = rep(c(rep(1,4),rep(24,4)), 4)) %>% cbind(output_metabolome[,-c(1:4)])
output_metabolome <- transform(output_metabolome, group = factor(group, levels = groups))
siglabel <- as.data.frame(lapply(df_p_mtb[,-c(1:3)], function(x) ifelse(x < 0.05, "*", "")))
sigdf <- data.frame(df_p_mtb[,c(1:3)],siglabel)
metabolitename <- colnames(df_p_mtb) %>% .[-c(1:3)]
colnames(sigdf) <- c(colnames(df_p_mtb[,c(1:3)]), metabolitename)
groups

sigdf_con <- data.frame(Name = c("Control_1h", "Control_24h"), time = c("1h", "24h"), group = c("Control", "Control"), lab = c("", ""))


for (i in c(6:ncol(linedata))) {
# i=6
  linedf <- linedata[,c(1:5,i)]
  linedf <- transform(linedf, group = factor(group, levels = groups))
  title <- colnames(linedf[6])
  sigdf_i <- data.frame(sigdf[,c(1:3)],"lab"=sigdf[,(i-2)])
  sigdf_i <- rbind(sigdf_con[1,], sigdf_i[c(1:3),], sigdf_con[2,], sigdf_i[c(4:6),])
  sigdf_i <- transform(sigdf_i, group = factor(group, levels = groups))
  
  linedf2 <-  linedf[order(linedf$time), ]
  class_2 <- linedf2 %>% distinct(class) %>% .$class
  linedf2 <- transform(linedf2, group = factor(group, levels = groups), class = factor(class, levels = class_2))
  
    p2 <- ggplot(linedf, aes(x = time, y = linedf[,6], fill = group, group = group)) +
    stat_summary(fun = "mean", geom = "bar", size = 0.5, alpha =0.5, position=position_dodge2(preserve = "single"), col = "black") +
      
    stat_summary(fun.max = function(x) mean(x) + std.error(x),
                 fun.min = function(x) mean(x) - std.error(x),
                 geom = 'errorbar', size =0.5, width =0.5, position = position_dodge(width = 0.9))+
      
    geom_jitter(size = 1.5, shape = 21 ,position = position_dodge(width = 0.9))+#, alpha =0.5
    scale_y_continuous(limits = c(0,max(linedf[,6])*1.1), expand = c(0,0), labels = scientific_notation)+
    scale_fill_manual(values = col_groups)+

    labs(y = "Peak height", title = title, x = "Time")+
    geom_text(data = sigdf_i, aes(x = time, y = max(linedf[,6]*0.1), label =  lab, group = group),
              position = position_dodge(width = 0.9), size = 8, color= "#ca0909")+

    theme(
      legend.position = "none",
      legend.title = element_blank(),
      panel.background = element_rect(fill = "transparent", colour = NA),
      axis.line = element_line(colour="black", linewidth = 1),
      axis.title.y = element_text(size = 12),
      axis.text.y = element_text(size = 10),
      panel.grid = element_blank(),
      plot.title = element_text(hjust = 0.5),
      axis.title.x = element_blank(),
      axis.text.x = element_text(size = 12, colour="black"),
      axis.ticks.x=element_blank(),
      aspect.ratio = 0.8
    )
  
 p2
 
   filename <- paste(title, "svg", sep=".")
  filepath <- paste("D:\\2024\\01_kanzo_multiomics\\01_HydrophilicMetabolomics\\DEM\\barplot",filename, sep="/")
  svg(filepath, width = 4, height = 2)
  print(p2)
  dev.off()

}

```


```{r Sorce fode for Figure 2f}
ibrary(pathview)
library(enrichplot)
library(ggplot2)
library(dplyr)
library(openxlsx)
library(tidyr)


GO_result <- read.xlsx("Source-data_hydrophipic metabolome.xlsx", 5) %>% mutate("-LogP.adj" = -log10(Holm.adjust)) %>% mutate("HitsRatio" = Hits/Total) %>% filter(Holm.adjust < 0.05) #Fig2f


heatmap_data_p <- GO_result %>%
  select(Pathway, Class,  "-LogP.adj") %>%
  tidyr::pivot_wider(names_from = Class, values_from = "-LogP.adj") %>%  as.data.frame()

rownames(heatmap_data_p) <- heatmap_data_p$Pathway
heatmap_data_p <- heatmap_data_p[,-1] 


annot.col <- data.frame("sig_count" = df_count$count[1:6], "class" = rep(groups[-1], 2))
rownames(annot.col) <- colnames(heatmap_data_p)
annotation_colors <- list("sig_count" = colorRampPalette(c("#fdefd5","#F2A71B"))(100),
                          "class" = c("LPS" = "#B71C1C", "ILG" = "#0D47A1", "GU" = "#00695C"))

hdata <- heatmap_data_p %>% as.matrix() 


new_mat <- apply(hdata, 2, function(col) {
  top5_indices <- order(col, decreasing = TRUE)[1:3]
  new_col <- rep("", length(col))
  new_col[top5_indices] <- "+"
  return(new_col)
})


new_mat <- matrix(new_mat, nrow = nrow(hdata), ncol = ncol(hdata))
rownames(new_mat) <- rownames(hdata)
colnames(new_mat) <- colnames(hdata)


library(pheatmap)
heatmap <- pheatmap(hdata,
                    color = colorRampPalette(c("#FAEFEF", "#BB0000"))(500),
                    annotation_col = annot.col,
                    annotation_colors = annotation_colors,
                    cellwidth = 10, cellheight = 10,
                    cluster_rows = F,cluster_cols = F,
                    show_colnames =T,show_rownames = T,
                    treeheight_row = F,treeheight_col = F,
                    border_color='gray40',
                    na_col = "transparent",
                    annotation_names_col = FALSE,
                    display_numbers = new_mat,
                    fontsize_row = 10, fontsize_col = 10)

heatmap

svg(paste0("Up_v3_top3.svg"), width =10, height = 5)
print(heatmap)
dev.off()
```
````
