## Supplementary material for "Unraveling anti-inflammatory metabolic signatures of *Glycyrrhiza uralensis* and isoliquiritigenin through multiomics": Source data: Source code for Figure 3 (lipidome).Rmd

````
---
title: "R Notebook"
output: html_notebook
---

```{r class summary}
library(pheatmap)
library(loadings)
library(tidyverse)
# library(ggfortify)
library(ggplot2)
library(caret)
library(tidyverse)
library(openxlsx)
library(dplyr)

setwd("D:/2024/01_kanzo_multiomics/05_paper_work/Source data/3_lipidmics")


lip <- read.xlsx("Source-data_lipidome_0327.xlsx",1) #Lipidome matrix for Fig3

df <- lip %>% .[-c(1:4),c(1:94)]
colnames(df) <- lip[4,c(1:94)]
class <- df %>% distinct(Ontology, .keep_all = FALSE) %>%.[,1]
sumdfs <- data.frame(Ontology = class)


for (i in c(35:ncol(df))) {
  
  classsums <- c()
  sample <- df[,c(1:34,i)]
  name <- colnames(df[i])
  
  for (j in c(1:length(class))) {
    tarclass <- sample %>% filter(Ontology == class[j]) %>% .[,35] %>% as.numeric()
    
    classsum <- ifelse(length(tarclass) > 1, sum(tarclass), tarclass)
    classsums <- c(classsums, classsum)
  }
  
  sumdf <- data.frame( name = classsums)
  sumdfs <- cbind(sumdfs, classsums)
  rm(sumdf)
  
}


colnames(sumdfs) <- c("Name",colnames(df[,c(35:ncol(df))]))
outputdf <- data.frame(Name = colnames(sumdfs),t(sumdfs))
colnames(outputdf) <- outputdf[1,]
outputdf <- outputdf %>% .[-1,]
order <- read.xlsx("Source-data_lipidome_0327.xlsx",4) %>% .$Name #Fig3d and S3a

outputdf$Name <- factor(outputdf$Name, levels = order)
outputdf <- outputdf[order(outputdf$Name), ]
NameInfo <- read.xlsx("Source-data_lipidome_0327.xlsx",4) %>% .[,c(2:4)]
outputdf <- data.frame(Name = outputdf[,1], NameInfo, lapply(outputdf[,-1], as.numeric)) ##Fig3d and S3a

```


```{r cell counts correction}
library(exactRankTests)
library(ggplot2)
library(dplyr)
library(openxlsx)
library(ggsignif)
library(plotrix)
library(loadings)
library(tidyverse)
library(pheatmap)
library(plotrix)

cellcount <- read.xlsx("Source-data_lipidome_0327.xlsx", 2) %>% #Sample metadata for Fig3
  mutate_all(~ gsub("control", "Control", ., ignore.case = TRUE)) %>% mutate(across(-c(1:3), as.numeric))

lipid_metabolome <-  read.xlsx("Source-data_lipidome_0327.xlsx", 3) %>% #Fig 3a, 3b, 3f, and S3d
  mutate_all(~ gsub("control", "Control", ., ignore.case = TRUE)) %>% mutate(across(-c(1:4), as.numeric))
lipid_class <- read.xlsx("Source-data_lipidome_0327.xlsx",4)%>%
  mutate_all(~ gsub("control", "Control", ., ignore.case = TRUE)) %>% mutate(across(-c(1:4), as.numeric))


df_metabolome <- data.frame()
df_lipidclass <- data.frame()
df_info <- data.frame()

for (i in c(1:nrow(cellcount))) {
  df_metabo <- lipid_metabolome %>% filter(class == cellcount[i,1]) %>% .[,-c(1:4)]/cellcount[i,4]
  df_class <- lipid_class %>% filter(class == cellcount[i,1]) %>% .[,-c(1:4)]/cellcount[i,4]
  
  df_metabolome <- rbind(df_metabolome, df_metabo) 
  df_lipidclass <- rbind(df_lipidclass, df_class)
  df_info <- rbind(df_info, cellcount[i,],cellcount[i,],cellcount[i,])
}

samplename <- paste0(df_info$class,"_", rep(c(1:3), 20)) 

##lipid (metabolites)
col_names <- colnames(df_metabolome)
col_names <- gsub("\\.", " ", col_names)
colnames(df_metabolome) <- col_names
output_metabolome <- data.frame(Name = samplename, df_info[,-4]) %>% cbind(df_metabolome)

##lipid (class)
col_names <- colnames(df_lipidclass)
col_names <- gsub("\\.", " ", col_names)
colnames(df_lipidclass) <- col_names
output_class <- data.frame(Name = samplename, df_info[,-4]) %>% cbind(df_lipidclass)

```


```{r Figure 3a}

classes <- output_class %>% distinct(class) %>% .$class

output_class_average <- data.frame()
colname <- output_class[,-c(1:4)] %>% colnames()

for (i in c(1:length(classes))) {
  # i=1
  tardf <- output_class %>% filter(class == classes[[i]])
  tartime <- tardf %>% distinct(time) %>% .$time
  targroup <-  tardf %>% distinct(group) %>% .$group
  average <- tardf[,-c(1:4)] %>% colSums() %>% t() %>% as.data.frame()
  tarline <- data.frame(class = classes[[i]], group = targroup, time = tartime, average)
  output_class_average <- rbind(output_class_average, tarline)
  
}


classes <- output_metabolome %>% distinct(class) %>% .$class
output_lipid_average <- data.frame()
colname <- output_metabolome[,-c(1:4)] %>% colnames()

for (i in c(1:length(classes))) {
  # i=1
  tardf <- output_metabolome %>% filter(class == classes[[i]])
  tartime <- tardf %>% distinct(time) %>% .$time
  targroup <-  tardf %>% distinct(group) %>% .$group
  average <- tardf[,-c(1:4)] %>% colSums() %>% t() %>% as.data.frame()
  tarline <- data.frame(class = classes[[i]], group = targroup, time = tartime, average)
  output_lipid_average <- rbind(output_lipid_average, tarline)
  
}

colnames(output_lipid_average) <- c("class", "group", "time", colname)


library(circlize)
library(ComplexHeatmap)
path <- "D:/2024/01_kanzo_multiomics/02_Lipidomics/20241107/heatmap/"


# Figure 3a
df_heat <- output_lipid_average %>%
  mutate(time = factor(time, levels = times[-4])) %>% arrange(time)

hdata <- data.frame(df_heat[,-c(1:3)]) %>% scale() %>% as.data.frame()
rownames(hdata) <- df_heat$class
colnames(hdata) <- df_heat[,-c(1:3)] %>% colnames()

hdata_t<- t(hdata) %>% as.data.frame()
col_fun <- colorRamp2(c(min(hdata_t), 0, max(hdata_t)), c("navy", "white", "firebrick3"))

split_vector <- rep(1:5, each = 4, length.out = ncol(hdata_t))

heatmap <- Heatmap(hdata_t,
                   col = col_fun,
                   cluster_columns = FALSE,
                   column_split = split_vector,   
                   border_gp = gpar(col = "black", lwd = 1),
                   show_row_names = FALSE,
                   show_column_names = TRUE,
                   column_title = NULL,
                   show_row_dend = FALSE,
                   column_gap = unit(0, "mm"), 
                   column_names_gp = gpar(fontsize = 8),
                   heatmap_legend_param = list(title = "Z-score", at = c(floor(min(hdata_t)), 0, ceiling(max(hdata_t)))),
                   width = unit(6, "cm"), height = unit(8, "cm")
)

heatmap


 svg(paste0(path, "heatmap_vs250320.svg"), width = 4, height = 6) #3.5
 plot(heatmap)
 dev.off()
 

# obtain row cluster number
ht_draw <- draw(heatmap)
dend <- row_dend(ht_draw)
clusters <- cutree(as.hclust(dend), h = 7)
cluster_membership_df <- data.frame(Cluster = clusters)
unique(clusters)
cluster_colors <- setNames(viridis(6), unique(clusters))
row_anno <- rowAnnotation(Cluster = as.factor(clusters),
                          col = list(Cluster = cluster_colors),
                          show_legend = TRUE)
 
cluster1_rows <- cluster_membership_df %>% filter(Cluster == 1) %>% rownames() # cluster1; increased in LPS group following 24h after LPS stimuli.
filtered_matrix <- hdata_t[cluster1_rows, , drop = FALSE]


# Figure 3a (zoomed up)
 heatmap <- Heatmap(filtered_matrix,
                   col = col_fun,
                   cluster_columns = FALSE,
                   column_split = split_vector,  
                   border_gp = gpar(col = "black", lwd = 1),
                   show_column_names = TRUE,
                   column_title = NULL,
                   show_row_dend = FALSE,
                   column_gap = unit(0, "mm"), 
                   column_names_gp = gpar(fontsize = 12), 
                   heatmap_legend_param = list(title = "Z-score", at = c(floor(min(hdata_t)), 0, ceiling(max(hdata_t)))),
                   width = unit(8, "cm"), height = unit(50, "cm")
)

heatmap


 svg(paste0(path, "heatmap_250320_Cluster1.svg"), width = 10, height = 30)
 plot(heatmap)
 dev.off()

```


```{r Figure 1c and 3b}


# Figure 1c

times <- df_info %>% distinct(time) %>% .$time

for (i in c(1:length(times))) {
 # i=1
  tartime <- times[i]
  xdata <- output_metabolome %>% filter(time== tartime)
  pca <- prcomp(xdata[,-c(1:4)], scale=TRUE)
  score <- xdata[,c(1:4)] %>% cbind(pca$x)
  
  rotation <- data.frame(Name = colnames(df_metabolome), pca$rotation)
  path <- c("D:\\2024\\01_kanzo_multiomics\\02_Lipidomics\\20241107\\PCA\\")
  write.xlsx(rotation, paste0(path, "rotation_", tartime, ".xlsx"), rowNames = F)
  
  
  col_class <- c("#424242", "#B71C1C", "#0D47A1","#00695C")
  order <- xdata %>% distinct(class, .keep_all = F) %>% .$class
  score <- transform(score, class = factor(class, levels = order))
  
  p <- ggplot(score, aes(x=PC1, y=PC2, fill = class))+
  geom_point(color = "black",size = 5, shape = 21)+
  labs(x = paste("PC1:",round(pca$sdev[1], digits=2),"%"),
       y = paste("PC2:",round(pca$sdev[2], digits=2),"%"))+
      scale_fill_manual(values = c(col_class)
                    )+
  theme(panel.background = element_rect(fill = "transparent", colour = "black", size = 1),
        panel.grid = element_blank(),
        axis.text = element_text(size = 15, color = "black"), 
        axis.title = element_text(size = 18, colour = "black"),
        axis.line = element_line(colour="transparent"),
        legend.position = "bottom",
        legend.key.size = unit(0.1, "cm"),
        legend.key = element_rect(fill = NA, color = NA), 
        legend.title = element_blank(),
        legend.text = element_text(size = 12),
        aspect.ratio = 1)
  
  p
  
  svg(paste0(path, "ScorePlot_", tartime, ".svg"), width = 3, height = 3) #each time
  print(p)
  dev.off()
  
}


# Figure 3b
control <- colorRampPalette(c("#F5F5F5", "#424242"))
LPS <- colorRampPalette(c("#F5F5F5", "#B71C1C"))
ILG <- colorRampPalette(c("#F5F5F5", "#0D47A1"))
GU <- colorRampPalette(c("#F5F5F5", "#00695C"))
 
groups <- df_info %>% distinct(group) %>% .$group
times <- df_info %>% distinct(time) %>% .$time

num <- 5
col_con <- control(num)
col_LPS <- LPS(num)
col_ILG <- ILG(num)
col_GU <- GU(num)
pal <- list(col_con, col_LPS, col_ILG, col_GU)
 
path <- "D:/2024/01_kanzo_multiomics/02_Lipidomics/20241107/PCA_time-couse"


for (i in c(1:length(groups))) {
  # i=1
  xdata <- output_metabolome %>% filter(group== groups[i]) 
  pca <- prcomp(xdata[,-c(1:4)], scale=TRUE)
  
  score <- xdata[,c(1:4)] %>% cbind(pca$x)
  order <- xdata %>% distinct(class, .keep_all = F) %>% .$class
  score <- transform(score, class = factor(class, levels = order))
  rotation <- data.frame(pca$rotation)
  
  filename <- paste0(path,"/rotation_", groups[i], "_v2.xlsx")
  write.xlsx(rotation,filename ,rowNames = TRUE)

  p <- ggplot(score, aes(x=PC1, y=PC2, fill = class))+
  geom_point(color = "black",size = 5, shape = 21)+
  labs(x = paste("PC1:",round(pca$sdev[1], digits=2),"%"),
       y = paste("PC2:",round(pca$sdev[2], digits=2),"%"))+
  scale_fill_manual(values = pal[[i]])+
  theme(panel.background = element_rect(fill = "transparent", colour = "transparent", size = 1),
        panel.grid = element_blank(),
        axis.text = element_text(size = 15, color = "black"), 
        axis.title = element_text(size = 18, colour = "black"),
        axis.line = element_line(colour="black"),
        legend.position = "bottom",
        legend.key.size = unit(0.1, "cm"),
        legend.key = element_rect(fill = NA, color = NA), 
        legend.title = element_blank(),
        legend.text = element_text(size = 12),
        aspect.ratio = 1)

 p
 
 filename <- paste0(path, "/pca_1vs2_", groups[i], "_legend.svg")
 svg(filename, width = 5, height = 3) 
 print(p)
 dev.off()

}
 

```


```{r Figure 3c and 3e}
library(plotly)

metabolome_ontology <- read.xlsx("Source-data_lipidome_0327.xlsx", 1) %>% .[-c(1:4),12] #Lipidome matrix for Fig3
times <- output_metabolome %>% distinct(time) %>% .$time
groups <- output_metabolome %>% distinct(group) %>% .$group
classes <- output_metabolome %>% distinct(class) %>% .$class

# Tukey'HSD
for (i in c(1:length(times))) {
  # i=1
  tardf <- output_metabolome %>% filter(time == times[i])
  
  for (j in c(5:ncol(tardf))) {
    # j=5
    testdf <- tardf[,c(1:4)] %>% mutate("Height" = tardf[,j])
    tukeydf <- TukeyHSD(aov(testdf$Height ~ testdf$group))
    result <- data.frame(pair = rownames(tukeydf$`testdf$group`), tukeydf$`testdf$group`)
    
    p_LPS <- filter(result, pair == "LPS-Control") %>% .$p.adj
    p_ILG <- filter(result, pair == "ILG-Control") %>% .$p.adj
    p_GU <- filter(result, pair == "GU-Control") %>% .$p.adj
    
    p <- rbind(p_LPS,p_ILG,p_GU) %>% as.data.frame()
    colnames(p) <- colnames(tardf)[j]
    

     if(j == 5){
      p_ont <- data.frame("Name"= paste0(c("LPS", "ILG", "GU"), "_", times[i]),  "group"= groups[-1], "time" = times[i]) %>% cbind(p)
      } else{
      p_ont <- cbind(p_ont, p)
      }
  }
  
       if(i == 1){
      df_p_ont <- p_ont
      } else{
      df_p_ont <- rbind(df_p_ont, p_ont)
      }
  
}


# Fold-change (vs.Control)
for (i in c(1:length(times))) {
  # i=1
  tardf <- output_metabolome %>% filter(time == times[i])
  
  for (j in c(5:ncol(tardf))) {
    # j=5
    testdf <- tardf[,c(1:4)] %>% mutate("Height" = tardf[,j])
    
    Mean_con <- filter(testdf, group == "Control")  %>% .$Height %>% mean()
    fc_LPS <- filter(testdf, group == "LPS") %>% .$Height %>% mean()/Mean_con
    fc_ILG <- filter(testdf, group == "ILG") %>% .$Height %>% mean()
    fc_GU <- filter(testdf, group == "GU") %>% .$Height %>% mean()
    
    fc <- data.frame(c(fc_LPS, fc_ILG, fc_GU))
    colnames(fc) <- colnames(tardf)[j]
    
     if(j == 5){
      fc_i <- data.frame("Name"= paste0(c("LPS", "ILG", "GU"), "_", times[i]), "group"= groups[-1],"time" = times[i]) %>%
      cbind(fc)
      } else{
      fc_i <- cbind(fc_i, fc)
      }
  }
  
       if(i == 1){
      df_fc <- fc_i
      } else{
      df_fc <- rbind(df_fc, fc_i)
      }
  
}


col_names <- colnames(df_metabolome)
col_names <- gsub("\\.", " ", col_names)
colnames(df_fc)<- c("Name","group", "time", col_names)
colnames(df_p_ont)<- c("Name", "group","time", col_names)


# Figure 3c
library(stringr)
df_siginf <- data.frame(t(df_p_ont[,-c(1:3)]), t(df_fc[,-c(1:3)]))
colnames(df_siginf) <- c(paste0("p_",df_p_ont$Name), paste0("fc_",df_fc$Name))

for (i in c(1:15)) {
  # i=1
  tardf <- df_siginf[,c(i, (i+15))]
  lab <- ifelse(tardf[,1] < 0.05, ifelse(tardf[,2] > 2, "up", ifelse(tardf[, 2] < 0.5, "down", "x")),"xx")
  if (i==1) {
    df_siglab <- data.frame("Name"= rownames(df_siginf), lab)
  }else{
    df_siglab <- data.frame(df_siglab, lab)
  }
}

colnames(df_siglab) <- c("Name",df_p_ont$Name)


df_count_up <- data.frame(df_p_ont[,c(1:3)], "count" = colSums(df_siglab[,-1] == "up"), type = "up") %>% filter( time != "4h") %>% transform(time = factor(time, levels = times[-4]), group = factor(group, levels = groups[-1]))
df_count_down <- data.frame(df_p_ont[,c(1:3)], "count" = colSums(df_siglab[,-1] == "down"), type = "down") %>% filter( time != "4h") %>% transform(time = factor(time, levels = times[-4]), group = factor(group, levels = groups[-1]))
df_count <- rbind(df_count_up, df_count_down)

df_count <- transform(df_count, time = factor(time, levels = times), group = factor(group, levels = groups[-1]), type = factor(type, levels = c("up", "down"))) %>%
  mutate("lab" = paste0(group, "_",type))

df_count <- df_count %>% filter( time != "4h") %>% filter(type == "down")

  p <- ggplot(df_count_down, aes(x= time, y= count, fill = group))+
    geom_line(aes(group = group, col = group),size = 1)+
    scale_color_manual(values= c("#B71C1C", "#0D47A1","#00695C"))+
    geom_point(shape = 21, color = "black",size = 3)+
    scale_fill_manual(values= c("#B71C1C", "#0D47A1","#00695C"))+
    labs(y = "Significantly changed lipids")+
    theme(panel.background = element_rect(fill = "transparent", colour = "transparent", size = 1),
        panel.grid = element_blank(),
        axis.text = element_text(size = 12, color = "black"), 
        axis.title = element_text(size = 12, colour = "black"),
        axis.line = element_line(colour="black"),
        legend.position = "right",
        axis.title.x = element_blank(),
        legend.key.size = unit(0.1, "cm"),
        legend.key = element_rect(fill = NA, color = NA), 
        legend.title = element_blank(),
        legend.text = element_text(size = 12),
        aspect.ratio = 0.8)

 p


 
 path <- "D:/2024/01_kanzo_multiomics/05_paper_work/Source data/3_lipidmics/"
 # filename <- paste0(path, "lineplot_up_v2",".svg")
 filename <- paste0(path, "lineplot_down_v2",".svg")
 svg(filename, width = 3.5, height = 2)
 print(p)
 dev.off()
 
 

# Figure 3e
path <- "D:/2024/01_kanzo_multiomics/02_Lipidomics/20241107/FC_dotplot_2/"
colorpal <- c("white","#FAAB00", "#FAEC00","#2b8cbe", "#08519c", "#a50f15", "#810f7c","#99d8c9","#006d2c","grey40")

for (i in c(1:length(times))) {
  # i=2
  FC_i <- df_fc %>% filter(time == times[i]) %>% .[,-c(1:3)] %>% t() %>% data.frame()
  p_i <- df_p_ont %>% filter(time == times[i]) %>% .[,-c(1:3)] %>% t() %>% data.frame()
  df_plot <- data.frame("Name" = row.names(FC_i), "Ontology" = metabolome_ontology, FC_i, p_i)
  colnames(df_plot) <- c("Name","Ontology", paste0(groups[-1],"_FC"), paste0(groups[-1],"_p"))
  
  df_plot$lab <- ifelse(df_plot$LPS_p < 0.05 & df_plot$LPS_FC >= 2, "up", 
                 ifelse(df_plot$LPS_p < 0.05 & df_plot$LPS_FC <= 0.5, "down", "n.s."))
  df_plot <- transform(df_plot, lab = factor(lab, levels = c("up", "down", "n.s.")))
  

  df_plot <- df_plot  %>%
    mutate(
    ont_lab = case_when(
      Ontology == "TG" ~ "TG",
      Ontology == "DG" ~ "DG",
      Ontology == "CE" ~ "CE",
      str_detect(Ontology, "HexCer") ~ "HexCer",
      Ontology == "Hex2Cer" ~ "Hex2Cer",
      Ontology %in% c("Cer_AP", "Cer_NDS", "Cer_NS") ~ "Ceramide",
       Ontology %in% c("PA", "PC", "PE", "PG", "PI", "PS") ~ "PA/PC/PE/PG/PI/PS",
      str_detect(Ontology, "Ether") ~ "Ether lipid",
      Ontology %in% c("LPE", "LPC") ~ "LysoPhospholipid",
      TRUE ~ "Others"
    ))
  
  col_order <- c("PA/PC/PE/PG/PI/PS",  "LysoPhospholipid", "Ether lipid","DG","TG","CE", "Ceramide", "HexCer", "Hex2Cer","Others")
  df_plot <- transform(df_plot, ont_lab = factor(ont_lab, levels = col_order))
  df_plot <- df_plot %>%
  mutate(order = factor(ont_lab, levels = rev(c(col_order)))) %>%
  arrange(order)
 
  
  g <- ggplot(df_plot, aes(x= log2(ILG_FC), y= log2(GU_FC)))+
    geom_point(aes(fill = ont_lab, shape = lab),color = "black",size = 2)+
    scale_shape_manual(values = c("up" = 24, "down"= 25, "n.s."= 21))+
    labs(x = "Log2 Fold-change (ILG vs.Control)",
         y = "Log2 Fold-change (GU vs.Control)")+

    scale_fill_manual(values = colorpal)+
    geom_vline(xintercept = 0, linetype="dashed")+
    geom_hline(yintercept = 0, linetype="dashed")+
    theme(panel.background = element_rect(fill = "transparent", colour = "black", size = 1),
          panel.grid = element_blank(),
          axis.text = element_text(size = 15, color = "black"), 
          axis.title = element_text(size = 18, colour = "black"),
          axis.line = element_line(colour="transparent"),
          legend.position = "bottom",
          legend.key.size = unit(0.1, "cm"),
          legend.key = element_rect(fill = NA, color = NA), 
          legend.title = element_blank(),
          legend.text = element_text(size = 12),
          aspect.ratio = 1)
  
  g
  
  
  
  
  filename <- paste0(path, "ILG-GU(vs.Control)_", times[i], "_v2.svg")
  svg(filename, width = 5, height = 5.5)
  print(g)
  dev.off()
  
  
  # plotly
  # library(plotly)
#   df_plotly <- df_plot %>%
#   mutate(
#     shape = case_when(
#       lab == "up" ~ "triangle-up",
#       lab == "down" ~ "triangle-down",
#       TRUE ~ "circle"
#     )
#   )
# 
# 
# p <- plot_ly(
#   data = df_plotly,
#   x = ~log2(ILG_FC),
#   y = ~log2(GU_FC),
#   type = 'scatter',
#   mode = 'markers',
#   text = ~Name,
#   hoverinfo = 'text',
#   color = ~ont_lab, 
#   colors = colorpal,
#   marker = list(
#     size = 10,
#     line = list(color = 'black', width = 1)
#   )
# ) %>%
#   add_trace(
#     marker = list(
#       symbol = ~shape
#     )
#   ) %>%
#   layout(
#     xaxis = list(title = "Log2 Fold-change (ILG vs.Control)"),
#     yaxis = list(title = "Log2 Fold-change (GU vs.Control)"),
#     shapes = list(
#       list(type = "line", x0 = 0, x1 = 0, y0 = -Inf, y1 = Inf, line = list(dash = 'dash')), 
#       list(type = "line", x0 = -Inf, x1 = Inf, y0 = 0, y1 = 0, line = list(dash = 'dash'))  
#     ),
#     legend = list(orientation = 'h', x = 0.5, xanchor = 'center', y = -0.2),
#     margin = list(l = 80, r = 80, t = 50, b = 80),
#     showlegend = TRUE
#   )
# 
# # Save as html files
# htmlwidgets::saveWidget(p, paste0(path, "ILG-GU(vs.Control)_", times[i], "scatterplot.html"), selfcontained = TRUE)


}


```


```{r Figure 3d and S3b}
library(plotrix)
scientific_notation <- function(x) {
  x <- format(x, scientific = TRUE)
  x <- gsub("^(.*)e", "'\\1'e", x)
  x <- gsub("e", "%*%10^", x)
  x <- gsub('\\+', '', x)
  parse(text = x)
}

groups <- output_class %>% distinct(group, .keep_all = F) %>% .$group
times <- output_class %>% distinct(time, .keep_all = F) %>% .$time
col_groups <- c("#424242", "#B71C1C", "#0D47A1","#00695C")

linedata <- output_class %>% filter(time %in% c("1h", "24h"))
linedata <- data.frame(linedata[,c(1:4)], xtime = rep(c(rep(1,3),rep(24,3)), 4)) %>% cbind(linedata[,-c(1:4)]) %>%
  transform(time = factor(time, levels = c("1h", "24h")), group = factor(group, levels = groups))

siglabel <- as.data.frame(lapply(df_p_ont[,-c(1:3)], function(x) ifelse(x < 0.05, "*", "")))
sigdf <- data.frame(df_p_ont[,c(1:3)],siglabel)
metabolitename <- colnames(df_p_ont) %>% .[-c(1:3)]
colnames(sigdf) <- c(colnames(df_p_ont[,c(1:3)]), metabolitename)
groups

sigdf_con <- data.frame(Name = c("Control_1h", "Control_24h"), time = c("1h", "24h"), group = c("Control", "Control"), lab = c("", ""))


for (i in c(6:ncol(linedata))) {
# i=6
  linedf <- linedata[,c(1:5,i)]
  linedf <- transform(linedf, group = factor(group, levels = groups))
  title <- colnames(linedf[6])
  sigdf_i <- data.frame(sigdf %>% filter(time %in% c("1h", "24h")) %>% .[,c(1:3)],
                        "lab"= sigdf %>% filter(time %in% c("1h", "24h")) %>% .[,(i-2)])
  sigdf_i <- rbind(sigdf_con[1,], sigdf_i[c(1:3),], sigdf_con[2,], sigdf_i[c(4:6),])
  sigdf_i <- transform(sigdf_i, group = factor(group, levels = groups))
  
  linedf2 <-  linedf[order(linedf$time), ]
  linedf2 <- transform(linedf2, group = factor(group, levels = groups), time = factor(time, levels = c("1h", "24h")))
  
    p <- ggplot(linedf2, aes(x = time, y = linedf2[,6], fill = group, group = group)) +
    stat_summary(fun = "mean", geom = "bar", size = 0.5, alpha =0.5, position=position_dodge2(preserve = "single"), col = "black") +
      
    stat_summary(fun.max = function(x) mean(x) + std.error(x),
                 fun.min = function(x) mean(x) - std.error(x),
                 geom = 'errorbar', size =0.5, width =0.5, position = position_dodge(width = 0.9))+
      
    geom_jitter(size = 1.5, shape = 21 ,position = position_dodge(width = 0.9))+#, alpha =0.5
    scale_y_continuous(limits = c(0,max(linedf2[,6])*1.1), expand = c(0,0), labels = scientific_notation)+
    scale_fill_manual(values = col_groups)+

    labs(y = "Abundance", title = title, x = "Time")+
    geom_text(data = sigdf_i, aes(x = time, y = max(linedf2[,6]*0.1), label =  lab, group = group),
              position = position_dodge(width = 0.9), size = 8, color= "#ca0909")+

    theme(
      legend.position = "none",
      legend.title = element_blank(),
      panel.background = element_rect(fill = "transparent", colour = NA),
      axis.line = element_line(colour="black", linewidth = 1),
      axis.title.y = element_text(size = 12),
      axis.text.y = element_text(size = 10),
      panel.grid = element_blank(),
      plot.title = element_text(hjust = 0.5),
      axis.title.x = element_blank(),
      axis.text.x = element_text(size = 12, colour="black"),
      axis.ticks.x=element_blank(),
      aspect.ratio = 0.8
    )
  
 p
 
   filename <- paste(title, "svg", sep=".")
  filepath <- paste(path, "barplot",filename, sep="/")
  svg(filepath, width = 4, height = 2)
  print(p)
  dev.off()

}


```


```{r Figure 3f and S3d}


metabolome_ontology <- metabolome_ontology <- read.xlsx("Source-data_lipidome_0327.xlsx", 1) %>% .[-c(1:4),12] #Lipidome matrix for Fig3
heatdf <- output_lipid_average %>% filter(time == "24h")

t_metabo <- data.frame(Ontology = metabolome_ontology, t(heatdf[,-c(1:3)]))
colnames(t_metabo) <- c("Ontology",heatdf$class)

ontology <- metabolome_ontology %>% unique()


for (i in c(1:length(ontology))) {
  # i=1
  tardf <- t_metabo %>% filter(Ontology == ontology[i]) %>% select(-Ontology)
  t_tardf <- t(tardf) %>% scale() %>% as.data.frame()
  test <- t(t_tardf) %>% as.data.frame()
  
  annot <- data.frame(abundance =  colSums(t_tardf))
  
  
  row_clustering <- if (nrow(test) > 1) TRUE else FALSE
  heatmap <- pheatmap::pheatmap(test,
                              col = colorRampPalette(c("navy","white", "firebrick3"))(500),
                              # breaks = seq(-5, 5, length.out = 501),
                              breaks = c(seq(min(test)-0.5, 0, length.out=250), seq(0.01, max(test)+0.5, length.out=251)),
                              #color = viridis::viridis(256),
                              annotation_row = annot,
                              annotation_names_row = F,
                              # annotation_colors = annot_col,
                              # annotation_colors = anno_color,
                              border_color='gray20',
                              # na_col = "grey",
                              treeheight_row = 0,
                              cellheight = 10,  cellwidth = 10,
                              show_colnames = T,
                              show_rownames = T,
                              cluster_cols = F,
                              cluster_rows = row_clustering
                              )


heatmap

  filename <- paste(ontology[i], "svg", sep=".")
  filepath <- paste("D:/2024/01_kanzo_multiomics/02_Lipidomics/20240708/heatmap/lipid_24h",filename, sep="/")
  svg(filepath, width = 10)
  print(heatmap)
  dev.off()
}

```
````
