## Supplementary material for "Unraveling anti-inflammatory metabolic signatures of *Glycyrrhiza uralensis* and isoliquiritigenin through multiomics": Source data: Source code for Figure 4 (phosphoproteome).Rmd

````
---
title: "R Notebook"
output: html_notebook
---

```{r anova}

library(dplyr)
library(exactRankTests)
library(ggplot2)
library(dplyr)
library(openxlsx)
library(ggsignif)
library(plotrix)
library(loadings)
library(tidyverse)
library(pheatmap)
library(plotrix)

scientific_notation <- function(x) {
  x <- format(x, scientific = TRUE)
  x <- gsub("^(.*)e", "'\\1'e", x)
  x <- gsub("e", "%*%10^", x)
  x <- gsub('\\+', '', x)
  parse(text = x)
}
setwd("D:/2024/01_kanzo_multiomics/05_paper_work/Source data/4_phosphoproteome")
# path <- "D:\\2024\\01_kanzo_multiomics\\03_Phosphoproteomics\\"
df_log <- read.xlsx("Source-data_phosphoproteome.xlsx",1) # Log2 normalized matrix for Fig4
df_raw <- read.xlsx("Source-data_phosphoproteome.xlsx",2) # Raw intensity matrix for Fig4


times <- df_log %>% distinct(time)
Index <- colnames(df_log[,-c(1:4)])
Prs_df <- data.frame("Index" = Index)


for (j in c(1:4)) {
  # j=1
  tartime <- times$time[j]
  Prs <- list()
  df_anova <- df_log %>% dplyr::filter(time == tartime)
  
  for (i in c(5:ncol(df_anova))) {
  # i=5
  result <- anova(aov(df_anova[,i]~class, data=df_anova))
  Prs[i-4]<- result$`Pr(>F)` %>% .[[1]]
  
  }
  Pr <- unlist(Prs)
  Prs_df <- cbind(Prs_df, Pr)
}


colnames(Prs_df) <- c("Index", paste0("P_",times$time))
sig_index <- Prs_df %>% dplyr::filter(P_0h < 0.05 | P_15m < 0.05 | P_2h < 0.05 | P_24h < 0.05)

df_log_sig <- df_log %>% dplyr::select(1:4, sig_index$Index)

times <- df_log_sig %>% distinct(time) %>% .$time
classes <- df_log_sig %>% distinct(class) %>% .$class
groups <- df_log_sig %>% distinct(group) %>% .$group


```


```{r Figure 2d, 2e, S3a, S3c,and 5d}

raw_index <- intersect(colnames(df_raw), sig_index$Index)
df_raw_sig <- df_raw %>% select(1:4, all_of(raw_index))

linedata <- data.frame(df_raw[,c(1:4)], xtime = rep(c(rep(0,3),rep(1,3), rep(2,3), rep(3,3)), 4)) %>% cbind(df_raw[,-c(1:4)])
groups <- linedata %>% distinct(group) %>% .$group
times <-  linedata %>% distinct(time) %>% .$time
col_class <- c("#424242", "#B71C1C", "#0D47A1","#00695C")

for (i in c(6:ncol(linedata))) {
# i=6
  linedf <- linedata[,c(1:5,i)]
  linedf <- transform(linedf, group = factor(group, levels = groups))
  title <- colnames(linedf[6])
  

  p <- ggplot(linedf, aes(x = xtime, y = linedf[,6], col = group, group = group)) +
    stat_summary(fun = "mean", geom = "line", size = 1) +
    geom_jitter(size = 0.75, shape = 16, width = .1, alpha =0.5)+ #svg: size = 0.5,, png: size = 1
    stat_summary(fun = "mean", geom = "point", size = 2.5, shape = 21, fill = "white") +
    scale_y_continuous(limits = c(0,max(linedf[,6])*1.1), expand = c(0,0), labels = scientific_notation)+
    scale_x_continuous(breaks=c(0,3), labels =c("0h", "24h"))+
    scale_color_manual(values = col_class)+
    labs(y = "Peak height", title = title, x = "Time")+
    theme(
      legend.position = "none",
      legend.title = element_blank(),
      axis.line = element_line(colour="black", size = 0.75),
      panel.background = element_rect(fill = "transparent", colour = NA),,
      axis.title = element_text(size = 7.5, colour="gray20"),
      axis.text.y = element_text(size = 7.5, colour="gray20"),
      axis.text.x = element_text(size = 10, colour="gray20"),
      plot.title = element_text(size = 10, colour="gray20",hjust = 0.5),
      panel.grid = element_blank(),
      axis.ticks.x=element_blank(), 
      aspect.ratio = 0.8
    )
  
 p

  filename <- paste(title, "svg", sep=".")
  filepath <- paste("D:/2024/01_kanzo_multiomics/03_Phosphoproteomics/Lineplot/v5/",filename, sep="/")
  svg(filepath, width = 3, height = 2)
  print(p)
  dev.off()

}


```


```{r Figure 4a}
library(circlize)
library(ComplexHeatmap)

for (i in c(1:length(classes))) {
  # i=1
  tardf <- df_log_sig %>% filter(class == classes[[i]])
  tartime <- tardf %>% distinct(time) %>% .$time
  targroup <-  tardf %>% distinct(group) %>% .$group
  average <- tardf[,-c(1:4)] %>% colSums() %>% t() %>% as.data.frame()
  tarline <- data.frame(class = classes[[i]], group = targroup, time = tartime, average)
  
  if(i==1){
     df_log_average <- tarline
  }else{
     df_log_average <- rbind(df_log_average, tarline)
  }
  }

df_heat <- df_log_average %>%
  mutate(time = factor(time, levels = times)) %>%
  arrange(time)

hdata <- data.frame(df_heat[,-c(1:3)]) %>% scale() %>% as.data.frame()
rownames(hdata) <- df_heat$class
colnames(hdata) <- df_heat[,-c(1:3)] %>% colnames()

hdata_t<- t(hdata) %>% as.data.frame()
col_fun <- colorRamp2(c(min(hdata_t), 0, max(hdata_t)), c("navy", "white", "firebrick3"))
split_vector <- rep(1:ceiling(ncol(hdata_t) / 4), each = 4, length.out = ncol(hdata_t))


# Figure 4a
heatmap <- Heatmap(hdata_t,
                   col = col_fun,
                   cluster_columns = FALSE,
                   column_split = split_vector, 
                   border_gp = gpar(col = "black", lwd = 1),
                   show_row_names = FALSE,
                   show_column_names = TRUE,
                   column_title = NULL,
                   show_row_dend = FALSE,
                   column_gap = unit(0, "mm"), 
                   column_names_gp = gpar(fontsize = 8),
                   heatmap_legend_param = list(title = "Z-score", at = c(floor(min(hdata_t)), 0, ceiling(max(hdata_t)))),
                   width = unit(4, "cm"), height = unit(6, "cm")
)

heatmap


 svg(paste0(path, "/heatmap/", "heatmap_v2.svg"), width = 4, height = 6)
 plot(heatmap)
 dev.off()
 

```


```{r Figure 1c and 4b}


# Figure 1c
col_class <- c("#424242", "#B71C1C", "#0D47A1","#00695C")

for (i in c(1:length(times))) {
  # i=1
  tartime <- times[i]
  xdata <- df_log_sig %>% filter(time== tartime)
  pca <- prcomp(xdata[,-c(1:4)], scale=TRUE)
  
  score <- xdata[,c(1:4)] %>% cbind(pca$x)
  order <- xdata %>% distinct(class, .keep_all = F) %>% .$class
  score <- transform(score, class = factor(class, levels = order))
  rotation <- pca$rotation %>% as.data.frame()

  p <- ggplot(score, aes(x=PC1, y=PC2, fill = class))+
  geom_point(color = "black",size = 5, shape = 21)+
  labs(x = paste("PC1:",round(pca$sdev[1], digits=2),"%"),
       y = paste("PC2:",round(pca$sdev[2], digits=2),"%"))+
        scale_fill_manual(values = c(col_class))+
  theme(panel.background = element_rect(fill = "transparent", colour = "black", size = 1),
        panel.grid = element_blank(),
        axis.text = element_text(size = 15, color = "black"),
        axis.title = element_text(size = 18, colour = "black"),
        axis.line = element_line(colour="transparent"),
        # legend.position = "bottom",
        legend.position = "none",
        legend.key.size = unit(0.1, "cm"),
        legend.key = element_rect(fill = NA, color = NA),
        legend.title = element_blank(),
        legend.text = element_text(size = 12),
        aspect.ratio = 1)


 p

 filename <- paste0("D:/2024/01_kanzo_multiomics/03_Phosphoproteomics/PCA/pca_pc3-4_", tartime, ".svg")
 svg(filename, width = 3, height = 3)
 print(p)
 dev.off()
 
  }


# Figure 4b
control <- colorRampPalette(c("#F5F5F5", "#424242"))
LPS <- colorRampPalette(c("#F5F5F5", "#B71C1C"))
ILG <- colorRampPalette(c("#F5F5F5", "#0D47A1"))
GU <- colorRampPalette(c("#F5F5F5", "#00695C"))

num <- 4
col_con <- control(num)
col_LPS <- LPS(num)
col_ILG <- ILG(num)
col_GU <- GU(num)
pal <- list(col_con, col_LPS, col_ILG, col_GU)
 
for (i in c(1:length(groups))) {
  # i=1
  xdata <- df_log_sig %>% filter(group== groups[i])
  pca <- prcomp(xdata[,-c(1:4)], scale=TRUE)
  
  score <- xdata[,c(1:4)] %>% cbind(pca$x)
  order <- xdata %>% distinct(class, .keep_all = F) %>% .$class
  score <- transform(score, class = factor(class, levels = order))
  rotation <- data.frame(pca$rotation)
  
  filename <- paste0("D:/2024/01_kanzo_multiomics/05_paper_work/Source data/pca/score_", groups[i], ".xlsx")
  write.xlsx(score,filename ,rowNames = TRUE)

 p <- ggplot(score, aes(x=PC1, y=PC2, fill = class))+
 geom_point(color = "black",size = 5, shape = 21)+
 labs(x = paste("PC1:",round(pca$sdev[1], digits=2),"%"),
      y = paste("PC2:",round(pca$sdev[2], digits=2),"%"))+
 scale_fill_manual(values = pal[[i]])+
 theme(panel.background = element_rect(fill = "transparent", colour = "transparent", size = 1),
       panel.grid = element_blank(),
       axis.text = element_text(size = 15, color = "black"),
       axis.title = element_text(size = 18, colour = "black"),
       axis.line = element_line(colour="black"),
       legend.position = "none",
       legend.key.size = unit(0.1, "cm"),
       legend.key = element_rect(fill = NA, color = NA),
       legend.title = element_blank(),
       legend.text = element_text(size = 12),
       aspect.ratio = 1)

p

filename <- paste0(path, "PCA/pca_", groups[i], ".svg")
svg(filename, width = 3, height = 3) #3.5
print(p)
dev.off()
 
  }

```


```{r Figure 4c}

#Tukey's HSD (vs. Control)
times <- df_log_sig %>% distinct(time) %>% .$time

for (i in c(1:length(times))) {
  # i=1
  tardf <- df_log_sig %>% filter(time == times[i])
  
  for (j in c(5:ncol(tardf))) {
    # j=5
    testdf <- tardf[,c(1:4)] %>% mutate("Log2Height" = tardf[,j])
    tukeydf <- TukeyHSD(aov(testdf$Log2Height ~ testdf$group))
    result <- data.frame(pair = rownames(tukeydf$`testdf$group`), tukeydf$`testdf$group`)
    
    p_LPS <- filter(result, pair == "LPS-Control") %>% .$p.adj
    p_ILG <- filter(result, pair == "ILG-Control") %>% .$p.adj
    p_GU <- filter(result, pair == "GU-Control") %>% .$p.adj
    
    p <- rbind(p_LPS,p_ILG,p_GU) %>% as.data.frame()
    colnames(p) <- colnames(tardf)[j]
    

     if(j == 5){
      p_ont <- data.frame("Name"= paste0(c("LPS", "ILG", "GU"), "_", times[i]), "time" = times[i]) %>% cbind(p)
      } else{
      p_ont <- cbind(p_ont, p)
      }
  }
  if(i == 1){
    df_p_ont <- p_ont
  }else{
    df_p_ont <- rbind(df_p_ont, p_ont)
  }
}

## Fold-change (vs. Control)
for (i in c(1:length(times))) {
  # i=3
  tardf <- df_log_average %>% filter(time == times[i])
  
  rowinfo <- data.frame(class =  tardf[-1,1], group= tardf[-1,2], time = tardf[-1,3])
  fcdf <- rbind((tardf[2,-c(1:3)]-tardf[1,-c(1:3)]), (tardf[3,-c(1:3)]-tardf[1,-c(1:3)]), (tardf[4,-c(1:3)]-tardf[1,-c(1:3)]))
  result <- cbind(rowinfo,fcdf) 

  if(i == 1){
    df_result <- result
  }else{
    df_result <- rbind(df_result, result)
  }
}


path <- "D:/2024/01_kanzo_multiomics/03_Phosphoproteomics/volcano-plot/"
df <- data.frame(t(df_p_ont[,-c(1,2)]), t(df_result[,-c(1:3)]))
colnames(df) <- c(paste0("p_", df_p_ont$Name), paste0("fc_", df_p_ont$Name))


for (i in c(1:12)) {
  # i=1
  tardf <- df[,c(i, (i+12))]
  colnames(tardf) <- c("p", "fc") 
  tardf <- tardf %>% mutate(sig = ifelse(p < 0.05, ifelse(fc > 1, "up", ifelse(fc < -1, "down", "x")), "xx"))
  if(i==1){
    sigdf <- data.frame(Name = rownames(df), tardf[,3])
  }else{
    sigdf <- sigdf %>% cbind(tardf[,3])
  }
}
colnames(sigdf) <- c("Name", df_p_ont$Name)


# Count the number of significantly changed phosphopeptides 
library(stringr)
df_count <- data.frame("Name"= df_p_ont$Name, "class" = str_split_fixed(df_p_ont$Name, "_", 2)[,1],
                       "time"= str_split_fixed(df_p_ont$Name, "_", 2)[,2],
                       "up" = colSums(sigdf == "up")[-1], "down"=colSums(sigdf == "down")[-1])

df_count_up <- data.frame(df_count[,c(1:3)], count = df_count[,4],type = "up") %>%
  transform(time = factor(time, levels = times), class = factor(class, levels = groups))
df_count_down <- data.frame(df_count[,c(1:3)], count = df_count[,5],type = "down") %>%
  transform(time = factor(time, levels = times), class = factor(class, levels = groups))

# Figure 4c 
  p <- ggplot(df_count_down, aes(x=time, y=count, fill = class))+ # Increased: df_count_up, Decreased: df_count_down
    geom_line(aes(group = class, col = class),size = 1)+
    geom_point(color = "black",size = 3, shape = 21)+
    scale_fill_manual(values= c("#B71C1C", "#0D47A1","#00695C"))+
    scale_color_manual(values= c("#B71C1C", "#0D47A1","#00695C"))+
    labs(y = "Significantly changed peptides")+
    theme(panel.background = element_rect(fill = "transparent", colour = "transparent", size = 1),
        panel.grid = element_blank(),
        axis.text = element_text(size = 12, color = "black"), 
        axis.title = element_text(size = 12, colour = "black"),
        axis.line = element_line(colour="black"),
        legend.position = "right",
        axis.title.x = element_blank(),
        legend.key.size = unit(0.1, "cm"),
        legend.key = element_rect(fill = NA, color = NA), 
        legend.title = element_blank(),
        legend.text = element_text(size = 12),
        aspect.ratio = 0.8)

 p
 
 filename <- paste0(path, "lineplot_down",".svg")
 svg(filename, width = 3.5, height = 3)
 print(p)
 dev.off()

```


```{r Figure 4d}

library(clusterProfiler)
library(org.Mm.eg.db)
library(ReactomePA)
library(pathview)
library(enrichplot)
library(ggplot2)
library(dplyr)
library(openxlsx)
library(tidyr)

raw_info <- read.xlsx("D:/2024/01_kanzo_multiomics/05_paper_work/Source data/20220825_TsugawaLab_STY_79.9663_FB_20230124.xlsx",) %>% .[,c(1:8)]
colnames(raw_info) <- c("Name", colnames(raw_info)[-1])
test_result <-  data.frame("Name" = rownames(df), sigdf) %>% left_join(raw_info[,c(1,6,7)], by = "Name")

for (i in c(2:length(groups))) {
  # i=3
  test_result_i <- test_result %>% dplyr::select(1, 14,15, all_of(paste0(groups[i], "_",times)))
  colnames(test_result_i) <- c(colnames(test_result)[c(1,14,15)], paste0("siglab_",times))
  up_0h <- test_result_i %>% filter(siglab_0h == "up") %>% distinct(Gene) %>% unlist()
  up_15m <- test_result_i %>% filter(siglab_15m == "up") %>% distinct(Gene) %>% unlist()
  up_2h <- test_result_i %>% filter(siglab_2h == "up") %>% distinct(Gene) %>% unlist()
  up_24h <- test_result_i %>% filter(siglab_24h == "up") %>% distinct(Gene) %>% unlist()
  test <- list(up_0h = up_0h, up_15m = up_15m, up_2h = up_2h, up_24h = up_24h)
  
  sigcount <- data.frame("count" = c(length(up_0h),length(up_15m),length(up_2h),length(up_24h)), class = paste0(groups[i], "_", times))
  
  for (j in c(1:length(times))) {
    # j=2
    gene_entrez <- bitr(test[[j]], fromType = "SYMBOL", toType = "ENTREZID", OrgDb = org.Mm.eg.db)
    
    #KEGG
    kegg_enrich <- enrichKEGG(gene = gene_entrez$ENTREZID, organism = 'mmu')
    kegg_table <- as.data.frame(kegg_enrich) %>% mutate("group" = groups[i], "time" = times[j], "class" = paste0(groups[i], "_", times[j]))
    if (nrow(kegg_table)>0) {
      kegg_table <- kegg_table %>% mutate(order = c(1:nrow(kegg_table)))
    } else {
      message("kegg_table does not exist. Skip the process.")
    }
    
    
    if(j==1){
      GO_KEGG <- kegg_table
    }else{
      GO_KEGG <- rbind(GO_KEGG, kegg_table)
    }
  }
  
  if(i==2){
    GO_KEGG_all <- GO_KEGG
    sigcount_all <- sigcount
  }else{
    GO_KEGG_all <- rbind(GO_KEGG_all, GO_KEGG)
    sigcount_all <- rbind(sigcount_all, sigcount)
  }
}

write.xlsx(GO_KEGG_all, paste0(path, "up_GO_KEGG_all.xlsx"), rowNams = F)

#Down
for (i in c(2:length(groups))) {
  # i=2
  test_result_i <- test_result %>% dplyr::select(1, 14,15, all_of(paste0(groups[i], "_",times)))
  colnames(test_result_i) <- c(colnames(test_result)[c(1,14,15)], paste0("siglab_",times))
  down_0h <- test_result_i %>% filter(siglab_0h == "down") %>% distinct(Gene) %>% unlist()
  down_15m <- test_result_i %>% filter(siglab_15m == "down") %>% distinct(Gene) %>% unlist()
  down_2h <- test_result_i %>% filter(siglab_2h == "down") %>% distinct(Gene) %>% unlist()
  down_24h <- test_result_i %>% filter(siglab_24h == "down") %>% distinct(Gene) %>% unlist()
  test <- list(down_0h = down_0h, down_15m = down_15m, down_2h = down_2h, down_24h = down_24h)

  sigcount <- data.frame("count" = c(length(down_0h),length(down_15m),length(down_2h),length(down_24h)), class = paste0(groups[i], "_", times))

  for (j in c(1:length(times))) {
    # j=2
    gene_entrez <- bitr(test[[j]], fromType = "SYMBOL", toType = "ENTREZID", OrgDb = org.Mm.eg.db)

    #KEGG
    kegg_enrich <- enrichKEGG(gene = gene_entrez$ENTREZID, organism = 'mmu')#pairwise_termsim()
    kegg_table <- as.data.frame(kegg_enrich) %>% mutate("group" = groups[i], "time" = times[j], "class" = paste0(groups[i], "_", times[j]))
    if (nrow(kegg_table)>0) {
      kegg_table <- kegg_table %>% mutate(order = c(1:nrow(kegg_table)))
    } else {
      message("kegg_table does not exist. Skip the process.。")
    }


    if(j==1){
      GO_KEGG <- kegg_table
    }else{
      GO_KEGG <- rbind(GO_KEGG, kegg_table)
    }
  }

  if(i==2){
    down_GO_KEGG_all <- GO_KEGG
    down_sigcount_all <- sigcount
  }else{
    down_GO_KEGG_all <- rbind(down_GO_KEGG_all, GO_KEGG)
    down_sigcount_all <- rbind(down_sigcount_all, sigcount)
  }
}


rank <-6
types <- "up" # Increased:up, Decreased: down

sigcount_df <- rbind(sigcount_all %>% mutate("type" ="up"),
                     down_sigcount_all %>% mutate("type" ="down"))
upterms <- GO_KEGG_all %>% filter(order < rank) %>% distinct(Description) %>% .$Description
downterms <- down_GO_KEGG_all %>% filter(order < rank) %>% distinct(Description) %>% .$Description

df_KEGG <- rbind(GO_BP_all %>% filter(Description %in% upterms) %>% mutate("type" ="up"),
               down_GO_KEGG_all %>% filter(Description %in% downterms) %>% mutate("type" ="down")) %>% mutate(Description = str_remove(Description, " - Mus musculus \\(house mouse\\)"))


df_KEGG <- df_KEGG %>% mutate("-log10P" = -log10(df_KEGG$p.adjust))


plot_df_KEGG <- df_KEGG %>% filter(type == types) # up down
plot_sigcount <- sigcount_df %>% filter(type == types)  # up down

plot_df_KEGG <- transform(plot_df_KEGG, class = factor(class, levels = unique(plot_df_KEGG$class)))

unique_data <- plot_df_KEGG %>%
  distinct(Description, class, .keep_all = TRUE)

heatmap_data <- unique_data %>%
  dplyr::select(Description, class, `-log10P`) %>%  
  pivot_wider(names_from = class, values_from = `-log10P`) %>% as.data.frame()

rownames(heatmap_data) <- heatmap_data$Description
heatmap_data_2 <- heatmap_data[,-1]

missing_cols <- setdiff(classes[-c(1:4)], colnames(heatmap_data_2))
for (col in missing_cols) {
  heatmap_data_2[[col]] <- NA  
}

heatmap_data_2 <- heatmap_data_2 %>% 
  dplyr::select(all_of(classes[-c(1:4)]))

annot.col <- data.frame("sig_count" = plot_sigcount$count, "class" = rep(groups[-1], each = 4))
rownames(annot.col) <-classes[-c(1:4)]
annotation_colors <- list("sig_count" = colorRampPalette(c("#fef6e7","#F2A71B"))(100),
                          "class" = c("LPS" = "#B71C1C", "ILG" = "#0D47A1", "GU" = "#00695C"))

heatmap_order <- unique_data %>%
  dplyr::select(Description, class, order) %>%  
  pivot_wider(names_from = class, values_from = order) %>% as.data.frame()

missing_cols <- setdiff(classes[-c(1:4)], colnames(heatmap_data_2))
for (col in missing_cols) {
  heatmap_data_2[[col]] <- NA 
}

heatmap_data_2 <- heatmap_data_2 %>% 
  dplyr::select(all_of(classes[-c(1:4)]))

heatmap_order_lab <- heatmap_order[,-1] %>% mutate_all(~ ifelse(. < rank, "+", ""))
missing_cols <- setdiff(classes[-c(1:4)], colnames(heatmap_order_lab))
for (col in missing_cols) {
  heatmap_order_lab[[col]] <- NA
}

heatmap_order_lab <- heatmap_order_lab %>% 
  dplyr::select(all_of(classes[-c(1:4)])) %>% mutate_all(~ ifelse(is.na(.), "", .))

rownames(heatmap_order_lab) <- heatmap_order$Description


library(pheatmap)
heatmap <- pheatmap(heatmap_data_2,
                    color = colorRampPalette(c("#FAEFEF", "#BB0000"))(500),
                    annotation_col = annot.col,
                    annotation_colors = annotation_colors,
                    cellwidth = 10, cellheight = 10,
                    col_split = 4,
                    cluster_rows = F,cluster_cols = F,
                    show_colnames =T,show_rownames = T,
                    treeheight_row = F,treeheight_col = F,
                    border_color='gray40',
                    border_width = 0.1,
                    na_col = "transparent",
                    annotation_names_col = FALSE,
                    display_numbers = heatmap_order_lab,
                     number_color = "black",
                    fontsize_row = 10, fontsize_col = 10)

heatmap

svg(paste0(path, "BP_",types,"-top",(rank-1),"_v1.svg"), width = 12, height = 6)
print(heatmap)
dev.off()
 

```


```{r Figure 4e}

# Heatmap(square)
Index <- colnames(df_log_sig)[-c(1:4)]
split_vec <- strsplit(Index, "_")

df_Index <- data.frame(
  Name = Index,
  ID = sapply(split_vec, `[`, 1), 
  site = sapply(split_vec, `[`, 2)
)

Index_unique <- df_Index %>% distinct(ID) %>% .$ID
asterisks <- cbind(df_p_ont[,c(1,2)], ifelse(df_p_ont[,-c(1,2)] < 0.05, "*", ""))


test <- data.frame(matrix("", nrow = 4, ncol = ncol(df_p_ont)-2)) %>% lapply(as.character) %>% as.data.frame()

con2asterisks <-data.frame("Name"= paste0("Control_", times), "time" = times, test)
colnames(con2asterisks) <- colnames(asterisks)
asterisks_2 <- rbind(con2asterisks, asterisks)


col_annot <- data.frame(group = factor(groups)) #column annotation
rownames(col_annot) <- groups
annot_colors <- list(group = setNames(groups, c("#424242", "#B71C1C", "#0D47A1","#00695C")))
annot_colors <- list(group = c("Control" = "#424242", "LPS" = "#B71C1C", "ILG" = "#0D47A1", "GU" = "#00695C"))

path <- "D:/2024/01_kanzo_multiomics/03_Phosphoproteomics/heatmap/ForPathway/v2"
colnames(df_log_average) <- colnames(df_log_sig)[-1]

for (i in c(1:length(times))) {
  # i=1
  hdf <- df_log_average %>% filter(time == times[i])
  adf <- asterisks_2 %>% filter(time == times[i])
  
  for (j in c(1:length(Index_unique))) {
    # j=1
    tarname <- df_Index %>% filter(ID == Index_unique[j]) %>% .$Name
    hdf_ID <- select(hdf, all_of(tarname)) %>% scale() %>% t() %>% data.frame()
    adf_ID <- select(adf, all_of(tarname)) %>% t() %>% data.frame()
    site <- df_Index %>% filter(ID == Index_unique[j]) %>% .$site
    rownames(hdf_ID) <- site
    colnames(hdf_ID) <- groups
    rownames(adf_ID) <- site
    colnames(adf_ID) <- groups
    
    cluster_rows_condition <- ifelse(nrow(hdf_ID) > 1, TRUE, FALSE)
    
     heatmap <- pheatmap::pheatmap(hdf_ID,
                              col = colorRampPalette(c("navy","white", "firebrick3"))(500),
                              breaks = c(seq(min(hdf_ID), 0, length.out=250), seq(0.01, max(hdf_ID), length.out=251)),
                              annotation_col = col_annot,           
                              annotation_colors = annot_colors,     
                              display_numbers = adf_ID,          
                               celltext_color = "black",
                              fontsize_number = 12,
                              border_color='gray20',
                              fontsize_row = 6,
                              treeheight_row = 0,
                              cellheight = 10,  cellwidth = 10,
                              show_colnames = F,
                              show_rownames = T,
                              annotation_names_col = F,
                              legend = F,
                              annotation_legend = F,
                              # legend_labels = c("", "0", ""),
                              cluster_rows = cluster_rows_condition,
                              cluster_cols = F)
 
 
 svg(paste0(path,"/",times[i],"/", Index_unique[j], ".svg"), width = 2, height = 2) #3.5
 print(heatmap)
 dev.off()
    
  }
  
}


# Heatmap (pie chart)
FC_vsCon <- data.frame(df_log_average[c(5:8), c(1:3)], df_log_average[c(5:8), -c(1:3)]-df_log_average[c(1:4), -c(1:3)]) %>%
  rbind(data.frame(df_log_average[c(9:12), c(1:3)], df_log_average[c(9:12), -c(1:3)]-df_log_average[c(1:4), -c(1:3)]))%>%
  rbind(data.frame(df_log_average[c(13:16), c(1:3)], df_log_average[c(13:16), -c(1:3)]-df_log_average[c(1:4), -c(1:3)]))

colnames <- gsub("\\.", "-", colnames(FC_vsCon))
colnames(FC_vsCon) <- colnames

for (i in c(1:length(times))) {
  # i=1
  df_i <- FC_vsCon %>% filter(time == times[i])
   for (j in c(4:ncol(df_i))) {
     # j=4
     tardf <- df_i[,j] %>% t() %>% as.data.frame()
     colnames(tardf) <- c(groups[-1])
     tardf$x <- 1
     tardf$y <- 1
     tardf <- pivot_longer(tardf,cols= groups[-1],names_to = "colorby") %>%
       mutate("ratio" = rep(1/3, 3), "radius" = rep(0.75, 3)) %>%
       mutate("start" = c(0, 2*pi/3, 4*pi/3),
              "end" = c(2*pi/3, 4*pi/3, 2*pi))
     
     g <- ggplot(tardf)+
       geom_arc_bar(
         aes(x0 = x, y0 = y, r0 = 0, r = 1,
             start = start, end = end, fill = value),
         color = "black", size = 0.5) +
       coord_equal()+ 
       scale_fill_gradient2(low = "#0D47A1", mid = "white", high = "#B71C1C", midpoint = 0,
                            limits = c(-10, 10), oob = scales::squish)+
       theme(panel.background = element_rect(fill = "transparent", colour = "transparent", size = 1),
             plot.background = element_rect(fill = "transparent", color = NA),
             panel.grid = element_blank(),
             axis.text = element_blank(),
             axis.title =element_blank(),
             axis.line = element_line(colour="transparent"),
             legend.position = "bottom",
             plot.margin = margin(1, 0, 0, 0, "cm"),
             legend.key = element_rect(fill = NA, color = NA), 
             legend.title = element_blank(),
             legend.text = element_text(size = 12),
             aspect.ratio = 1)
     g
     
     svg(paste0("D:/2024/01_kanzo_multiomics/03_Phosphoproteomics/kinase_pred/",
                times[i], "/",colnames(df_i)[j], ".svg"),  width = 2, height = 2)
     print(g)
     dev.off()
     
     
     filaname <- paste0("file:/D:/2024/01_kanzo_multiomics/03_Phosphoproteomics/kinase_pred/",
                        times[i], "/",colnames(df_i)[j], ".svg")
     path_row <- data.frame("path" = filaname, "name" = colnames(df_i)[j] )
     if(j==4){
       pathdf <- path_row
     }else{
       pathdf <- rbind(pathdf, path_row)
     }
   }
 
  write.csv(pathdf,paste0("D:/2024/01_kanzo_multiomics/03_Phosphoproteomics/kinase_pred/",
                          times[i], "/filepath_list.csv"), col.names = F)
}

```
````
