## Supplementary material for "Unraveling anti-inflammatory metabolic signatures of *Glycyrrhiza uralensis* and isoliquiritigenin through multiomics": Source data: Source code for Figure 5 (multiset-plsrog)_vs2.Rmd

````
---
title: "R Notebook"
output: html_notebook
---

This is an [R Markdown](http://rmarkdown.rstudio.com) Notebook. When you execute code within the notebook, the results appear beneath the code. 

Try executing this chunk by clicking the *Run* button within the chunk or by placing your cursor inside it and pressing *Ctrl+Shift+Enter*. 

```{Figure 5a and 5b}
library(ggplot2)
library(loadings)
library(openxlsx)
library(dplyr)
library(exactRankTests)
library(ggplot2)
library(dplyr)
library(openxlsx)
library(ggsignif)
library(plotrix)
library(loadings)
library(tidyverse)
library(pheatmap)
library(plotrix)

setwd("D:/2024/01_kanzo_multiomics/05_paper_work/Source data/5_integrated omics analysis")


metabolome <-  read.xlsx("Source-data_integrated-analyisis.xlsx",1,, check.names = FALSE)
proteome <-  read.xlsx("Source-data_integrated-analyisis.xlsx",2, check.names = FALSE) 
lipidome <-  read.xlsx("Source-data_integrated-analyisis.xlsx",3, check.names = FALSE) 
 
row.names(metabolome) <- metabolome[,1]
row.names(proteome) <- proteome[,1]
row.names(lipidome) <- lipidome[,1]


plsrog_met <- metabolome %>% filter(time == "24h") %>% .[-c(4,8,12,16),-c(1:4)]
plsrog_pro <- proteome %>% filter(time == "24h") %>% .[,-c(1:4)]
plsrog_lip <- lipidome %>% filter(time == "24h") %>% .[,-c(1:4)]

plsrog_met_2 <- plsrog_met[, colSums(plsrog_met != 0) > 0]
plsrog_pro_2 <- plsrog_pro[, colSums(plsrog_pro != 0) > 0]
plsrog_lip_2 <- plsrog_lip[, colSums(plsrog_lip != 0) > 0]

# Multiset PLS-ROG
X <- list("metabolome" = plsrog_met_2, "proteome" = plsrog_pro_2, "lipidome" = plsrog_lip_2)
Y <- read.xlsx("Source-data_integrated-analyisis.xlsx",4)
tau <- read.xlsx("Source-data_integrated-analyisis.xlsx",6) 
D <- read.xlsx("Source-data_integrated-analyisis.xlsx",5)

multiplsrog <- multipls_rog(X,Y,tau,D) 
multiplsrog <- multipls_loading(multiplsrog)


#score plot
scoredf <- data.frame(multiplsrog$U) %>% mutate(group = rep(c("Control", "LPS", "ILG", "GU"), each = 3)) %>% mutate(col = c(rep("black",3), rep("red4",3), rep("darkblue",3),rep("darkgreen",3))) %>%
  transform(group = factor(group, levels = c("Control", "LPS", "ILG", "GU")))

ScorePlot <- ggplot(scoredf, aes(x=X1, y=X2))+ 
  geom_point(shape = 21, fill = plotdata$col, color = "black", size = 5)+
  theme_classic() +   labs(x = "PLS1", y="PLS2") + theme(axis.title = element_text(size = 20), axis.text = element_text(size = 20))+
  theme(aspect.ratio = 1)

ScorePlot

svg("D:/2024/01_kanzo_multiomics/05_paper_work/Source data/5_integrated omics analysis/multi-PLSROG_24h.svg",width = 4, height = 4)
 print(ScorePlot)
 dev.off()

#A matrix with PLS-ROG score for response variable in each column
write.csv(data.frame(group = c(rep("control",3), rep("LPS",3), rep("ILG",3),rep("GU",3)),multiplsrog$U),"D:/2024/01_kanzo_multiomics/05_paper_work/Source data/5_integrated omics analysis/Score_multi-PLSROG_24h.csv" )

#A matrix with p-values
write.csv(data.frame(multiplsrog$loading$p.value[[3]]),"D:/2024/01_kanzo_multiomics/05_paper_work/Source data/5_integrated omics analysis/P_lip_24h.csv" )

#A matrix with loadongs
write.csv(data.frame(multiplsrog$loading$R[[1]]),"D:/2024/01_kanzo_multiomics/05_paper_work/Source data/5_integrated omics analysis/Liading_met_24h.csv" )
```

```{r}
library(ggplot2)
library(dplyr)
library(openxlsx)
library(tidyverse)

df <- read.xlsx("C:/Users/saki kiuchi/Downloads/Source data for Figure 5 (multiset-plsrog)_vs2.xlsx", 8)


# Figure 5b
df2 <- df %>% filter(Raw.p < 0.01 & Axis == "PLS1_POS") %>% rbind(df %>% filter(Axis == "PLS1_NEG") %>% .[9,])
bardf <- df2[order(df2$`"-LOG10(p)"`,decreasing = F),] %>% mutate("Hits ratio" = Hits/Total)

order <- bardf$Pathway %>% unlist()
bardf <- transform(bardf, Pathway = factor(Pathway, levels = order))


bardf$logP <- -log10(bardf$Raw.p)

min_p <- min(bardf$logP)
max_p <- max(bardf$logP)
min_hits <- min(bardf$Hits.ratio)
max_hits <- max(bardf$Hits.ratio)

scale_factor <- (max_p - min_p) / (max_hits - min_hits)

bar <- ggplot(bardf, aes(x = logP, y = reorder(Pathway, logP))) +
  geom_col(aes(fill = Axis), col= "black") +
  geom_point(aes(x = (Hits.ratio - min_hits) * scale_factor + min_p, color = Impact), size = 3) +
  scale_color_gradient(low = "white", high = "yellow") +
  scale_fill_manual(values = c("PLS1_POS" = "#0D47A1", "PLS1_NEG" = "#00695C")) +
  labs(x = "-Log10(P-value)") +
  theme(axis.text.y = element_text(size = 10, colour="black"),
        axis.text.x = element_text(size = 10, colour="black"),
        axis.title.y = element_blank(),
        panel.grid.major.y = element_blank(),
        panel.background = element_rect(fill = "transparent", colour = "transparent"),
        axis.line = element_line(colour="black"),
        panel.grid = element_blank(),
        aspect.ratio = 1.5) +
  scale_x_continuous(
    sec.axis = sec_axis(~ (. - min_p) / scale_factor + min_hits, name = "Hits ratio")
  )

bar

# svg("C:/Users/saki kiuchi/Downloads/PLSROG_PLS1=24h.svg",width = 6, height = 2.5)
#  print(bar)
#  dev.off()

 
# Figure S4c
df3 <- df %>% filter(Raw.p < 0.01 & Axis == "PLS2_POS") 
bardf <- df3[order(df2$`"-LOG10(p)"`,decreasing = F),] %>% mutate("Hits ratio" = Hits/Total)

order <- bardf$Pathway %>% unlist()
bardf <- transform(bardf, Pathway = factor(Pathway, levels = order))


bardf$logP <- -log10(bardf$Raw.p)

min_p <- min(bardf$logP)
max_p <- max(bardf$logP)
min_hits <- min(bardf$Hits.ratio)
max_hits <- max(bardf$Hits.ratio)

scale_factor <- (max_p - min_p) / (max_hits - min_hits)

bar <- ggplot(bardf, aes(x = logP, y = reorder(Pathway, logP))) +
  geom_col(fill = "#C62828", col= "black") +
  geom_point(aes(x = (Hits.ratio - min_hits) * scale_factor + min_p, color = Impact), size = 3) +
  scale_color_gradient(low = "white", high = "yellow") +
  labs(x = "-Log10(P-value)") +
  theme(axis.text.y = element_text(size = 10, colour="black"),
        axis.text.x = element_text(size = 10, colour="black"),
        axis.title.y = element_blank(),
        panel.grid.major.y = element_blank(),
        panel.background = element_rect(fill = "transparent", colour = "transparent"),
        axis.line = element_line(colour="black"),
        panel.grid = element_blank(),
        aspect.ratio = 1.5) +
  scale_x_continuous(
    sec.axis = sec_axis(~ (. - min_p) / scale_factor + min_hits, name = "Hits ratio")
  )

bar
# svg("C:/Users/saki kiuchi/Downloads/PLSROG_PLS2_24h.svg",width = 6, height = 2.5)
#  print(bar)
#  dev.off()

```
````
