## Supplementary material for "Unraveling anti-inflammatory metabolic signatures of *Glycyrrhiza uralensis* and isoliquiritigenin through multiomics": Source data: Source code for Figure 6.Rmd

````
---
title: "R Notebook"
output: html_notebook
---

This is an [R Markdown](http://rmarkdown.rstudio.com) Notebook. When you execute code within the notebook, the results appear beneath the code. 

Try executing this chunk by clicking the *Run* button within the chunk or by placing your cursor inside it and pressing *Ctrl+Shift+Enter*. 

```{r Figure 6a}
setwd("D:\\2023\\Others\\sakai")
library(ggplot2)
library(openxlsx)
library(dplyr)
library(plotrix)

scientific_notation <- function(x) {
  x <- format(x, scientific = TRUE)
  x <- gsub("^(.*)e", "'\\1'e", x)
  x <- gsub("e", "%*%10^", x)
  x <- gsub('\\+', '', x)
  parse(text = x)
}


setwd("D:/2024/01_kanzo_multiomics/05_paper_work/Source data/6_GABA")
data <- read.xlsx("Source code for GABA.xlsx", 1)
order <- c("control", "LPS","GABA/GU/LPS", "GABA/LPS", "GABA/GU", "GABA")
score <- transform(data, class = factor(class, levels = order))


data <- transform(data, class= factor(class, levels = order)) #見た目はdfと同じ
time <- data %>% distinct(Time) %>% .$Time

for (i in c(1:2)) {
  # i=1
  plotdf <- data %>% filter(Time == time[i])
  
  for (j in c(4:7)) {
    # j=4
    plotdf_j <- data.frame(plotdf[,c(1:3)], height = plotdf[,j])
    Name <- colnames(plotdf) %>% .[j]
    
    g <- ggplot(plotdf_j, aes(x = class, y = height, fill = class))+
      geom_boxplot(alpha=0.5)+
      scale_fill_manual(values = c("grey40", "#B20000","#4E58FC", "#000094","#5FA86F", "#006100"))+
      geom_jitter(shape = 21, width = .1, size = 2)+
      
      labs(y="Peak height")+
      scale_y_continuous(expand = c(0, 0), limits = c(0, max(plotdf_j$height)*1.2), labels = scientific_notation)+
      theme(
        #axis.text.x = element_blank(),
        panel.background = element_rect(fill = "transparent", colour = "transparent"),
        panel.grid = element_blank(),
        axis.line = element_line(colour="black"),
        axis.text.x = element_blank(),
        axis.text.y = element_text(size = 12, color= "grey10"),
        axis.title.y = element_text(size = 14, color= "grey10" ),
        axis.title.x = element_blank(),
        legend.background = element_blank(),
        legend.text = element_text(size = 12),
        plot.title = element_text(hjust=0.5, size = 18, color= "grey10"),
        legend.title = element_blank(),
        aspect.ratio = 0.75)
    
    
    g
    
    filename <- paste0(Name,"_", time[i],".svg")
    svg(filename, width = 6, height = 4)
    print(g)
    dev.off()
    
    tukeydf <- TukeyHSD(aov(plotdf_j$height ~ plotdf_j$class))
    result <- data.frame(Time = time[i], metabolite = Name, pair = rownames(tukeydf$`plotdf_j$class`), tukeydf$`plotdf_j$class`)
    
    if(i ==1 & j ==4){
      tukey_result <- result
    }else{
      tukey_result <- rbind(tukey_result, result)
    }
    
    
    
  }
}

write.xlsx(tukey_result, "tukey_result.xlsx")


```


```{r Figure 6b and S5b}
library(pheatmap)
library(loadings)
library(tidyverse)
library(ggplot2)
library(caret)
library(tidyverse)
library(openxlsx)
library(dplyr)
library(plotrix)

scientific_notation <- function(x) {
  x <- format(x, scientific = TRUE)
  x <- gsub("^(.*)e", "'\\1'e", x)
  x <- gsub("e", "%*%10^", x)
  x <- gsub('\\+', '', x)
  parse(text = x)
}


setwd("D:/2024/01_kanzo_multiomics/LCMS/20250306_allylglycine_vs2")
df <- read.xlsx("Height_1_2025_03_18_14_59_16.xlsx", 1)
df_2 <- df %>% .[-c(1:4),c(4,19,36:55)]  %>% mutate(across(3:22, as.numeric))
colnames(df_2) <- df[4,c(4,19,36:55)]

class <- colnames(df) %>% .[c(36:55)]
sampletype <- df[1,c(36:55)] %>% unlist()

df_target <- df_2 %>% filter(grepl("Confirmed", Comment))
df_target_t <- data.frame(colnames(df_target[,-c(1,2)]), class, sampletype,t(df_target[,-c(1,2)]))
colnames(df_target_t) <- c("Name", "Class", "SampleType", "GABA",df_target$`Metabolite name`[-1])

df_target_t <- df_target_t %>% mutate("corrected-GABA" = GABA/`GABA-d6`)
plotdf <- df_target_t %>% filter(SampleType == "Sample")
group_order <- plotdf %>% distinct(Class) %>% .$Class
plotdf_2 <- transform(plotdf, Class = factor(Class, levels = group_order))

plotdf_3 <- plotdf_2 %>% filter(Class != "ControlALG") 
pal <- colorRampPalette(c("#F5F5F5", "#424242"))
plot_pal <- pal(3)

# Figure 6b
tukeydf <- TukeyHSD(aov(plotdf_3$`corrected-GABA` ~ plotdf_3$Class))
result <- data.frame(pair = rownames(tukeydf$`plotdf_3$Class`), tukeydf$`plotdf_3$Class`)

p <- ggplot(plotdf_3, aes(x=Class, y=`corrected-GABA`, fill = Class))+
  stat_summary(fun = "mean", geom = "bar", width = .7, alpha = 0.9, col= "black") +
  stat_summary(fun.max = function(x) mean(x) + std.error(x), 
               fun.min = function(x) mean(x) - std.error(x), 
               geom = 'errorbar', width = .1)+
  geom_jitter(width = .1, size = 2, shape = 21, col = "black",fill = "gray30")+
  scale_y_continuous(expand = c(0,0), limits = c(0, max(plotdf_3$`corrected-GABA`)*1.2))+
  scale_fill_manual(values= plot_pal)+
  labs(y = "Relative abundance")+
  theme(panel.background = element_rect(fill = "transparent", colour = "transparent", size = 1),
        panel.grid = element_blank(),
        axis.text.y = element_text(size = 12, color = "black"), 
        axis.text.x = element_text(size = 12, vjust = 1, colour = "black"), 
        axis.title = element_text(size = 12, colour = "black"),
        axis.title.x = element_blank(),
        legend.position = "none",
        legend.key.size = unit(0.1, "cm"),
        legend.key = element_rect(fill = NA, color = NA), 
        legend.title = element_blank(),
        legend.text = element_text(size = 12),
        aspect.ratio = 1.25)

p

svg("GABA_corrected_vs2.svg", width = 3.5, height = 4)
print(p)
dev.off()


# Figure S5b
plotdf_4 <- plotdf_3 %>% filter(Class =="GU")
bardf_GU <- data.frame(Sample = c(plotdf_4$Name,plotdf_4$Name), Name = rep(c("GABA-d6","GABA"), each = 3), "PeakHeight" = c(plotdf_4$`GABA-d6`, plotdf_4$GABA))
bardf_GU <- transform(bardf_GU, Name = factor(Name, levels = c("GABA-d6","GABA")))

p <- ggplot(bardf_GU, aes(x=Name, y=PeakHeight, fill = Name))+
  stat_summary(fun = "mean", geom = "bar", width = .6, alpha = 1, col= "black") +
  stat_summary(fun.max = function(x) mean(x) + std.error(x), 
               fun.min = function(x) mean(x) - std.error(x), 
               geom = 'errorbar', width = .1)+
  geom_jitter(width = .1, size = 3, shape = 21, col = "black",fill = "white")+
  scale_y_continuous(expand = c(0,0), limits = c(0, max(bardf_GU$PeakHeight)*1.1), labels = scientific_notation)+
  scale_fill_manual(values= c("#b9bbca", "#0D47A1"))+
  labs(y = "Peak height")+
  theme(panel.background = element_rect(fill = "transparent", colour = "transparent", size = 1),
        panel.grid = element_blank(),
        axis.text.y = element_text(size = 12, color = "black"), 
        axis.text.x = element_text(size = 12, vjust = 1, colour = "black"), 
        axis.title = element_text(size = 12, colour = "black"),
        axis.line = element_line(colour="black"),
        legend.position = "none",
        axis.title.x = element_blank(),
        legend.key.size = unit(0.1, "cm"),
        legend.key = element_rect(fill = NA, color = NA), 
        legend.title = element_blank(),
        legend.text = element_text(size = 12),
        aspect.ratio = 1.5)

p

svg("Bar_GABAvsGABA-d6_GU_vs250318.svg", width = 3, height = 4) #3.5
print(p)
dev.off()

```


```{r Figure 6c}


library(ggplot2)
library(openxlsx)
library(dplyr)
library(ggsignif)
library(plotrix)
scientific_notation <- function(x) {
  x <- format(x, scientific = TRUE)
  x <- gsub("^(.*)e", "'\\1'e", x)
  x <- gsub("e", "%*%10^", x)
  x <- gsub('\\+', '', x)
  parse(text = x)
}


setwd("D:/2024/01_kanzo_multiomics/qPCR/20240918")

data <- read.xlsx("20240920_ILG-GABA_total -  Quantification Cq Results.xlsx", 1)
data$Cq <- lapply(data$Cq, gsub, pattern="NaN", replacement = "") %>% as.numeric()
Well <- data %>% distinct(Well) %>% .[,1]


for (i in c(1:length(Well))) {
  # i=1
  tarwell <- data %>% filter(Well == Well[[i]])
  df <- data.frame(Well = tarwell$Well[1], Target = tarwell$Target[1], class = tarwell$Biological.Set.Name[1],
                   CqMean = mean(tarwell$Cq, na.rm = T), Cq_1 = tarwell$Cq[1], Cq_2 = tarwell$Cq[2])
  
  if (i==1) {
    data_mean <- df
  }
  else{
    data_mean <- rbind(data_mean, df)
  }
}

class <- data_mean %>% distinct(class) %>% .[,1]


for (i in c(1:length(class))) {
  # i=2
  Class <- class[[i]]
  tarclass <- data_mean %>% filter(class == Class)
  Cq_ref <- tarclass %>% filter(Target == "GAPDH")
  df <- tarclass %>% mutate(dCt = CqMean-mean(Cq_ref$CqMean, na.rm = T))
  
  if (i==1) {
    data_dCq <- df
  }
  else{
    data_dCq <- rbind(data_dCq, df)
  }
  
}


target <- data_dCq %>% distinct(Target) %>% .[,1]
for (i in c(1:length(target))) {
  targene <- data_dCq %>% filter(Target == target[[i]])
  dCq_cali<- targene %>% filter(class == "control")
  df <- targene %>% mutate(ddCt = dCt-mean(dCq_cali$dCt, na.rm = T))
  
  if (i==1) {
    data_ddCq <- df
  }
  else{
    data_ddCq <- rbind(data_ddCq, df)
  }
}


order <- class
plotdf <- transform(data_ddCq, class = factor(class, levels = order))
plotdf <- plotdf %>% mutate("2^-ddCt" = 2^(-plotdf$ddCt)) %>% filter(Target != "GAPDH") %>% filter(class != "control")
plotdf <- plotdf %>% mutate(group = c(rep(rep(c("LPS","ILG", "GABA","GABA","ILG/GABA","ILG/GABA","ILG/GABA"), each= 3), 3) ))
order2 <- c("LPS","ILG", "GABA","ILG/GABA")
plotdf <- transform(plotdf, group = factor(group, levels = order2))


gene <- plotdf %>% distinct(Target)%>% .[,1]


for (i in c(1:length(gene))) {
  # i=1
  tardf <- plotdf %>% filter(Target == gene[[i]]) %>% filter(class != "ILG/GABA0.1") #%>% filter(class == "LPS" | class == "GABA10")#%>% filter(group != "GABA")
  tukeydf <- TukeyHSD(aov(tardf$`2^-ddCt` ~ tardf$class))
  result <- data.frame(Terget = gene[i], pair = rownames(tukeydf$`tardf$class`), tukeydf$`tardf$class`)
  
  p <- ggplot(tardf, aes(x = group, y = `2^-ddCt`, fill = class))+ 


    
    stat_summary(fun = "mean", geom = "bar", width = 0.8,col = "black", position=position_dodge2(preserve = "single"), alpha=0.8) +#, alpha=0.5
    stat_summary(fun.max = function(x) mean(x) + std.error(x),
                 fun.min = function(x) mean(x) - std.error(x),
                 geom = 'errorbar', width = .1,position = position_dodge(width = 0.8))+
    geom_jitter(size = 3, shape = 21,col = "black", position = position_dodge(width = 0.8))+
    labs(y = expression(2^{-Delta*Delta*Ct}))+# labs(y = expression({2^-ddCt}), expression(paste("IL-1", beta)))+
    scale_fill_manual(values = c("white", "#b9bbca","#357657", "#357657", "#0D47A1","#0D47A1"))+
    scale_y_continuous(expand = c(0, 0),limits = c(0, max(tardf$`2^-ddCt`)*1.2), labels = scientific_notation)+ #強制的に0からスタート
    theme(
      axis.text.x = element_text(size = 12, angle = 90, hjust = 1, vjust = 0.5, color= "grey10"),
      axis.text.y = element_text(size = 10, color= "grey10"),
      axis.title.y = element_text(size = 12, color= "grey10"),
      axis.title.x = element_blank(),
      legend.key.size = unit(0.5, 'cm'),
      legend.title = element_blank(),
      plot.title = element_text(hjust=0.5, size = 18, color= "grey10"),
      axis.line = element_line(colour="black"),
      panel.background = element_rect(fill = "transparent", colour = "transparent"),
      aspect.ratio = 0.8)
  
  p
  
  filepath <- paste0("D:/2024/01_kanzo_multiomics/qPCR/20240918/",gene[[i]],"_v5.svg")
  svg(filepath, width = 5, height = 3)
  print(p)
  dev.off()
  
  if(i ==1){
    tukey_result <- result
  }else{
    tukey_result <- rbind(tukey_result, result)
  }

}


write.xlsx(tukey_result, "tukey_result.xlsx")


```

```{r Figure S5c}

library(ggplot2)
library(openxlsx)
library(dplyr)
library(ggsignif)
library(plotrix)
scientific_notation <- function(x) {
  x <- format(x, scientific = TRUE)
  x <- gsub("^(.*)e", "'\\1'e", x)
  x <- gsub("e", "%*%10^", x)
  x <- gsub('\\+', '', x)
  parse(text = x)
}


setwd("D:/2024/01_kanzo_multiomics/qPCR/20250130_GABAR/reslut")


data <- read.xlsx("GABAR_total.xlsx", 1)
data$Cq <- lapply(data$Cq, gsub, pattern="NaN", replacement = "") %>% as.numeric()
nsample <- data %>% distinct(Sample) %>% .$Sample

for (i in c(1:length(nsample))) {
  # i=1
  tarsam <- data %>% filter(Sample == nsample[i]) %>% mutate("Cq_ref" = .$Cq[1]) %>% mutate("dCq" = .$Cq - .$Cq_ref)
  if(i==1){
    dCq <-tarsam
  }else{
    dCq <- rbind(dCq ,tarsam)
  }
}


target <- dCq %>% distinct(Target) %>% .[,1]
nrun <- dCq %>% distinct(Replicate) %>% .[,1]

for (i in c(2:length(target))) {
  # i=2
  # Class <- class[[i]]
  targene <- dCq %>% filter(Target == target[[i]])

  for (j in c(1:length(nrun))) {
    # j=1
    tardf <- targene %>% filter(Replicate == nrun[j]) %>% mutate("ddCq" = .$dCq - .$dCq[1])
    if(j==1){
      ddCq <-tardf
    }else{
      ddCq <- rbind(ddCq ,tardf)
    }
  }
  
  if(i == 2){
    ddCq_df <- ddCq
  }else{
    ddCq_df <- rbind(ddCq_df, ddCq)
  }
}

target


#Plotデータの成形
order <- ddCq_df %>% distinct(Biological.Set.Name) %>% .[,1]
plotdf <- transform(ddCq_df, Biological.Set.Name = factor(Biological.Set.Name, levels = order))
plotdf <- plotdf %>% mutate("2^-ddCt" = 2^(-plotdf$ddCq)) %>% filter(Target != "GAPDH") %>% filter(Biological.Set.Name != "Control")


group <- plotdf %>% distinct(Biological.Set.Name) %>% .$Biological.Set.Name
keep_values <- c("LPS", "GABA+LPS", "LPS+Mus", "LPS+Bac", "agonist+LPS", "GABA+antagonist+LPS")
plotdf <- plotdf %>% filter(Biological.Set.Name %in% keep_values) %>% 
  transform(Biological.Set.Name = factor(Biological.Set.Name, levels = keep_values))

gene <- plotdf %>% distinct(Target) %>% .[,1]

for (i in c(1:length(gene))) {
  # i=2
  tardf <- plotdf %>% filter(Target == gene[[i]])
  
  tukeydf <- TukeyHSD(aov(tardf$`2^-ddCt` ~ tardf$Biological.Set.Name))
  result <- data.frame(Terget = gene[i], pair = rownames(tukeydf$`tardf$Biological.Set.Name`), tukeydf$`tardf$Biological.Set.Name`)
  
  p <- ggplot(tardf, aes(x = Biological.Set.Name, y = `2^-ddCt`, fill = Biological.Set.Name))+ 
    stat_summary(fun = "mean", geom = "bar", width = .8,col = "black", position=position_dodge2(preserve = "single")) +
    stat_summary(fun.max = function(x) mean(x) + std.error(x), 
                 fun.min = function(x) mean(x) - std.error(x), 
                 geom = 'errorbar', width = .1,position = position_dodge(width = 0.8))+
    geom_jitter(size = 2, shape = 21,col = "black", width = 0.15)+
    labs(y = expression(2^{-Delta*Delta*Ct}))+
    scale_fill_manual(values = c("#B71C1C",colorRampPalette(c("#cccbcc","#1f3555"))(length(keep_values)-1)))+
    scale_y_continuous(expand = c(0, 0),limits = c(0, max(tardf$`2^-ddCt`)*1.2), labels = scientific_notation)+ 
    theme(
      axis.text.x = element_text(size = 12, angle = 90, hjust = 1, vjust = 0.5, color= "grey10"),
      axis.text.y = element_text(size = 10, color= "grey10"),
      axis.title.y = element_text(size = 12, color= "grey10"),
      axis.title.x = element_blank(),
      legend.key.size = unit(0.5, 'cm'),
      legend.title = element_blank(),
      plot.title = element_text(hjust=0.5, size = 18, color= "grey10"),
      axis.line = element_line(colour="black"),
      panel.background = element_rect(fill = "transparent", colour = "transparent"),
      aspect.ratio = 0.75)
  
  p
  
  filepath <- paste0("D:/2024/01_kanzo_multiomics/qPCR/20250130_GABAR/reslut/",gene[[i]],"_brep1_vs250327.svg")
  svg(filepath, width = 8, height = 4)
  print(p)
  dev.off()
  
  if(i ==1){
    tukey_result <- result
  }else{
    tukey_result <- rbind(tukey_result, result)
  }
  
}


write.xlsx(tukey_result, "tukey_result_wo_vs250327.xlsx")

```


```{r Figure S5a}

library("ggbreak")

scientific_notation <- function(x) {
  x <- format(x, scientific = TRUE)
  x <- gsub("^(.*)e", "'\\1'e", x)
  x <- gsub("e", "%*%10^", x)
  x <- gsub('\\+', '', x)
  parse(text = x)
}


allcomp <- read.xlsx("D:\\2023\\Zenotof\\NuturalProduct\\20231110_BEHC18_Glycyrrhiza_stdmix_kiuchi_V2\\Result\\Height_0_2023_11_13_20_36_39.xlsx", 5)
levels(as.factor(allcomp$Class))
order <- allcomp %>% distinct(Class) %>% unlist()
allcomp <- transform(allcomp, Class= factor(Class, levels = order))


g <- ggplot(allcomp, aes(x = Class, y = ng))+ 
  stat_summary(fun = "mean", geom = "bar", width = .5, col = "black",fill = "#b9bbca") +
  stat_summary(fun.max = function(x) mean(x) + + std.error(x), 
               fun.min = function(x) mean(x) - + std.error(x), 
               geom = 'errorbar', width = .1)+
  geom_jitter(shape = 21, width = .1, size = 3, fill = "white")+
  labs(y="Concentration (ng / mg)")+
  scale_y_break(breaks = c(300, 2500), scales = 4)+
  scale_y_break(breaks = c(3, 30), scales = 2)+
  scale_y_continuous(expand = c(0, 0), limits = c(0, max(allcomp$ng)*1.1),labels = scientific_notation)+
  theme(
        axis.text = element_text(size = 16, color= "grey10"),
        axis.title.y = element_text(size = 16, color= "grey10", angle = 90, hjust = 0.5 ),
        axis.title.x = element_blank(),
        axis.line = element_line(colour="transparent", size = 1),
        panel.grid = element_blank(),
        axis.text.y.right = element_blank(), axis.ticks.y.right = element_blank(),
        legend.background = element_blank(),
        legend.text = element_text(size = 12),
        plot.title = element_text(hjust=0.5, size = 18, color= "grey10"),
        legend.key.size = unit(0.5, 'cm'),
        panel.background = element_rect(fill = "transparent", colour = "grey10"),
        legend.title = element_blank(),
  )

g


svg("D:\\2023\\Zenotof\\NuturalProduct\\20231124_BEHC18_GABA_std_test_10min\\barplot_20250319.svg", width = 3.5, height = 3.5)
print(g)
dev.off()


```
````
